# Cyclitol secondary metabolism is a central feature of *Burkholderia* leaf symbionts

**DOI:** 10.1101/2022.09.27.509721

**Authors:** Bram Danneels, Monique Blignaut, Guillaume Marti, Simon Sieber, Peter Vandamme, Marion Meyer, Aurélien Carlier

## Abstract

The symbioses between plants of the Rubiaceae and Primulaceae families with *Burkholderia* bacteria represent unique and intimate plant-bacterial relationships. Many of these interactions have been identified through PCR-dependent typing methods, but there is little information available about their functional and ecological roles. We assembled seventeen new endophyte genomes representing endophytes from thirteen plant species, including those of two previously unknown associations. Genomes of leaf endophytes belonging to *Burkholderia s.l*. show extensive signs of genome reduction, albeit to varying degrees. Except for one endophyte, none of the bacterial symbionts could be isolated on standard microbiological media. Despite their taxonomic diversity, all endophyte genomes contained gene clusters linked to the production of specialized metabolites, including genes linked to cyclitol sugar analog metabolism and in one instance non-ribosomal peptide synthesis. These genes and gene clusters are unique within *Burkholderia s.l*. and are likely horizontally acquired. We propose that the acquisition of secondary metabolite gene clusters through horizontal gene transfer is a prerequisite for the evolution of a stable association between these endophytes and their hosts.

## Introduction

Interactions with microbes play an important part in the evolution and ecological success of plants. For example, mycorrhizal associations are present in a vast majority of land plants, and the association with nitrogen-fixing bacteria provided legumes with an important evolutionary advantage (Brundrett, 1991; van Rhijn and Vanderleyden, 1995; Vessey *et al.*, 2005; Smith and Read, 2008). Nevertheless, microbes may also be harmful for plants as microbial pathogen interactions are responsible for major crop losses (Dangl and Jones, 2001; McCann, 2020). Many plant-microbe interactions only occur temporarily: contacts between microbes and the host are often limited to a sub-population or a specific developmental phase of the host. However, in some associations microbes are transferred from parents to offspring in a process called vertical transmission, resulting in permanent associations with high potential for co-evolution (Gundel *et al.*, 2017). While vertically-transmitted microbes are common in the animal kingdom, they have been more rarely described in plants (Fisher *et al.*, 2017).

A particular case of vertically transmitted microbes in plants are the bacterial leaf endophytes found in three different plant families: the monocot Dioscoreaceae, and the dicot Rubiaceae and Primulaceae. In the genera *Psychotria, Pavetta, Sericanthe* (Rubiaceae) and *Ardisia* (Primulaceae) this association may manifest in the form of conspicuous leaf nodules that house extracellular symbiotic bacteria (Miller, 1990; Van Oevelen *et al.*, 2002; Lemaire, Robbrecht, *et al.*, 2011; Lemaire, Van Oevelen, *et al.*, 2012; Ku and Hu, 2014). In some of these systems, the symbiont was detected in seeds, indicating that they can be transmitted vertically (Miller and Donnelly, 1987; Sinnesael *et al.*, 2018). Molecular analysis of the leaf nodules revealed that all endophytes are members of the *Burkholderia sensu lato*, more specifically to the newly defined *Caballeronia* genus (Van Oevelen *et al.*, 2002; Ku and Hu, 2014). Similar leaf endophytes, also belonging to the *Burkholderiaceae*, are present in Rubiaceae species that do not form leaf nodules, including some *Psychotria species* (Lemaire, Lachenaud, *et al.*, 2012; Verstraete *et al.*, 2013). To date, only one symbiont of Rubiaceae and Primulaceae has been cultivated: the endophyte of *Fadogia homblei*, which has been identified as *Paraburkholderia caledonica* (Verstraete *et al.*, 2011). Interestingly, members of *P. caledonica* are also commonly isolated from the rhizosphere or soil and have been detected in leaves of some *Vangueria* species (Verstraete *et al.*, 2014).

Speculations about possible functions of these leaf symbioses have long remained unsubstantiated because efforts to isolate leaf nodule bacteria or to culture bacteria-free plants were unsuccessful (Miller, 1990). Recently, sequencing and assembly of leaf symbiont genomes of several *Psychotria, Pavetta* or *Ardisia* species allowed new hypotheses about the ecological function of leaf symbiosis. Leaf symbiotic *Burkholderia* of *Ardisia crenata*, are responsible for the production of FR900359, a cyclic depsipeptide with potent bioactive and insecticidal properties (Fujioka *et al.*, 1988; Carlier *et al.*, 2016). Similarly, analysis of the genome of *Ca*. Burkholderia kirkii (*Ca*. B. kirkii), the leaf symbiont of *Psychotria kirkii*, revealed a prominent role of secondary metabolism (Carlier and Eberl, 2012). In this species, two biosynthetic gene clusters harboured on a plasmid encode two homologs of a 2-*epi*-5-*epi*-valiolone synthase (EEVS). EEVS are generally required for the production of cyclitol sugar analogs, a family of bioactive natural products with diverse targets (Mahmud, 2003, 2009). *Ca*. B. kirkii is likely involved in the synthesis of two cyclitol metabolites: kirkamide, a C_7_N aminocyclitol with insecticidal properties, and streptol glucoside, a derivative of valienol with broad allelopathic activities (Sieber *et al.*, 2015; Georgiou *et al.*, 2021). Similar gene clusters containing putative EEVS were also detected in the genomes of other *Psychotria* and a *Pavetta* leaf symbionts (Pinto-Carbó *et al.*, 2016), further highlighting the importance of cyclitol compounds in these leaf symbioses.

C_7_ cyclitols are a group of natural products derived from the pentose phosphate pathway intermediate sedoheptulose-7-phosphate (SH7P) (Mahmud, 2003). Proteins of the sugar phosphate cyclase family are key enzymes in the synthesis of C_7_ cyclitols. Enzymes of this family catalyse the cyclization of sugar compounds, an important step in primary and secondary metabolism (Wu *et al.*, 2007). Within this family, three main categories of enzymes use SH7P as a substrate: desmethyl-4-deoxygadusol synthase (DDGS), 2-*epi*-valiolone synthase (EVS) and 2-*epi*-5-*epi*-valiolone synthase (EEVS), of which EEVS is the only known enzyme involved in C_7_N aminocyclitol synthesis (Osborn *et al.*, 2017). EEVS were originally only found in bacteria, where they catalyse the first step in the biosynthesis of C_7_N aminocyclitol secondary metabolites (Mahmud, 2003; Sieber *et al.*, 2015). More recently, EEVS homologs have been discovered in some Eukaryotes such as fish, reptiles, and birds as well (Osborn *et al.*, 2015, 2017).

A second common feature of the leaf endophytes in Rubiaceae and Primulaceae is their reduced genomes. Leaf nodule *Burkholderia* symbionts of Rubiaceae and Primulaceae typically have smaller genomes than free-living relatives, as well as a lower coding capacity (Pinto-Carbó *et al.*, 2016). This reductive genome evolution is thought to be a result of increased genetic drift sustained in bacteria that are strictly host-associated, which leads to fixation of deleterious and/or neutral mutations and eventually to the loss of genes (Pettersson and Berg, 2007). This process is best documented in obligate insect symbionts such as *Buchnera* and *Serratia*, endosymbionts of aphids, or in *Sodalis*-allied symbionts of several insect groups (Shigenobu *et al.*, 2000; Toh *et al.*, 2006; Manzano-Marín *et al.*, 2018). Some of these symbionts have extremely small genomes and may present an extensive nucleotide bias towards adenosine and thymine (AT-bias) (Moran *et al.*, 2008). The process of genome reduction has multiple stages: first, recently host-restricted symbionts begin accumulating pseudogenes and insertion elements (McCutcheon and Moran, 2012; Lo *et al.*, 2016; Manzano-Marín and Latorre, 2016). Non-coding and selfish elements eventually get purged from the genomes over subsequent generations, which together with the general deletional bias in bacteria results in a decrease in genome size (Mira *et al.*, 2001). This ultimately leads to symbionts with tiny genomes, with only a handful of essential genes necessary for survival or performing their role in the symbiosis. This process has been well documented in the leaf nodule symbionts of *Psychotria, Pavetta* and *Ardisia* species, but little is known about the genomes and functions of endophytes in species that do not form leaf nodules, notably Rubiaceae species of the *Vangueria* and *Fadogia* genera.

Here, we performed a comparative study of Rubiaceae and Primulaceae leaf endophytes from leaf nodulating and non-nodulating plant species using genomes assembled from shotgun metagenome sequencing data as well as isolates. We constructed a dataset of 26 leaf symbiont genomes (of which 17 new genomes from this study) from 22 plant species in 5 genera. All leaf symbionts show signs of genome reduction, in varying degree, and horizontal acquisition of secondary metabolite clusters is a universal phenomenon in these bacteria.

## Material and Methods

### Sample collection and DNA extraction

Leaves of Rubiaceae and Primulaceae species were freshly collected from different locations in South Africa or requested from the living collection of botanical gardens (Table S1). Attempts to isolate the endophytes were made for all fresh samples collected in South Africa (Table S1). Leaf tissue was surface sterilized using 70% ethanol, followed by manual grinding of the tissue in 0.4% NaCl. Supernatants were plated on 10% tryptic soy agar medium (TSA, Sigma) and R2A medium (Oxoid) and incubated at room temperature for 3 days or longer until colonies appeared. Single colonies were picked and passaged twice on TSA medium. Isolates were identified by PCR and partial sequencing of the 16S rRNA gene using the pA/pH primer pair (5’-AGAGTTTGATCCTGGCTCAG and 5’-AAGGAGGTGATCCAGCCGCA) (Edwards *et al.*, 1989). PCR products were sequenced using the Sanger method at Eurofins Genomics (Ebersberg, Germany).

DNA was extracted from whole leaf samples as follows. Whole leaves were ground in liquid nitrogen using a mortar and pestle. Total DNA was extracted using the protocol of Inglis *et al.* (Inglis *et al.*, 2018). Total DNA from a *Fadogia homblei* isolate was extracted following Wilson (Wilson, 2001). Sequencing library preparation and 2×150 paired-end metagenome sequencing was performed by the Oxford Wellcome Centre for Human Genetics or by Novogene Europe (Cambridge, UK) using the Illumina NovaSeq 6000. Sequencing reads were classified using Kraken v2.1.2 against a custom database comprising complete prokaryotic and plastid genome sequences deposited NCBI RefSeq (accessed 4/4/2021), and visualised using KronaTools v2.7.1 (Ondov *et al.*, 2011; Wood *et al.*, 2019).

### Bacterial genome assembly

Sequencing reads were trimmed and filtered using fastp v0.21.0 with default settings (Chen *et al.*, 2018). Overlapping paired-end reads were merged using NGmerge with default settings (Gaspar, 2018). For sequencing reads derived from new leaf samples, metagenome assemblies were created using metaSPAdes v3.15 on default settings but including the merged reads (Nurk *et al.*, 2017). Metagenomes were binned using Autometa, using a minimal contig length of 500 bp, taxonomy filtering (-m) and maximum-likelihood recruitment (using the -r option)(Miller *et al.*, 2019). Genome bins identified as *Caballeronia, Paraburkholderia*, or *Burkholderia* by Autometa were further assembled by mapping the original reads to these bins using smalt v0.7.6 (Ponsting and Ning, 2010). Mapped reads were extracted using samtools v1.9 (Li *et al.*, 2009) and reassembled using SPAdes v3.15 (Bankevich *et al.*, 2012) in default settings but using the --careful option, and binned again using Autometa. Contigs likely derived from eukaryotic contamination were removed after identification by blastn searches (e-value < 1e^−6^) against the NCBI nucleotide database (accessed January 2021) (Camacho *et al.*, 2009). Per-contig coverage information was calculated using samtools and contigs with less than 10% or more than 500% of the average coverage were manually investigated, and sequences likely derived from other bacterial or eukaryotic genomes were removed. Genome assembly for reads derived from isolates were assembled using Skesa v2.4.0 using default settings (Souvorov *et al.*, 2018). Assembly statistics were compiled using Quast v5.1.0 (Gurevich *et al.*, 2013).

To provide a more homogenous dataset for comparative genomics, Illumina read data for six previously published Rubiaceae symbionts, and the symbionts of *Ardisia crenata* and *Fadogia homblei* were re-assembled as above but using the published draft genomes as trusted contigs for both metaSPAdes and SPAdes assemblies (Table S2). The resulting assemblies were compared to the published assemblies using dotplots created by MUMmer (Marçais *et al.*, 2018). Genome assemblies of the symbionts of *Psychotria kirkii* (Carlier and Eberl, 2012; Carlier *et al.*, 2013) and *Psychotria punctata* (Pinto-Carbó *et al.*, 2016) were downloaded from Genbank (Table S2). To assess whether the (re-)assembled genomes represent new species, genomes were analysed using TYGS (Type Strain Genome Server) (Meier-Kolthoff and Göker, 2019), and NCBI Blastn-based Average Nucleotide Identities (ANI) values calculated using the JSpecies web server (Richter *et al.*, 2016) and the pyANI python package (https://github.com/widdowquinn/pyani).

### Genome annotation and pseudogene prediction

Assembled genomes were annotated using the online RASTtk pipeline (Brettin *et al.*, 2015), using GenemarkS as gene predictor, and locus tags were added using the Artemis software v18.1.0 (Carver *et al.*, 2012). Prediction of pseudogenes was performed using an updated version of the pseudogene prediction pipeline previously used for leaf symbionts (Carlier *et al.*, 2013). Briefly, orthologs of predicted proteins sequences of each genome in a dataset of published *Burkholderia* genomes (Table S3) were determined using Orthofinder v2.5.2 (Emms and Kelly, 2019) with default settings. The nucleotide sequences of each gene, including 200bp flanking regions, were aligned to the highest scoring sequence in each orthogroup using TFASTY v3.6 (Pearson, 2000). Genes were considered as pseudogenes if the alignment spanned over 50% of the query protein and the query protein contained a frameshift, or a nonsense mutation resulting in an uninterrupted alignment shorter than 80% of the target sequence. Moreover, ORFs were classified as non-functional if at least one of the following criteria was true: amino acid sequence shorter than 50 residues which did not cluster in an orthogroup, and sequence without any significant blastx hit against the reference database (e-value cut off = 0.001); proteins without predicted orthologs in the *Burkholderia* dataset, but which showed a blastx hit against the reference set in an alternative reading frame; and finally proteins without any hit in the *Burkholderia* genome database or in the NCBI nr database. Blastx and blastp searches were performed using DIAMOND v2 (Buchfink *et al.*, 2021). For the genomes of the symbionts of *P. kirkii* and *P. punctata* the original gene and pseudogene predictions were used. Insertion elements in both newly assembled and re-assembled genomes were predicted using ISEscan v1.7.2.3 with default settings (Xie and Tang, 2017).

### Phylogenetic analysis

16S rRNA sequences were extracted from the endophyte (meta)genomes using Barrnap v0.9 (https://github.com/tseemann/barrnap). For genomes where no complete 16S rRNA could be detected, reads were mapped to the 16S rRNA gene of the closest relative with a complete 16S rRNA sequence. These reads were assembled using default SPAdes (Prjibelski *et al.*, 2020) using the --careful option. Near complete (>95%) 16S rRNA sequences could be extracted using these methods, except for the hypothetical endophyte of *Pavetta revoluta*. The 16S rRNA sequences were identified using the EzBiocloud 16S rRNA identification service (https://www.ezbiocloud.net/identify). Phylogenetic analysis of the leaf endophytes and *Burkholderia s.l*. genomes was performed using the UBCG pipeline v3.0 (Na *et al.*, 2018). The pipeline was run using the default settings, except for the gap-cutoff (-f 80). The resulting superalignment of 92 core genes was used for maximum-likelihood phylogenetic analysis using RAxML, using the GTRGAMMA evolution model, and performing 100 bootstrap replications (Stamatakis, 2014). Plastid reference alignments were created using Realphy v1.12 using standard settings and the *Coffea arabica* chloroplast genome (NCBI accession NC_008535.1) as reference (Bertels *et al.*, 2014). Published chloroplast genomes of *Ardisia mamillata* (NCBI accession MN136062), *Psychotria kirkii* (NCBI accession KY378696), *Pavetta abyssinica* (NCBI accession KY378673), *Pavetta schumanniana* (NCBI Accession MN851271), and *Vangueria infausta* (NCBI accession MN851269) were also included in the alignment. Phylogenetic trees were constructed using PhyML v3.3.3 with automatic model selection, and 1000 bootstrap replicates (Guindon *et al.*, 2010). For plant species with uncertain taxonomic identification, seven plant markers were extracted by blastn searches against the metagenome: ITS, nad4, rbcL and rpl16 of *Pavetta abyssinica* (NCBI accessions MK607930.1, KY492180.1, Z68863.1, and KY378673.1), matK from *Pavetta indica* (NCBI accession KJ815920.1), petD from *Pavetta bidentata* (NCBI accession JN054223.1), and trnTF from *Pavetta sansibarica* (NCBI accession KM592134.1).

Core-genome phylogenies of symbiont genomes were constructed by individually aligning the protein sequences of all single-copy core genes using MUSCLE, back-translating to their nucleotide sequence using T-Coffee v13.45 (Di Tommaso *et al.*, 2011), and concatenating into one superalignment. Maximum-likelihood phylogenetic analysis was performed using RAxML, using the GTRGAMMA evolution model, 100 bootstrap replicates, and using partitioning to allow the model parameters to differ between genes. Phylogenetic trees were visualised and edited using iTOL (Letunic and Bork, 2019).

### Comparative genomics

Ortholog prediction between leaf symbiont genomes and a selection of reference genomes of the *Burkholderia, Paraburkholderia* and *Caballeronia* genera (BPC-set; selected using NCBI datasets tool (https://www.ncbi.nlm.nih.gov/datasets/genomes); Table S3) was performed using Orthofinder v2.5.2 using default settings (Emms and Kelly, 2019). Core genome overlap was visualised in Venn diagrams using InteractiVenn (Heberle *et al.*, 2015). Non-essential core genes were identified by blastp searches against the database of essential genes (DEG)(Zhang, 2004), identifying as putative essential genesORFs with significant matches in the database (e-value < 1e^−6^). Standardised functional annotation was performed using eggNOG-mapper v2.1.2 (Huerta-Cepas *et al.*, 2019; Cantalapiedra *et al.*, 2021). Enrichment of protein families in leaf symbiont genomes was determined by comparing the proportion of members of leaf symbionts and the BPC-set in orthogroups. Enriched KEGG pathways were identified by comparing the average per-genome counts of genes in every pathway between leaf symbiont genomes and genomes from the BPC-set. Presence of motility and secretion system clusters was investigated using the TXSScan models implemented in MacSyFinder (Abby *et al.*, 2014, 2016). Homologues of the *Ca*. B. kirkii putative 2-*epi*-5-*epi*-valiolone synthase (EEVS) were identified by blastp searches against the proteomes of the leaf symbiont genomes (e-value cut-off: 1e^−6^). Putative EEVS genes were searched against the SwissProt database, and functional assignment was done by transferring the information from the closest match within the sugar phosphate cyclase superfamily (Schneider *et al.*, 2004; Osborn *et al.*, 2017). Contigs containing these genes were identified and extracted using Artemis, and aligned using Mauve (Lòpez-Fernàndez *et al.*, 2015). Gene phylogenies were constructed by creating protein alignments using MUSCLE followed by phylogenetic tree construction using FastTree (Price *et al.*, 2009), including the protein sequences of three closely related proteins in other species, determined by blastp searches against the RefSeq protein database (accessed July 2021).

The data generated in this study have been deposited in the European Nucleotide Archive (ENA) at EMBL-EBI under accession number PRJEB52430 (https://www.ebi.ac.uk/ena.browser/view/PREJB52430).

## Results

### Detection and identification of leaf endophytes

To gain insight into potential association of various Primulaceae and Rubiaceae species with *Burkholderia s.l*. endosymbionts, we collected samples from 16 Rubiaceae (1 *Fadogia* sp., 5 *Pavetta* spp., 2 *Psychotria* spp., and 8 *Vangueria* spp.) and 3 Primulaceae (3 *Ardisia* spp.) species (Table S1). We extracted DNA from entire leaves and submitted the samples to shotgun sequencing without pre-processing of the samples to remove host or organellar DNA. We found evidence for endophytic *Burkholderia* in 14 out of 19 species investigated (Table S1). In these samples, the proportion of sequencing reads identified as *Burkholderiaceae* ranged from 5% to 57% of the total, except for the *Pavetta revoluta* sample (0.4%) and 1 of 2 *Vangueria infausta* samples (0.9%). Analysis of 16S rRNA sequences revealed 100% pairwise identity over 1529 bp suggesting that the same endophyte species was present in both *V. infausta* samples. In *Pavetta revoluta*, the closest relative of the leaf endophyte based on 16S rRNA sequence similarity was *Caballeronia calidae* (98.89% identity over 808 bp; Table S1). Of the nine species with significant amounts of *Burkholderia s.l*. reads and for which isolation attempts were made (Table S1), only the endophyte of *Fadogia homblei* could be cultured (isolate R-82532). Leaf samples of four species (*Psychotria capensis, Psychotria zombamontana, Pavetta ternifolia*, and *Pavetta capensis*) contained low amounts of bacterial DNA (<2% of reads), and likely do not have stable symbiotic endophyte associations. Seven percent of the reads obtained from the *Pavetta indica* sample were classified as bacterial, but with a diverse range of taxa present indicating possible contamination with surface bacteria (Figure S1). Plastid phylogenies indicated that samples attributed to *Pavetta capensis* and *Pavetta indica* did not cluster with other *Pavetta* species (Figure S2). Analysis of genetic markers revealed that our *Pavetta indica* sample was likely a misidentified *Ixora* sp. Analysis of *Pavetta capensis* marker genes revealed the specimen is likely part of the Apocynaceae plant family, with a 100% identity match against the *rbcL* sequence of *Pleiocarpa mutica*. These samples were not taken into account in further analyses.

Analysis of the 16S rRNA sequences extracted from metagenome-assembled genomes (MAGs) identified all leaf endophytes as *Burkholderia s.l*. (Table S1). Phylogenetic analysis shows that all endophytes of *Psychotria, Pavetta*, and *Ardisia* cluster within the genus *Caballeronia*, while the endophytes of *Vangueria* and *Fadogia* belong to the *Paraburkholderia* genus (Figure 1A). All endophytes of *Ardisia* are closely related to each other and form a clade with *Caballeronia udeis* and *Caballeronia sordidicola*. Based on the commonly used ANI (95-96%) cut-off, these endophytes are separate species from *C. udeis* and *C. sordidicola* (ANI <94%; 16S rRNA sequence identity <98.4). The endophytes of *Ardisia crenata* and *Ardisia virens* are very closely related and belong to the same species: *Ca*. Burkholderia crenata (ANI >99%; 16S rRNA sequence identity 99.8%) (Table S4). Similarly, the endophytes of *Ardisia cornudentata* and *Ardisia mamillata* belong to the same species (ANI = 95.56%), which we tentatively named *Ca*. Burkholderia ardisicola (species epithet from *Ardisia*, the genus of the host species, and the Latin suffix - *cola* (from L. n. *incola*), dweller, see species description in Supplementary Information). Endophytes of *Psychotria* and *Pavetta* are scattered across the *Caballeronia* phylogeny, but all are taxonomically distinct from free-living species (Figure 1A; ANI <93% with closest non-endophyte relatives). Each of these endophytes also represents a distinct bacterial species with pairwise Average Nucleotide Identity (ANI) values below the commonly accepted species threshold of 95-96%, except for *Ca*. P. schumanniana and *Ca*. B. kirkii whose genomes share 95.65% ANI (Table S4). The endophytes of *Vangueria* and *Fadogia* form three distinct lineages of *Paraburkholderia*. The endophytes of *Vangueria dryadum* and *Vangueria macrocalyx* are nearly identical (ANI >99.9%; identical 16S rRNA), but do not belong to any known *Paraburkholderia* species (ANI <83% with closest relative *Paraburkholderia* species). We tentatively assigned these bacteria to a new species which we named *Ca. Paraburkholderia dryadicola* (from a Dryad, borrowed from the species epithet of one of the host species, and Latin suffix – *cola*, see species description in Supplementary Information). Similarly, the endophytes of *V. infausta, V. esculenta, V. madagascariensis, V. randii*, and *V. soutpansbergensis* cluster together with *Paraburkholderia phenoliruptrix* (Figure 1A). While the endophyte of *Vangueria soutpansbergensis* forms a separate species (named here *Ca. Paraburkholderia soutpansbergensis*; ANI <95% with *P. phenoliruptrix*) the other endophytes fall within the species boundaries of *P. phenoliruptrix*. (ANI 95-96% between these endophytes and *P. phenoliruptrix*). Lastly, the endophytes of *Fadogia homblei* and *Vangueria pygmaea* showed identical 16S rRNA sequences, and clustered with *Paraburkholderia caledonica, P. strydomiana*, and *P. dilworthii* (Figure 1A). Similarly high ANI values (>97.5%) and 16S rRNA sequence similarity (>99.7%) ambiguously fall within the species boundaries of both *P. caledonica* and *P. strydomiana*. Because endophytes of *F. homblei* were previously classified as *P. caledonica* (Verstraete *et al.*, 2011, 2014), we propose classifying the endophytes of *F. homblei* and *V. pygmaea* as members of *P. caledonica*, and consider *P. strydomiana* a later heterotypic synonym of *P. caledonica*.

**Figure 1:**
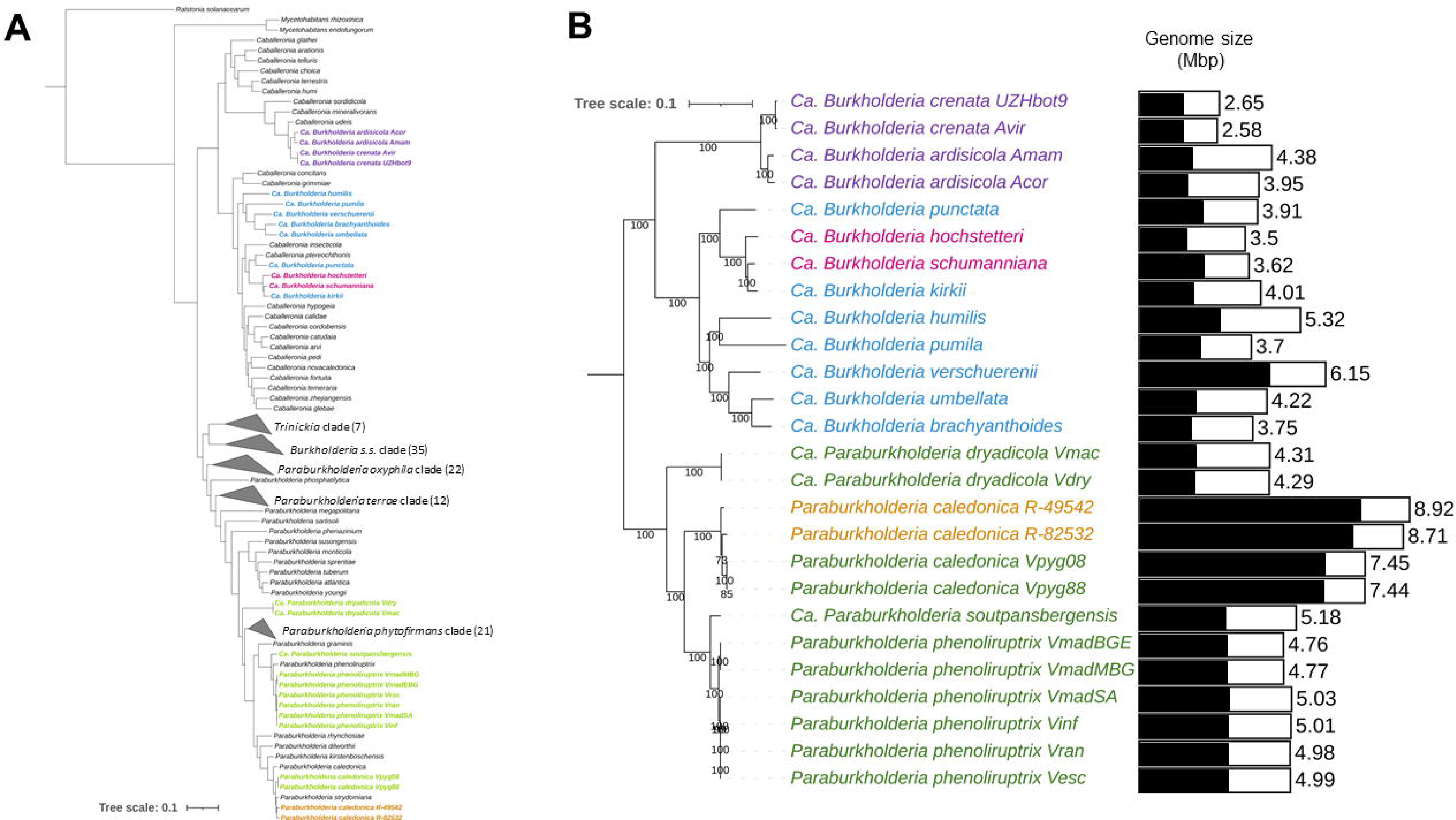
Phylogeny of *Burkholderia, Caballeronia*, and *Paraburkholderia*, including the leaf endophytes. **(A)** UBCG phylogeny of the *Burkholderia s.l*. based on 92 conserved genes. Bootstrap support values based on 100 replications are displayed on the branches. Branches with <50% support were collapsed. *Ralstonia solanacearum* was used as outgroup to root the tree. Coloured samples in boldface represent the leaf endophytes from Rubiaceae and Primulaceae **(B)** Core genome phylogeny of leaf endophytes based on alignment of 423 single-copy core genes. Bootstrap support values based on 100 replicates are shown on the branches. Samples are colour-coded based on the host genus: Purple – *Ardisia*; Blue – *Psychotria*; Pink – *Pavetta*; Green – *Vangueria*; Orange – *Fadogia*; Black bars represent the coding capacity of the genome (the proportion of the genome coding for functional proteins).

Phylogenetic analysis based on the core genomes of endophytes indicates a general lack of congruence with the host plant phylogeny (Figure S3). Endophytes of *Ardisia* are monophyletic within the *Caballeronia* genus and follow the host phylogeny. In contrast, endophytes of *Pavetta* are not monophyletic and are nested within the *Psychotria* endophytes. Similarly, the *Fadogia homblei* endophyte clusters with endophytes of *Vangueria*.

### Leaf endophyte genomes show signs of genome reduction

We could assemble nearly complete bacterial genomes for all samples where we detected *Burkholderia* endophytes, except for those of the *Pavetta revoluta* and one *Vangueria infausta* sample with too few bacterial reads. Binning analysis grouped endophyte sequences in a single bin per sample, with high completeness (>95%) and purity (>97%). Most assemblies ranged between 3.5 and 5 Mbp in size, with 2 outliers: 2.58 Mbp for Ca. B. crenata Avir, and 8.92 Mbp for *P. caledonica* R-49542 (Table 1). The %G+C of all genomes fell in the range of 59-64 %G+C, which is within the range of free-living *Paraburkholderia* and *Caballeronia* genomes (Vandamme *et al.*, 2017). All genomes showed signs of ongoing genome reduction. Because of rampant null or frameshift mutations, a large proportion of predicted CDS code for non-functional proteins. As a result, coding capacity is low for all endophyte genomes varying between 83% in *P. caledonica* R-49542 and 40% in *Ca*. B. ardisicola Acor (Figure 1B, Table 1). In addition, insertion sequence (IS) elements make up a large amount of the genomes: 1.97% of the assembly size on average, but up to almost 10% in some symbionts of *Psychotria* (Table 1). Reassembly of previously investigated endophytes of *Psychotria* and *Pavetta* yielded genomes of similar size to the original assemblies, except for Ca. *Burkholderia schumanniana*. The original genome assembly size was estimated at 2.4 Mbp, while our reassembly counted 3.62 Mbp. A dot plot between both assemblies indicated that the size discrepancy is not solely due to differential resolution of repeated elements (Figure S4). Thus, our new assembly includes 1.2 Mbp of genome sequence that was missed in the original assembly.

**Table 1:** Genome statistics of newly assembled and re-assembled leaf endophyte genomes. Coding capacity refers to the proportion of the genome that codes for functional proteins.

| Endophyte | Host species | Type | Assembly size (Mb) | Contigs | N50 (bp) | %G+C | Annotated genes | Functional genes | Pseudo genes | Coding Capacity (%) | IS elements | IS total length (bp) | Proportion IS (%) |
| --- | --- | --- | --- | --- | --- | --- | --- | --- | --- | --- | --- | --- | --- |
| *Ca. Burkholderia ardisicola* Acor | *Ardisia cornudentata* | New assembly | 3,95 | 332 | 19528 | 59,23 | 6975 | 2026 | 4949 | 40,30 | 35 | 32834 | 0,83 |
| *Ca. Burkholderia crenata* UZHbot9 | *Ardisia crenata* | Re-assembly | 2,65 | 607 | 6399 | 59,02 | 3982 | 1670 | 2312 | 54,73 | 96 | 69678 | 2,63 |
| *Ca. Burkholderia ardisicola* Amam | *Ardisia mamillata* | New assembly | 4,38 | 333 | 19687 | 59,47 | 7472 | 2297 | 5175 | 40,95 | 59 | 48385 | 1,10 |
| *Ca. Burkholderia crenata* Avir | *Ardisia virens* | New assembly | 2,58 | 605 | 6517 | 59,05 | 3839 | 1648 | 2191 | 56,13 | 63 | 42190 | 1,64 |
| *Paraburkholderia caledonica* R-49542 | *Fadogia homblei* | New assembly | 8,92 | 148 | 145314 | 61,59 | 9185 | 7695 | 1490 | 82,83 | 96 | 117852 | 1,32 |
| *Paraburkholderia caledonica* R-82532 | *Fadogia homblei* | New assembly | 8,71 | 123 | 239289 | 61,53 | 9054 | 7353 | 1701 | 81,03 | 77 | 91097 | 1,05 |
| *Ca. Burkholderia hochstetteri* | *Pavetta hochstetteri* | New assembly | 3,50 | 324 | 18152 | 62,51 | 5453 | 1823 | 3630 | 44,53 | 29 | 38062 | 1,09 |
| *Ca. Burkholderia schumanniana* | *Pavetta schumanniana* | Re-assembly | 3,62 | 412 | 14848 | 63,47 | 4938 | 2453 | 2485 | 59,95 | 69 | 44239 | 1,22 |
| *Ca. Burkholderia brachyanthoides* | *Psychotria brachyanthoides* | Re-assembly | 3,75 | 648 | 8356 | 61,00 | 6284 | 2109 | 4175 | 46,54 | 223 | 149135 | 3,98 |
| *Ca. Burkholderia humilis* | *Psychotria humilis* | Re-assembly | 5,32 | 238 | 103328 | 59,60 | 7828 | 3264 | 4564 | 50,04 | 64 | 63278 | 1,19 |
| *Ca. Burkholderia kirkii* | *Psychotria kirkii* | Reference | 4,01 | 203 | 44916 | 62,91 | 6329 | 2069 | 4260 | 45,80 | 375 | 353298 | 8,81 |
| *Ca. Burkholderia pumila* | *Psychotria pumila* | Re-assembly | 3,70 | 463 | 12628 | 59,13 | 6835 | 2192 | 4643 | 45,41 | 195 | 153499 | 4,15 |
| *Ca. Burkholderia punctata* | *Psychotria punctata* | Reference | 3,91 | 48 | 100248 | 64,00 | 4864 | 2539 | 2325 | 54,61 | 310 | 358729 | 9,17 |
| *Ca. Burkholderia umbellata* | *Psychotria umbellata* | Re-assembly | 4,22 | 333 | 28025 | 61,30 | 6967 | 2306 | 4661 | 44,37 | 91 | 68761 | 1,63 |
| *Ca. Burkholderia verschuerenii* | *Psychotria verschuerenii* | Re-assembly | 6,15 | 401 | 27267 | 62,07 | 7440 | 4839 | 2601 | 70,21 | 88 | 60714 | 0,99 |
| *Ca. Paraburkholderia dryadicola* Vdry | *Vangueria dryadum* | New assembly | 4,29 | 153 | 50748 | 61,26 | 7076 | 2229 | 4847 | 43,21 | 38 | 35272 | 0,82 |
| *Paraburkholderia phenoliruptrix* Vesc | *Vangueria esculenta* | New assembly | 4,99 | 180 | 50333 | 63,54 | 6347 | 3329 | 3018 | 59,78 | 46 | 54597 | 1,09 |
| *Paraburkholderia phenoliruptrix* Vinf | *Vangueria infausta* | New assembly | 5,00 | 181 | 49920 | 63,51 | 6377 | 3320 | 3057 | 59,29 | 50 | 58387 | 1,17 |
| *Ca. Paraburkholderia dryadicola* Vmac | *Vangueria macrocalyx* | New assembly | 4,31 | 150 | 54987 | 61,30 | 7111 | 2243 | 4868 | 43,06 | 40 | 37709 | 0,87 |
| ***Paraburkholderia phenoliruptrix***<br>**VmadMBG** | *Vangueria*<br>*madagascariensis* | New assembly | 4,77 | 247 | 34361 | 63,48 | 5912 | 3214 | 2698 | 61,09 | 45 | 55093 | 1,15 |
| ***Paraburkholderia phenoliruptrix***<br>**VmadEBG** | *Vangueria*<br>*madagascariensis* | New assembly | 4,76 | 242 | 34985 | 63,48 | 5901 | 3212 | 2689 | 60,97 | 45 | 53173 | 1,12 |
| ***Paraburkholderia phenoliruptrix***<br>**VmadSA** | *Vangueria*<br>*madagascariensis* | New assembly | 5,03 | 194 | 50250 | 63,49 | 6444 | 3291 | 3153 | 59,22 | 47 | 48936 | 0,97 |
| ***Paraburkholderia caledonica***<br>**Vpyg88** | *Vangueria pygmaea* | New assembly | 7,44 | 92 | 232014 | 61,89 | 7426 | 6194 | 1232 | 82,23 | 54 | 74083 | 1,00 |
| ***Paraburkholderia caledonica***<br>**Vpyg08** | *Vangueria pygmaea* | New assembly | 7,45 | 106 | 232088 | 61,90 | 7449 | 6193 | 1256 | 82,33 | 60 | 79510 | 1,07 |
| ***Paraburkholderia phenoliruptrix***<br>**Vran** | *Vangueria randii* | New assembly | 4,98 | 205 | 50270 | 63,33 | 6379 | 3294 | 3085 | 59,47 | 59 | 73129 | 1,47 |
| ***Ca. Paraburkholderia***<br>**soutpansbergensis** | *Vangueria*<br>*soutpansbergensis* | New assembly | 5,18 | 51 | 337347 | 63,12 | 6801 | 3259 | 3542 | 55,24 | 35 | 44578 | 0,86 |

*Burkholderia* leaf endophytes in Rubiaceae and Primulaceae shared a core genome of 607 genes (Figure S5). Even within specific phylogenetic lineages the core genomes were small: 774 genes in endophytes belonging to the *Caballeronia* symbionts of *Psychotria* and *Pavetta*, 1001 genes in endophytes of *Caballeronia* symbionts of *Ardisia*, and 1199 in *Paraburkholderia* endophytes of *Fadogia* and *Vangueria*. This corresponds to 29.5%, 52.4%, and 28.4% of the average functional proteome for each species cluster, respectively. Only 28 proteins of the total core genome did not show significant similarity with proteins from the database of essential genes (Table S5). Eleven of these proteins have unknown functions and a five are membrane-related. Fifteen genes of the endophyte core genome did not have orthologs in >95% of related *Burkholderia, Caballeronia*, and *Paraburkholderia* genomes (Table S6). No COG category was specifically enriched in this set of proteins.

Because secretion of protein effectors is often a feature of endophytic bacteria (Brader *et al.*, 2017), we searched for genes encoding various secretion machineries in the genomes of *Burkholderia* endophytes. Flagellar genes, as well as Type III, IV or VI secretion system were not conserved in all leaf endophytes (Figure S6). The most eroded symbionts of *Psychotria, Pavetta*, and *Ardisia* lack almost all types of secretion systems, and most also lack a functional flagellar apparatus. Type V secretion systems are present in *Ca*. Burkholderia ardisicola Acor, *Ca*. B. pumila, and *Ca*. B. humilis. The genomes of *Paraburkholderia* symbionts of *Vangueria* and *Fadogia* were generally richer in secretions systems, but only T1SS and T2SS are conserved. A Type V secretion system is present in all *Paraburkholderia* endophytes except *Ca*. Paraburkholderia dryadicola. The flagellar apparatus is missing in *Ca*. P. dryadicola, *Ca*. P. soutpansbergensis, and *P. phenoliruptrix* Vesc, and is incomplete in some other *P. phenoliruptrix* endophytes. Lastly, only the genomes of *Paraburkholderia caledonica* endophytes encode a complete set of core Type VI secretion system proteins.

### Genes related to secondary metabolism are enriched in leaf endophytes

We wondered if specific metabolic pathways might be enriched in genomes of leaf symbionts, despite rampant reductive evolution. We assigned KEGG pathway membership for each predicted functional CDS (thus excluding predicted pseudogenes) in leaf symbiont genomes as well as a set of free-living representative *Paraburkholderia* or *Caballeronia* species. The number of genes assigned to a majority of the KEGG pathways (256 pathways in total) was significantly smaller in endophyte genomes compared to their free-living relatives.

A small portion (86 pathways) did not differ between leaf symbionts and free-living representatives. Genes belonging to a single pathway were significantly enriched in leaf endophytes: acarbose and validamycin biosynthesis (KEGG pathway map00525). Acarbose and validamycin are aminocyclitols synthesized via 2-*epi*-5-*epi*-valiolone synthase (EEVS). EEVS catalyses the first committed step of C_7_N aminocyclitol synthesis^23,24^, and likely plays a role in the production of kirkamide, a natural C_7_N aminocyclitol present in leaves of *Psychotria kirkii* and other nodulated Rubiaceae, as well as streptol and streptol glucoside, 2 cyclitols with herbicidal activities (Pinto-Carbó *et al.*, 2016). Indeed, of 10 *Ca*. Burkholderia kirkii genes assigned to KEGG pathway map00525, 8 genes were previously hypothesised to play a direct role in the synthesis of C_7_N aminocyclitol or derived compounds (Pinto-Carbó *et al.*, 2016). Similarly, 7 out of 11 orthogroups most enriched in leaf endophytes were linked to cyclitol synthesis (Table S7). To gain a better understanding of the distribution of cyclitol biosynthetic clusters in leaf endophytes, we searched for homologs of the two 2-*epi*-5-*epi*-valiolone synthase (EEVS) genes of *Ca*. Burkholderia kirkii (locus tags BKIR_C149_4878 and BKIR_C48_3593) in the other leaf endophyte genomes. We detected putative EEVS homologs in all but the two genomes of *Ca*. B. crenata. For *Ca*. B. crenata UZHbot9 we have previously shown the genome encodes a non-ribosomal peptide synthase likely responsible for the synthesis of the depsipeptide FR900359 (Fujioka *et al.*, 1988; Carlier *et al.*, 2016; Crüsemann *et al.*, 2018), and these genes were also detected in *Ca*. B. crenata Avir. Because EEVSs are phylogenetically related to 3-dehydroquinate synthases (DHQS), we aligned the putative EEVS sequences retrieved from leaf endophytes to EEVS and DHQS sequences in the Swissprot database. All putative EEVS sequences retrieved from leaf endophytic *Burkholderia* were phylogenetically related to *bona fide* EEVS proteins, but not to dehydroquinate synthase (DHQS) and other sedoheptulose 7-phosphate cyclases. EEVS are otherwise rare in *Burkholderia* s. l., with putative EEVSs present in only 11 out of 5674 publicly available *Burkhoderiaceae* genomes (excluding leaf symbiotic bacteria) in the NCBI RefSeq database as of June 2022 (Figure S7).

### Evolution of cyclitol metabolism in leaf endophytic Burkholderia

Phylogenetic analysis of the endophyte EEVS protein sequences showed the presence of two main clades of *Burkholderia* EEVS homologs, as well as a divergent homolog in the genome of *Ca*. B. ardisicola Acor, and a second divergent homolog in *Ca*. P. dryadicola (Figure 2A). The gene context of these EEVS genes in the different clades reveals that the two main EEVS clades correspond to the two conserved gene clusters previously hypothesized to play a role in kirkamide and streptol glucoside biosynthesis in *Ca*. Burkholderia kirkii (Carlier *et al.*, 2013). The gene order of these clusters is very similar in every genome, with a similar genomic context in closely related genomes (Table 2–3). These gene clusters are generally flanked by multiple mobile elements, consistent with acquisition via horizontal gene transfer. Furthermore, the EEVS phylogeny did not follow the species phylogeny, indicating that HGT or gene conversion occurred (Figure 2). For clarity, we named the two main putative cyclitol biosynthetic gene clusters S-cluster (for streptol) and K-cluster (for kirkamide) based on previous biosynthetic hypotheses from *in silico* analysis of the putative cyclitol gene clusters of *P. kirkii* (Figure 2) (Pinto-Carbó *et al.*, 2016). Both K and S-clusters encode a core set of proteins linked to sugar analog biosynthesis: a ROK family protein and a HAD family hydrolase, and both contain aminotransferases (although from different protein families). Two EEVS genes contain nonsense mutations and are likely not functional: the S-cluster EEVS of *Ca*. Burkholderia humilis, and the K-cluster EEVS of *Ca*. Burkholderia brachyanthoides. The genome of *Ca*. B. humilis still contains an apparently functional K-cluster EEVS, while the pseudogenized EEVS of *Ca*. B. brachyanthoides is the only homolog in the genome. Interestingly, genes of the K-cluster appear to be exclusive to *Psychotria* and *Pavetta* symbionts, while the S-cluster is more widespread, including in the genomes of *Vangueria* endophytes. Accordingly, we detected kirkamide in leaf extracts of *Psychotria kirkii*, but in none of the *Fadogia* or *Vangueria* species we tested (see supplementary methods). We also detected signals that were consistent with streptol/valienol and streptol glucoside by UPLC-QToF-MS in all samples. However, these signals occurred in a noisy part of the chromatogram, and we could not confidently assign these m/z features to streptol or its derivatives (see supplementary methods).

**Figure 2:**
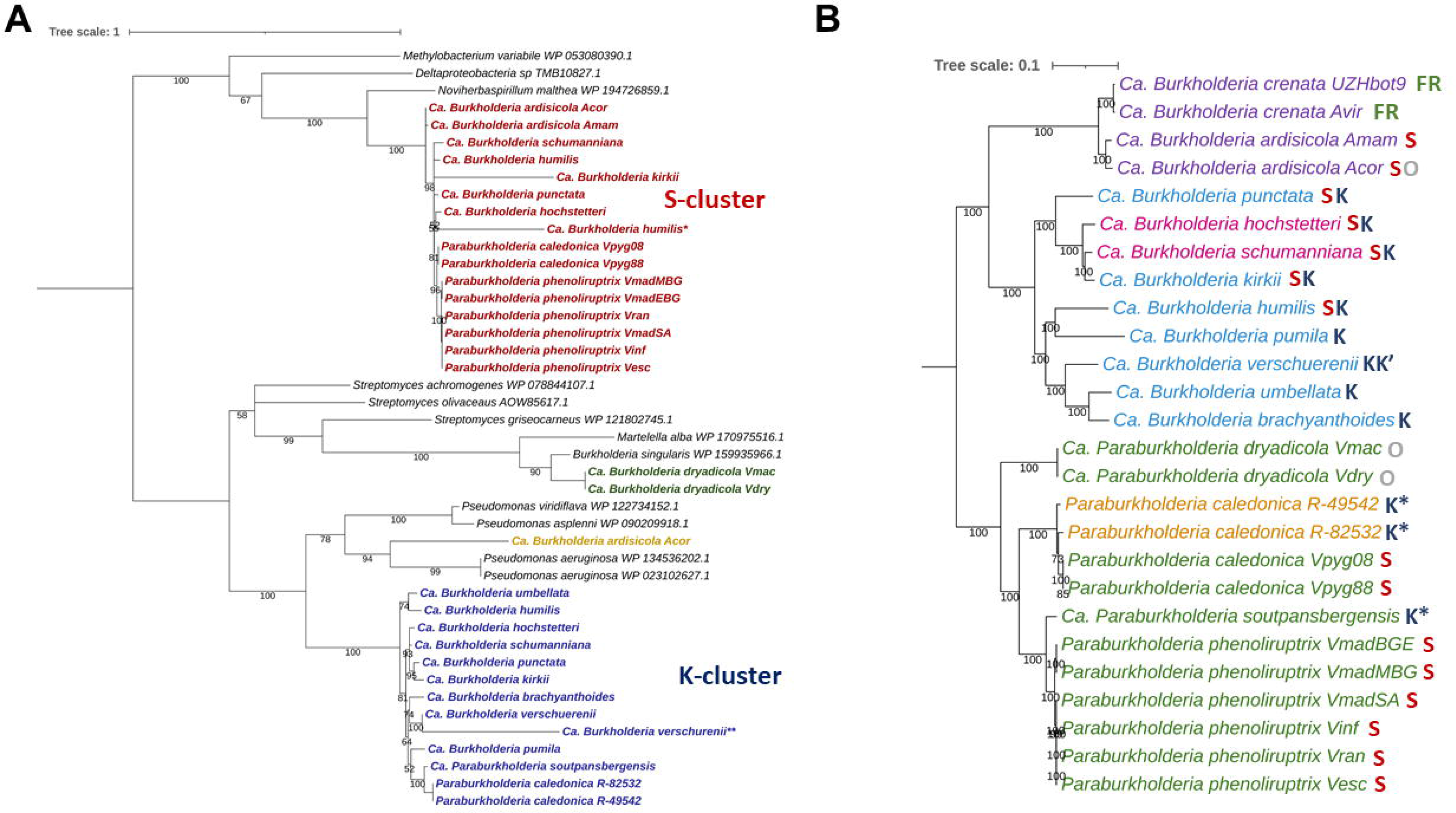
EEVS protein phylogeny and distribution in leaf endophytes. **(A)** EEVS protein phylogeny of detected EEVS-genes and their closest relatives. Local support values based on the Shimodaira-Hasegawa test are shown on the branches, and branches with support <50% are collapsed. Coloured samples in boldface are the EEVS homologs found in different leaf endophytes. Colours represent different clusters of similar EEVS genes. K- and S-cluster are named after their putative products (K for Kirkamide, and S for Streptol glucoside). NCBI accession numbers of the close relatives are given next to their species name. The tree is rooted using related 3-dehydroquinate synthase genes (not shown). *The EEVS gene in *Ca*. Burkholderia humilis contains an internal stop codon, creating two EEVS-like pseudogenes. The largest of both was used for the phylogeny. **This EEVS gene of *Ca*. Burkholderia verschuerenii is found outside of the K-cluster. **(B)** Distribution of specialised metabolism in the leaf endophytes. Samples are colour-coded based on the host species: Purple – *Ardisia*; Blue – *Psychotria*; Pink – *Pavetta*; Green – *Vangueria*; Orange – *Fadogia*. Codes next to the species represent presence of specialised metabolite clusters; FR – FR900359 depsipeptide; K – Kirkamide EEVS-cluster; S – Streptol glucoside EEVS-cluster; O – Other EEVS-cluster. K’ – Secondary EEVS cluster with EEVS similar to the K-cluster. K* - Only the K-cluster EEVS is present, not the accessory genes.

**Table 2:**
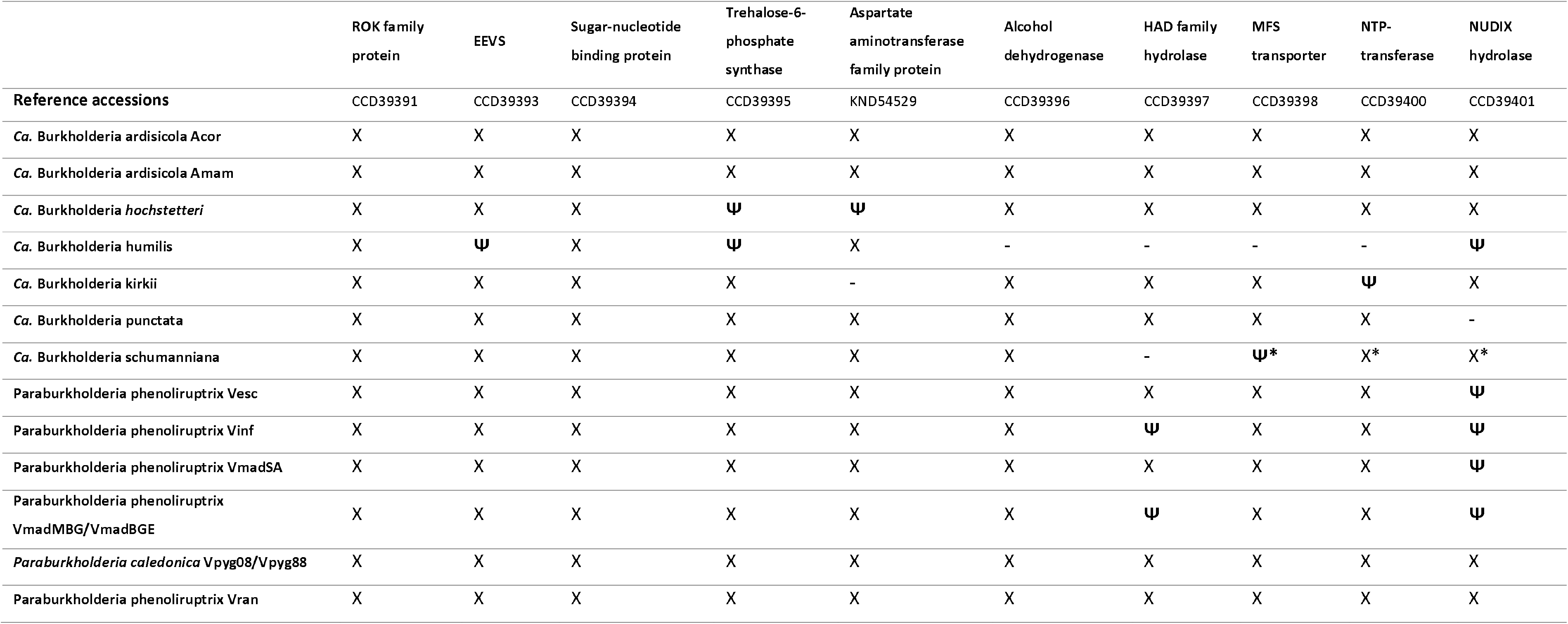
EEVS S-cluster organisation in endophyte genomes. Genomes of the same host with the same cluster layout are merged. X: Gene present; -: Gene absent; **Ψ**: Gene predicted to be pseudogene; *: genes present on a different contig than the EEVS gene; Abbreviations: EEVS – 2-*epi*-5-*epi*-valiolone synthase;

**Table 3:** EEVS K-cluster organisation in endophyte genomes. Genomes of the same host with the same cluster layout are merged. X: Gene present; -: Gene absent; **Ψ**: Gene predicted to be pseudogene; *: protein overlaps with contig end, other genes of the cluster not found on other contigs; Abbreviations: EEVS – 2-*epi*-5-*epi*-valiolone synthase.

|  | GNAT family N-acetyltransferase | Cupin Domain Containing protein | HAD family hydrolase | Gfo/Idh/MocA family oxidoreductase | 6-phospho-beta-glucosidase | DegT/DnrJ/EryC1/StrS family aminotransferase | ROK family protein | EEVS |
| --- | --- | --- | --- | --- | --- | --- | --- | --- |
| Reference accessions | CCD36711 | CCD36712 | CCD36713 | CCD36714 | CCD36715 | CCD36716 | CCD6717 | CCD36718 |
| *Paraburkholderia caledonica* R-49542/R-82532 | - | - | - | - | - | - | - | X |
| *Ca. Burkholderia brachyanthoides* | - | - | - | - | - | - | X/Ψ * | **Ψ** |
| *Ca. Burkholderia hochstetteri* | X | X | X | X | X | X | X | X |
| *Ca. Burkholderia humilis* | - | X | **Ψ** | X | X | **Ψ** | X | X |
| *Ca. Burkholderia kirkii* | X | X | X | X | X | X | X | X |
| *Ca. Burkholderia pumila* | - | X | X | X | X | X | X | X |
| *Ca. Burkholderia punctata* | X | X | X | X | X | X | X | X |
| *Ca. Burkholderia schumanniana* | X | X | X | X | X | X | X | X |
| *Ca. Burkholderia umbellata* | X | X | X | X | X | X | X | X |
| *Ca. Burkholderia verschuerenii* | X | X | X | X | X | X | X | X |
| *Ca. Paraburkholderia soutpansbergensis* | - | - | - | - | - | - | - | X |

The genomes of *Ca*. P. soutpansbergensis and *P. caledonica* R-49542 and R-82532 encoded EEVS homologs of the K-cluster, but the full complement of the genes of the K-cluster is missing (Table 3). In both cases the EEVS gene is flanked by IS elements. Accordingly, we did not detect kirkamide in leaf samples from either *Fadogia homblei* or *P. soutpansbergensis* in our chemical analyses. The genome of *Ca*. P. dryadicola encodes an EEVS that clusters outside of the K- and S-EEVS clusters. Genes with putative functions similar to those of the K-cluster are located in the vicinity of the EEVS in the genome of *Ca*. P. dryadicola: oxidoreductases, an aminotransferase, and an N-acetyltransferase (Table S8). Similarly, *Ca*. B. ardisicola Acor contains a second divergent EEVS, in addition to the S-cluster EEVS. This EEVS belongs to a larger gene cluster coding for similar functions also found in the other EEVS-clusters, but contains at least one frameshift mutation and no longer codes for a functional enzyme (Table S8). Lastly, *Ca*. B. verschuerenii contains a second, recently diverged EEVS paralog of the K-cluster. This EEVS is part of a small cluster of genes, with putative functions divergent from those found in the other EEVS-clusters and likely does not play a role in kirkamide synthesis (Table S8).

## Discussion

### Different evolutionary origins of leaf symbioses in different plant genera

In this work, we investigated the evolution of associations between *Burkholderia s. l*. bacteria and plants of the Rubiaceae and Primulaceae families, and attempted to identify key characteristics of these associations. To this end, we re-analyzed publicly available genome data from previous research, and sequenced and assembled the genomes of an additional 17 leaf endophytes. In addition to leaf endophytes which had been previously detected (Lemaire, Smets, *et al.*, 2011; Verstraete *et al.*, 2011, 2013; Ku and Hu, 2014), we document here the presence of *Burkholderia s.l*. symbionts in *Pavetta hochstetteri* and *Vangueria esculenta*, and possibly *Pavetta revoluta*. In contrast to previous findings (Lemaire, Lachenaud, *et al.*, 2012), we could not detect evidence of leaf endophytes in *Psychotria capensis*, but did confirm the absence of leaf endophytes in *Psychotria zombamontana*. Phylogenetic placement of hosts and endophytes are consistent with previous data, except for the placement of *Vangueria macrocalyx* and its endophyte (Lemaire, Lachenaud, *et al.*, 2012; Verstraete *et al.*, 2013). Both chloroplast sequences of *V. macrocalyx* and *V. dryadum* and the genomes of their endophytes were near identical while previous research showed a clear phylogenetic difference both between the host species and their endophytes (Verstraete *et al.*, 2013). Blastn analysis of plant genetic markers (ITS, petB, rpl16, trnTF) of both species against the NCBI nr database showed higher identities to markers from *Vangueria dryadum* than to those of *Vangueria macrocalyx*. However, since comparison of the vouchered *V. macrocalyx* specimen to other vouchered *Vangueria dryadum* and *V. macrocalyx* by expert botanists clearly separated both species, we decided to consider both species distinct.

Previous studies showed that Rubiaceae and Primulaceae species with heritable leaf symbionts are monophyletic within their respective genera (Lemaire, Vandamme, *et al.*, 2011; Verstraete *et al.*, 2013). Thus, while the transition to a symbiotic state arose separately in multiple plant genera, it likely evolved only once in each plant genus. The only exception is the *Psychotria* genus, where it likely arose twice: once in species forming leaf nodules, and once in species without leaf nodules (Lemaire, Lachenaud, *et al.*, 2012). The repeated emergence of leaf symbiosis is reflected on the microbial side as well. A parsimonious interpretation of whole genome phylogenetic analyses indicates that *Burkholderia* endophytes evolved independently at least 8 times, most probably from ancestors with an environmental lifestyle (Figure 1A). *Caballeronia* endophytes of *Ardisia* seem to have emerged once, with most closely related species commonly isolated from soil (Lim *et al.*, 2003; Vandamme *et al.*, 2013; Uroz and Oger, 2017). As previously reported, symbionts of *Psychotria* and *Pavetta* cluster in 3 distinct phylogenetic groups within the *Caballeronia* genus. Finally, symbionts of *Vangueria* and *Fadogia* belong to 5 distinct clades within the genus *Paraburkholderia*. Apart from *Ca*. P. dryadicola that is without closely related isolates, endophytic *Paraburkholderia* species also cluster together with species commonly isolated from soil (Verstraete *et al.*, 2014; Beukes *et al.*, 2019). High host-specificity is a hallmark of the *Psychotria, Pavetta*, and *Ardisia* leaf symbiosis, but this characteristic is not shared in *Vangueria* and *Fadogia*. Based on genome similarity, we identified at least three phylogenetically divergent endophyte species that can infect multiple hosts: *P. caledonica, P. phenoliruptrix*, and *Ca*. P. dryadicola. It is also possible that these plants are in the early stages of endophyte capture, where the plant is open to acquire endophytes from the soil, as previously hypothesized for *F. homblei* (Verstraete *et al.*, 2013). Endophytes might later evolve to become host-restricted and vertically transmitted, leading to diversification from their close relatives and forming new species. This could, for example, already be the case for *Ca*. P. soutpansbergensis, which is related to *P. phenoliruptrix* but shows a more divergent genome (ANI <95%). Overall, these results highlight the general plasticity of bacteria in the *Burkholderia s.l*., as well as the probable frequent occurrence of host-switching or horizontal transfer within leaf symbiotic associations.

### Genome reduction is a common treat of leaf endophytes

Bacterial genomes contain a wealth of information yet few leaf endophyte genomes are available. In this study we provide an additional thirteen leaf endophyte genome assemblies among which the first genomes of endophytes from *Vangueria* and *Fadogia*. Aside from the genomes of *P. caledonica* endophytes, all leaf endophyte genomes were small, mostly between 3.5 and 5 Mbp. This is well below the average 6.85 Mbp of the *Burkholderiaceae* family (Carlier *et al.*, 2016; Pinto-Carbó *et al.*, 2016). In addition to their small sizes, the genomes of *Psychotria, Pavetta*, and *Ardisia* endophytes show signs of advanced genome reduction. Only 41-70% of these genomes code for functional proteins, compared to an average of about 90% for free-living bacteria (Land *et al.*, 2015). Most of these genomes also contain a high proportion of mobile sequences, up to 9% of the total assembly. Together, this indicates ongoing reductive genome evolution, a process often observed in obligate endosymbiotic bacteria (Moran and Plague, 2004; Bennett and Moran, 2015). Interestingly, the genomes of *Vangueria* and *Fadogia* endophytes, which are not contained in leaf nodules, also show signs of genome erosion: most genomes of *P. phenoliruptrix* endophytes are at or below 5 Mbp in size, with over half of their proteome predicted as non-functional. The genomes of *Ca*. P. dryadicola even approach the level of genome reduction found in most *Psychotria* symbionts. The intermediate genome reduction in endophytes of *Vangueria* and *Fadogia* could be explained by the relatively recent origin of the symbiosis, although leaf symbiosis in *Fadogia* has been estimated to be older than in *Vangueria* (7.6 Mya vs. 3.7 Mya) (Verstraete *et al.*, 2017). Other factors likely contribute to the extent or pace of genome reduction in the endophytes, such as mode of transmission and transmission bottlenecks. The larger genome size and fewer pseudogenes compared to most other leaf endophytes may explain why we could isolate *P. caledonica* endophytes from *F. homblei*, but not other endophytes. We could not identify essential genes or pathways that were consistently missing in the genomes of *Burkholderia* endophytes. It is therefore possible that other endophytic bacteria may be culturable using more complex or tailored culture conditions.

### Secondary metabolism as key factor in the evolution of leaf symbiosis

Although leaf symbionts share a similar habitat and all belong to the *Burkholderia s. l*., their core genome is surprisingly small and consists almost entirely (95%) of genes that are considered essential for cellular life. This poor conservation of accessory functions perhaps reflects the large diversity and possible redundancy of functions encoded in the genomes of *Burkholderia s.l*. that associate with plants. Interestingly, the capacity for production of secondary metabolites is a key common trait of *Burkholderia* leaf endophytes. We previously showed that *Ca*. B. crenata produces FR900359, a cyclic depsipeptide isolated from *A. crenata* leaves (Carlier *et al.*, 2016). This non-ribosomal peptide possesses unique pharmacological properties and may contribute to the protection of the host plant against insects (Carlier *et al.*, 2016; Crüsemann *et al.*, 2018). However, our data suggests that the production of cyclitols is widespread in leaf endophytic *Burkholderia*. Indeed, with the exception of *Ca*. B. crenata cited above, we found evidence for the presence of cyclitol biosynthetic pathways in all genomes of leaf endophytic *Burkholderia*. We have previously reported the presence of two gene clusters containing a 2-*epi*-5-*epi*-valiolone synthase (EEVS) in the genomes of *Psychotria* and *Pavetta* symbionts (Pinto-Carbó *et al.*, 2016). These gene clusters are likely responsible for the production of 2 distinct cyclitols: kirkamide, a C_7_N aminocyclitol with insecticidal properties which has been detected in several *Psychotria* plants; and streptol-glucoside, a plant-growth inhibitor likewise detected in *Psychotria kirkii* (Sieber *et al.*, 2015; Pinto-Carbó *et al.*, 2016; Hsiao *et al.*, 2019). EEVS from leaf symbionts belong to four phylogenetic clusters, including the two EEVS genes previously detected in *Psychotria* and *Pavetta* symbionts (Pinto-Carbó *et al.*, 2016). Similar to these previously analysed leaf endophyte genomes, the EEVS gene clusters in the newly sequenced genomes are flanked by IS-elements, and their phylogeny is incongruent with the species phylogeny. This indicates that these genes and clusters are likely acquired via horizontal gene transfer. This hypothesis is strengthened by the fact that the closest homologs of the genes in the EEVS clusters are found in genera as diverse as *Pseudomonas, Streptomyces*, and *Noviherbaspirillum*, but are rare in the genomes of *Burkholderia s.l*. The presence of the two main EEVS gene clusters (K-cluster and S-cluster) is not strictly linked to the symbiont or host taxonomy. For example, the EEVS of the K-cluster (hypothesised to produce kirkamide) is present in all sequenced symbionts of *Psychotria* and *Pavetta* but also in the endophytes of *F. homblei* and *V. soutpansbergensis*. However, in the latter two, accessory genes of the K-cluster are absent. It is possible that this EEVS interacts with gene products of other secondary metabolite clusters (Osborn *et al.*, 2017). We also noticed that some endophyte genomes contain multiple EEVS genes or gene clusters. This could provide functional redundancy, protecting against the rampant genome erosion present in these genomes. For example, two genes of the S-cluster *Ca*. B. hochstetteri are likely pseudogenes, while the K-cluster gene is still complete. On the other hand, in *Ca*. Burkholderia humilis seven out of ten genes of the S-cluster (including the EEVS) are either missing or non-functional, and the K-cluster is heavily reduced with only four functional genes out of eight (including the EEVS). As one functional EEVS copy remains, it is possible that genes located elsewhere in the genome provide these functions, as kirkamide has previously been detected in extracts of *Psychotria humilis* (Pinto-Carbó *et al.*, 2016). Alternatively, this symbiosis may have reached a “point of no return” where host and symbiont have become dependent on each other and non-performing symbionts can become fixed in the population (Bennett and Moran, 2015).

The presence of gene clusters coding for specialised secondary metabolites in all leaf symbionts could indicate that secondary metabolite production is either a prerequisite for or a consequence of an endophytic lifestyle. The fact that *P. caledonica* leaf symbionts have EEVS genes of different origin favours the hypothesis that the acquisition of secondary metabolism precedes an endophytic lifestyle. In this case, the ancestor of both endophytes may have acquired differing EEVS genes or EEVS gene clusters through HGT followed by infection of the respective host plants. The lack of EEVS homolog in *Ca*. B. crenata indicates that production of cyclitols is not essential for leaf symbiosis. Interestingly, genomes of the sister species *Ca*. B. ardisicola encode an EEVS and the full S-cluster complement. Since there is strong phylogenetic evidence of co-speciation in the *Burkholderia/Ardisia* association (Lemaire, Smets, *et al.*, 2011; Ku and Hu, 2014), the common ancestor of *Ca*. B. ardisicola and *Ca*. B. crenata possibly possessed both cyclitols and *frs* pathways, and one of these pathways was lost in the lineages leading to contemporary *Ca*. B. crenata and *Ca*. B. ardisicola. Alternatively, the genome of the common ancestor of *Ardisia*-associated *Burkholderia* may have encoded cyclitol S-cluster and later acquisition of the *frs* gene cluster in the Ca. B. crenata lineage alleviated the requirement of EEVS-related metabolism. The model of horizontal acquisition of secondary functions supports the model of endophyte evolution described by Lemaire *et al* (Lemaire, Vandamme, *et al.*, 2011). Different environmental strains which acquired genes for secondary metabolite production could colonise different host plants in the early open phase of symbiosis. The different phylogenetic endophyte clades observed in the *Burkholderia s.l*. phylogeny could each represent distinct acquisitions of secondary metabolite gene clusters by divergent free-living bacteria followed by colonisation of different host plants. Many *Burkholderia* species associate with eukaryotic hosts, including plants (Eberl and Vandamme, 2016), and many of these associations may be transient in nature. However, useful traits such as synthesis of protective metabolites may help stabilise these relationships, resulting in long-term associations such as leaf symbiosis.

## Author contributions

AC, MM, and BD designed the research. MM identified and collected wild plant specimens from the Pretoria region (South Africa). BD, MB, SS, and AC performed the laboratory experiments and analyses. BD, MM and AC wrote the manuscript with input from all authors.

## Acknowledgments

We would like to thank Frédéric De Meyer and Mathijs Deprez for helping with some of the laboratory experiments. We would further like to thank Steven Janssens (Meise Botanic Garden) and Peter Brownless (Royal Botanic Garden Edinburgh) for facilitating the acquisition of plant material for this study. BD and AC would like to thank Klaas Vandepoele, Monica Höfte, Anne Willems and Paul Wilkin for helpful discussion and for proofreading the manuscript. We also thank Aurélien Bailly (University of Zürich, CH) for providing P. kirkii samples and help with interpreting mass spectrometry data. Magda Nel of the H.G.W.J. Schweickerdt Herbarium is thanked for her help with plant identification and Mamoalosi Selepe and Sewes Alberts of the Chemistry and Plant and Soil Sciences Departments, respectively (University of Pretoria) for chemical analysis. We also thank Chien-Chi Hsiao and Karl Gademann from University of Zürich (Switzerland) for providing the analytical standards. This work was supported by the Flemish Fonds Wetenschappelijk Onderzoek under grant G017717N to AC. AC also acknowledges support from the French National Research Agency under grant agreement ANR-19-TERC-0004-01 and from the French Laboratory of Excellence project “TULIP” (ANR-10-LABX-41; ANR-11-IDEX-0002-02) and from the French National Infrastructure for Metabolomics and Fluxomics, Grant MetaboHUB-ANR-11-INBS-0010. We thank the Oxford Genomics Centre at the Wellcome Centre for Human Genetics for the collection and preliminary analysis of sequencing data. The Oxford Genomics Centre at the Wellcome Centre for Human Genetics is funded by Wellcome Trust grant reference 203141/Z/16/Z. The funders had no role in study design, data collection and analysis, decision to publish, or preparation of the manuscript.

## Notes

The authors declare no conflict of interest.

## References

Abby, S.S., Cury, J., Guglielmini, J., Néron, B., Touchon, M., and Rocha, E.P.C. (2016) Identification of protein secretion systems in bacterial genomes. Sci Rep 6: 23080.

Abby, S.S., Néron, B., Ménager, H., Touchon, M., and Rocha, E.P.C. (2014) MacSyFinder: A Program to Mine Genomes for Molecular Systems with an Application to CRISPR-Cas Systems. PLoS One 9: e110726.

Bankevich, A., Nurk, S., Antipov, D., Gurevich, A.A., Dvorkin, M., Kulikov, A.S., et al. (2012) SPAdes: A new genome assembly algorithm and its applications to single-cell sequencing. J Comput Biol 19: 455–477.

Bennett, G.M. and Moran, N.A. (2015) Heritable symbiosis: The advantages and perils of an evolutionary rabbit hole. Proc Natl Acad Sci U S A 112: 10169–76.

Bertels, F., Silander, O.K., Pachkov, M., Rainey, P.B., and van Nimwegen, E. (2014) Automated reconstruction of whole-genome phylogenies from short-sequence reads. Mol Biol Evol 31: 1077–88.

Beukes, C.W., Steenkamp, E.T., van Zyl, E., Avontuur, J., Chan, W.Y., Hassen, A.I., et al. (2019) Paraburkholderia strydomiana sp. nov. and Paraburkholderia steynii sp. nov.: rhizobial symbionts of the fynbos legume Hypocalyptus sophoroides. Antonie Van Leeuwenhoek 112: 1369–1385.

Brader, G., Compant, S., Vescio, K., Mitter, B., Trognitz, F., Ma, L.-J., and Sessitsch, A. (2017) Ecology and Genomic Insights into Plant-Pathogenic and Plant-Nonpathogenic Endophytes. Annu Rev Phytopathol 55: 61–83.

Brettin, T., Davis, J.J., Disz, T., Edwards, R.A., Gerdes, S., Olsen, G.J., et al. (2015) RASTtk: A modular and extensible implementation of the RAST algorithm for building custom annotation pipelines and annotating batches of genomes. Sci Rep 5: 8365.

Brundrett, M. (1991) Mycorrhizas in Natural Ecosystems. In Advances in Ecological Research. pp. 171–313.

Buchfink, B., Reuter, K., and Drost, H.-G. (2021) Sensitive protein alignments at tree-of-life scale using DIAMOND. Nat Methods 18: 366–368.

Camacho, C., Coulouris, G., Avagyan, V., Ma, N., Papadopoulos, J., Bealer, K., and Madden, T.L. (2009) BLAST+: architecture and applications. BMC Bioinformatics 10: 421.

Cantalapiedra, C.P., Hernández-Plaza, A., Letunic, I., Bork, P., and Huerta-Cepas, J. (2021) eggNOG-mapper v2: Functional Annotation, Orthology Assignments, and Domain Prediction at the Metagenomic Scale. bioRxiv 2021.06.03.446934.

Carlier, A. and Eberl, L. (2012) The eroded genome of a Psychotria leaf symbiont: Hypotheses about lifestyle and interactions with its plant host. Environ Microbiol 14: 2757–2769.

Carlier, A., Fehr, L., Pinto-Carbó, M., Schäberle, T., Reher, R., Dessein, S., et al. (2016) The genome analysis of Candidatus Burkholderia crenata reveals that secondary metabolism may be a key function of the Ardisia crenata leaf nodule symbiosis. Environ Microbiol 18: 2507–2522.

Carlier, A.L., Omasits, U., Ahrens, C.H., and Eberl, L. (2013) Proteomics analysis of Psychotria leaf nodule symbiosis: improved genome annotation and metabolic predictions. Mol Plant Microbe Interact 26: 1325–33.

Carver, T., Harris, S.R., Berriman, M., Parkhill, J., and McQuillan, J.A. (2012) Artemis: an integrated platform for visualization and analysis of high-throughput sequence-based experimental data. Bioinformatics 28: 464–469.

Chen, S., Zhou, Y., Chen, Y., and Gu, J. (2018) fastp: an ultra-fast all-in-one FASTQ preprocessor. Bioinformatics 34: i884–i890.

Crüsemann, M., Reher, R., Schamari, I., Brachmann, A.O., Ohbayashi, T., Kuschak, M., et al. (2018) Heterologous Expression, Biosynthetic Studies, and Ecological Function of the Selective Gq-Signaling Inhibitor FR900359. Angew Chemie Int Ed 57: 836–840.

Dangl, J.L. and Jones, J.D.G. (2001) Plant pathogens and integrated defence responses to infection. Nature 411: 826–833.

Eberl, L. and Vandamme, P. (2016) Members of the genus Burkholderia: good and bad guys. F1000Research 5: 1007.

Edwards, U., Rogall, T., Blöcker, H., Emde, M., and Böttger, E.C. (1989) Isolation and direct complete nucleotide determination of entire genes. Characterization of a gene coding for 16S ribosomal RNA. Nucleic Acids Res 17: 7843–7853.

Emms, D.M. and Kelly, S. (2019) OrthoFinder: Phylogenetic orthology inference for comparative genomics. Genome Biol 20: 238.

Fisher, R.M., Henry, L.M., Cornwallis, C.K., Kiers, E.T., and West, S.A. (2017) The evolution of host-symbiont dependence. Nat Commun 8: 15973.

Fujioka, M., Koda, S., Morimoto, Y., and Biemann, K. (1988) Structure of FR900359, a cyclic depsipeptide from Ardisia crenata sims. J Org Chem 53: 2820–2825.

Gaspar, J.M. (2018) NGmerge: merging paired-end reads via novel empirically-derived models of sequencing errors. BMC Bioinformatics 19: 536.

Georgiou, A., Sieber, S., Hsiao, C.C., Grayfer, T., Gorenflos López, J.L., Gademann, K., et al. (2021) Leaf nodule endosymbiotic Burkholderia confer targeted allelopathy to their Psychotria hosts. Sci Rep 11: 1–15.

Guindon, S., Dufayard, J.-F., Lefort, V., Anisimova, M., Hordijk, W., and Gascuel, O. (2010) New algorithms and methods to estimate maximum-likelihood phylogenies: assessing the performance of PhyML 3.0. Syst Biol 59: 307–21.

Gundel, P.E., Rudgers, J.A., and Whitney, K.D. (2017) Vertically transmitted symbionts as mechanisms of transgenerational effects. Am J Bot 104: 787–792.

Gurevich, A., Saveliev, V., Vyahhi, N., and Tesler, G. (2013) QUAST: quality assessment tool for genome assemblies. Bioinformatics 29: 1072–1075.

Heberle, H., Meirelles, G.V., da Silva, F.R., Telles, G.P., and Minghim, R. (2015) InteractiVenn: a web-based tool for the analysis of sets through Venn diagrams. BMC Bioinformatics 16: 169.

Hsiao, C.-C., Sieber, S., Georgiou, A., Bailly, A., Emmanouilidou, D., Carlier, A., et al. (2019) Synthesis and Biological Evaluation of the Novel Growth Inhibitor Streptol Glucoside, Isolated from an Obligate Plant Symbiont. Chem - A Eur J 25: 1722–1726.

Huerta-Cepas, J., Szklarczyk, D., Heller, D., Hernández-Plaza, A., Forslund, S.K., Cook, H., et al. (2019) eggNOG 5.0: a hierarchical, functionally and phylogenetically annotated orthology resource based on 5090 organisms and 2502 viruses. Nucleic Acids Res 47: D309–D314.

Inglis, P.W., Pappas, M. de C.R., Resende, L. V., and Grattapaglia, D. (2018) Fast and inexpensive protocols for consistent extraction of high quality DNA and RNA from challenging plant and fungal samples for high-throughput SNP genotyping and sequencing applications. PLoS One 13: e0206085.

Ku, C. and Hu, J.-M.M. (2014) Phylogenetic and Cophylogenetic Analyses of the Leaf-Nodule Symbiosis in Ardisia Subgenus Crispardisia (Myrsinaceae): Evidence from Nuclear and Chloroplast Markers and Bacterial rrn Operons. Int J Plant Sci 175: 92–109.

Land, M., Hauser, L., Jun, S.R., Nookaew, I., Leuze, M.R., Ahn, T.H., et al. (2015) Insights from 20 years of bacterial genome sequencing. Funct Integr Genomics 15: 141–161.

Lemaire, B., Lachenaud, O., Persson, C., Smets, E., and Dessein, S. (2012) Screening for leaf-associated endophytes in the genus Psychotria (Rubiaceae). FEMS Microbiol Ecol 81: 364–372.

Lemaire, B., Van Oevelen, S., De Block, P., Verstraete, B., Smets, E., Prinsen, E., and Dessein, S. (2012) Identification of the bacterial endosymbionts in leaf nodules of Pavetta (Rubiaceae). Int J Syst Evol Microbiol 62: 202–209.

Lemaire, B., Robbrecht, E., van Wyk, B., Van Oevelen, S., Verstraete, B., Prinsen, E., et al. (2011) Identification, origin, and evolution of leaf nodulating symbionts of Sericanthe (Rubiaceae). J Microbiol 49: 935–941.

Lemaire, B., Smets, E., and Dessein, S. (2011) Bacterial leaf symbiosis in Ardisia (Myrsinoideae, Primulaceae): molecular evidence for host specificity. Res Microbiol 162: 528–534.

Lemaire, B., Vandamme, P., Merckx, V., Smets, E., and Dessein, S. (2011) Bacterial Leaf Symbiosis in Angiosperms: Host Specificity without Co-Speciation. PLoS One 6: e24430.

Letunic, I. and Bork, P. (2019) Interactive Tree Of Life (iTOL) v4: recent updates and new developments. Nucleic Acids Res 47: W256–W259.

Li, H., Handsaker, B., Wysoker, A., Fennell, T., Ruan, J., Homer, N., et al. (2009) The Sequence Alignment/Map format and SAMtools. Bioinformatics 25: 2078–2079.

Lim, Y.W., Baik, K.S., Han, S.K., Kim, S.B., and Bae, K.S. (2003) Burkholderia sordidicola sp. nov., isolated from the white-rot fungus Phanerochaete sordida. Int J Syst Evol Microbiol 53: 1631–1636.

Lo, W.S., Huang, Y.Y., and Kuo, C.H. (2016) Winding paths to simplicity: Genome evolution in facultative insect symbionts. FEMS Microbiol Rev 40: 855–874.

Lòpez-Fernàndez, S., Sonego, P., Moretto, M., Pancher, M., Engelen, K., Pertot, I., and Campisano, A. (2015) Whole-genome comparative analysis of virulence genes unveils similarities and differences between endophytes and other symbiotic bacteria. Front Microbiol 6: 419.

Mahmud, T. (2009) Progress in aminocyclitol biosynthesis. Curr Opin Chem Biol 13: 161–170.

Mahmud, T. (2003) The C7N aminocyclitol family of natural products. Nat Prod Rep 20: 137–166.

Manzano-Marín, A., Coeur d’acier, A., Clamens, A.-L., Orvain, C., Cruaud, C., Barbe, V., and Jousselin, E. (2018) A Freeloader? The Highly Eroded Yet Large Genome of the Serratia symbiotica Symbiont of Cinara strobi. Genome Biol Evol 10: 2178–2189.

Manzano-Marín, A. and Latorre, A. (2016) Snapshots of a shrinking partner: Genome reduction in Serratia symbiotica. Sci Rep 6: 32590.

Marçais, G., Delcher, A.L., Phillippy, A.M., Coston, R., Salzberg, S.L., and Zimin, A. (2018) MUMmer4: A fast and versatile genome alignment system. PLOS Comput Biol 14: e1005944.

McCann, H.C. (2020) Skirmish or war: the emergence of agricultural plant pathogens. Curr Opin Plant Biol 56: 147–152.

McCutcheon, J.P. and Moran, N.A. (2012) Extreme genome reduction in symbiotic bacteria. Nat Rev Microbiol 10: 13–26.

Meier-Kolthoff, J.P. and Göker, M. (2019) TYGS is an automated high-throughput platform for state-of-the-art genome-based taxonomy. Nat Commun 10: 2182.

Miller, I.J., Rees, E.R., Ross, J., Miller, I., Baxa, J., Lopera, J., et al. (2019) Autometa: automated extraction of microbial genomes from individual shotgun metagenomes. Nucleic Acids Res 47: e57–e57.

Miller, I.M. (1990) Bacterial Leaf Nodule Symbiosis. Adv Bot Res 17: 163–234.

Miller, I.M. and Donnelly, A.E. (1987) Location and distribution of symbiotic bacteria during floral development in Ardisia crispa. Plant, Cell Environ 10: 715–724.

Mira, A., Ochman, H., and Moran, N.A.N.A. (2001) Deletional bias and the evolution of bacterial genomes. Trends Genet 17: 589–596.

Moran, N.A., McCutcheon, J.P., and Nakabachi, A. (2008) Genomics and Evolution of Heritable Bacterial Symbionts. Annu Rev Genet 42: 165–190.

Moran, N.A. and Plague, G.R. (2004) Genomic changes following host restriction in bacteria. Curr Opin Genet Dev 14: 627–33.

Na, S.I., Kim, Y.O., Yoon, S.H., Ha, S. min, Baek, I., and Chun, J. (2018) UBCG: Up-to-date bacterial core gene set and pipeline for phylogenomic tree reconstruction. J Microbiol 56: 281–285.

Nurk, S., Meleshko, D., Korobeynikov, A., and Pevzner, P.A. (2017) metaSPAdes: a new versatile metagenomic assembler. Genome Res 27: 824–834.

Van Oevelen, S., De Wachter, R., Vandamme, P., Robbrecht, E., and Prinsen, E. (2002) Identification of the bacterial endosymbionts in leaf galls of Psychotria (Rubiaceae, angiosperms) and proposal of “Candidatus Burkholderia kirkii” sp. nov. Int J Syst Evol Microbiol 52: 2023–2027.

Ondov, B.D., Bergman, N.H., and Phillippy, A.M. (2011) Interactive metagenomic visualization in a Web browser. BMC Bioinformatics 12: 385.

Osborn, A.R., Almabruk, K.H., Holzwarth, G., Asamizu, S., LaDu, J., Kean, K.M., et al. (2015) De novo synthesis of a sunscreen compound in vertebrates. Elife 4:.

Osborn, A.R., Kean, K.M., Alseud, K.M., Almabruk, K.H., Asamizu, S., Lee, J.A., et al. (2017) Evolution and Distribution of C 7–Cyclitol Synthases in Prokaryotes and Eukaryotes. ACS Chem Biol 12: 979–988.

Pearson, W.R. (2000) Flexible Sequence Similarity Searching with the FASTA3 Program Package. In Bioinformatics Methods and Protocols. New Jersey: Humana Press, pp. 185–219.

Pettersson, M.E. and Berg, O.G. (2007) Muller’s ratchet in symbiont populations. Genetica 130: 199–211.

Pinto-Carbó, M., Sieber, S., Dessein, S., Wicker, T., Verstraete, B., Gademann, K., et al. (2016) Evidence of horizontal gene transfer between obligate leaf nodule symbionts. ISME J 10: 2092–105.

Ponsting, H. and Ning, Z. (2010) SMALT - A New Mapper for DNA Sequencing Reads. F1000Posters 1: 1.

Price, M.N., Dehal, P.S., and Arkin, A.P. (2009) FastTree: computing large minimum evolution trees with profiles instead of a distance matrix. Mol Biol Evol 26: 1641–50.

Prjibelski, A., Antipov, D., Meleshko, D., Lapidus, A., and Korobeynikov, A. (2020) Using SPAdes De Novo Assembler. Curr Protoc Bioinforma 70:.

van Rhijn, P. and Vanderleyden, J. (1995) The Rhizobium-plant symbiosis. Microbiol Rev 59: 124–142.

Richter, M., Rosselló-Móra, R., Oliver Glöckner, F., and Peplies, J. (2016) JSpeciesWS: a web server for prokaryotic species circumscription based on pairwise genome comparison. Bioinformatics 32: 929–931.

Schneider, M., Tognolli, M., and Bairoch, A. (2004) The Swiss-Prot protein knowledgebase and ExPASy: providing the plant community with high quality proteomic data and tools. Plant Physiol Biochem 42: 1013–1021.

Shigenobu, S., Watanabe, H., Hattori, M., Sakaki, Y., and Ishikawa, H. (2000) Genome sequence of the endocellular bacterial symbiont of aphids Buchnera sp. APS. Nature 407: 81–86.

Sieber, S., Carlier, A., Neuburger, M., Grabenweger, G., Eberl, L., and Gademann, K. (2015) Isolation and Total Synthesis of Kirkamide, an Aminocyclitol from an Obligate Leaf Nodule Symbiont. Angew Chemie Int Ed 54: 7968–7970.

Sinnesael, A., Eeckhout, S., Janssens, S.B., Smets, E., Panis, B., Leroux, O., and Verstraete, B. (2018) Detection of Burkholderia in the seeds of Psychotria punctata (Rubiaceae) – Microscopic evidence for vertical transmission in the leaf nodule symbiosis. PLoS One 13: e0209091.

Smith, S.E. and Read, D.J. (2008) Mycorrhizal Symbiosis, Academic Press.

Souvorov, A., Agarwala, R., and Lipman, D.J. (2018) SKESA: strategic k-mer extension for scrupulous assemblies. Genome Biol 19: 153.

Stamatakis, A. (2014) RAxML version 8: A tool for phylogenetic analysis and post-analysis of large phylogenies. Bioinformatics 30: 1312–1313.

Toh, H., Weiss, B.L., Perkin, S.A.H., Yamashita, A., Oshima, K., Hattori, M., and Aksoy, S. (2006) Massive genome erosion and functional adaptations provide insights into the symbiotic lifestyle of Sodalis glossinidius in the tsetse host. Genome Res 16: 149–156.

Di Tommaso, P., Moretti, S., Xenarios, I., Orobitg, M., Montanyola, A., Chang, J.M., et al. (2011) T-Coffee: A web server for the multiple sequence alignment of protein and RNA sequences using structural information and homology extension. Nucleic Acids Res 39: W13–7.

Uroz, S. and Oger, P. (2017) Caballeronia mineralivorans sp. nov., isolated from oak-Scleroderma citrinum mycorrhizosphere. Syst Appl Microbiol 40: 345–351.

Vandamme, P., De Brandt, E., Houf, K., Salles, J.F., van Elsas, J.D., Spilker, T., and LiPuma, J.J. (2013) Burkholderia humi sp. nov., Burkholderia choica sp. nov., Burkholderia telluris sp. nov., Burkholderia terrestris sp. nov. and Burkholderia udeis sp. nov.: Burkholderia glathei-like bacteria from soil and rhizosphere soil. Int J Syst Evol Microbiol 63: 4707–4718.

Vandamme, P., Peeters, C., De Smet, B., Price, E.P., Sarovich, D.S., Henry, D.A., et al. (2017) Comparative Genomics of Burkholderia singularis sp. nov., a Low G+C Content, Free-Living Bacterium That Defies Taxonomic Dissection of the Genus Burkholderia. Front Microbiol 8: 1679.

Verstraete, B., Van Elst, D., Steyn, H., Van Wyk, B., Lemaire, B., Smets, E., and Dessein, S. (2011) Endophytic Bacteria in Toxic South African Plants: Identification, Phylogeny and Possible Involvement in Gousiekte. PLoS One 6: e19265.

Verstraete, B., Janssens, S., and Rønsted, N. (2017) Non-nodulated bacterial leaf symbiosis promotes the evolutionary success of its host plants in the coffee family (Rubiaceae). Mol Phylogenet Evol 113: 161–168.

Verstraete, B., Janssens, S., Smets, E., and Dessein, S. (2013) Symbiotic ß-Proteobacteria beyond Legumes: Burkholderia in Rubiaceae. PLoS One 8: e55260.

Verstraete, B., Peeters, C., van Wyk, B., Smets, E., Dessein, S., and Vandamme, P. (2014) Intraspecific variation in Burkholderia caledonica: Europe vs. Africa and soil vs. endophytic isolates. Syst Appl Microbiol 37: 194–9.

Vessey, J.K., Pawlowski, K., and Bergman, B. (2005) Root-based N2-fixing Symbioses: Legumes, Actinorhizal Plants, Parasponia sp. and Cycads. Plant Soil 274: 51–78.

Wilson, K. (2001) Preparation of Genomic <scp>DNA</scp> from Bacteria. Curr Protoc Mol Biol 56: 2.4.1–2.4.5.

Wood, D.E., Lu, J., and Langmead, B. (2019) Improved metagenomic analysis with Kraken 2. Genome Biol 20: 257.

Wu, X., Flatt, P.M., Schlörke, O., Zeeck, A., Dairi, T., and Mahmud, T. (2007) A Comparative Analysis of the Sugar Phosphate Cyclase Superfamily Involved in Primary and Secondary Metabolism. ChemBioChem 8: 239–248.

Xie, Z. and Tang, H. (2017) ISEScan: automated identification of insertion sequence elements in prokaryotic genomes. Bioinformatics 33: 3340–3347.

Zhang, R. (2004) DEG: a database of essential genes. Nucleic Acids Res 32: 271D–272.

