## Supplementary Figures for "Cyclitol secondary metabolism is a central feature of *Burkholderia* leaf symbionts"

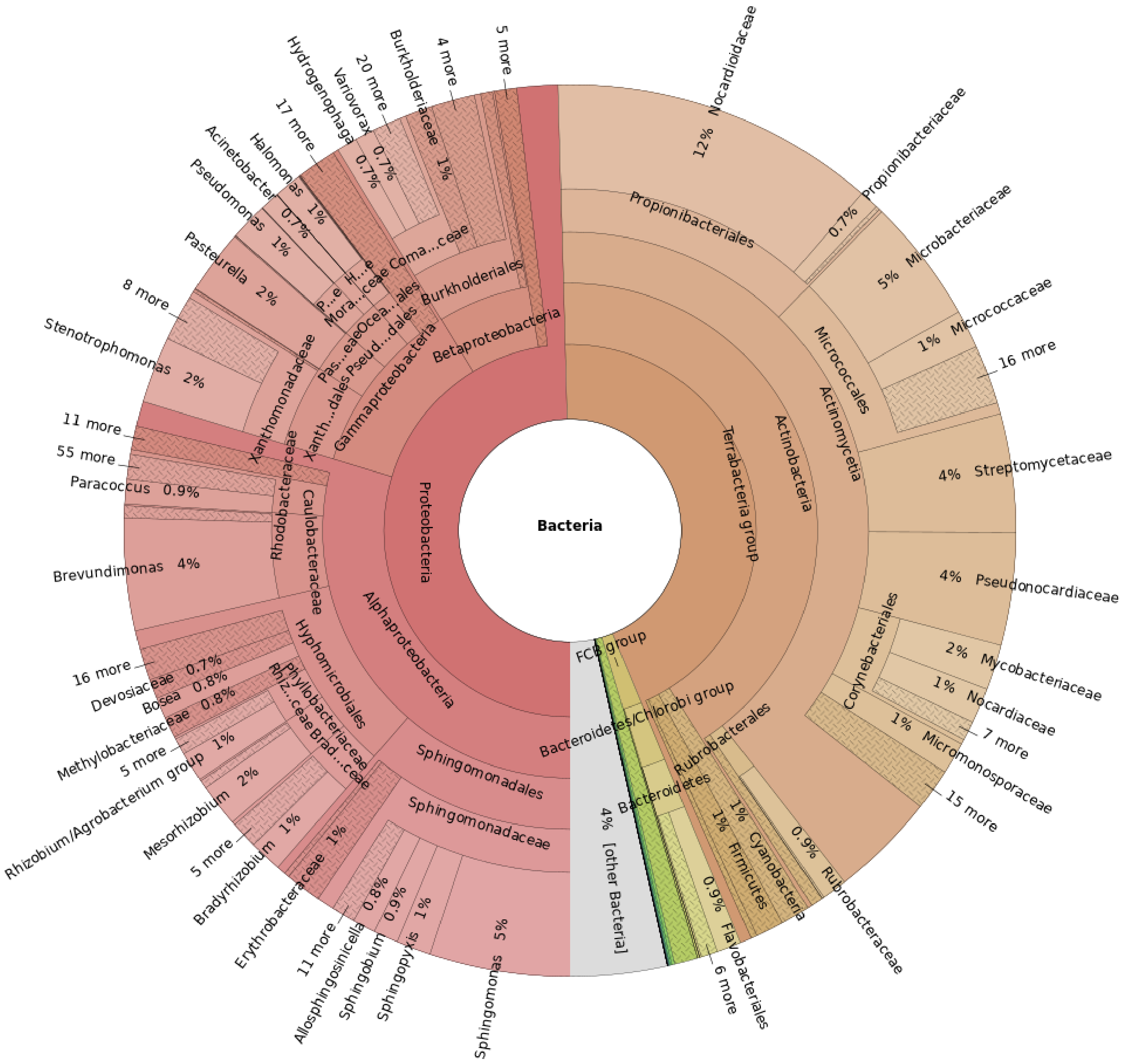


**Figure S1: Taxonomic distribution of bacterial reads from leaves of *Pavetta indica*.** Estimation based on blastn searches against the NCBI nucleotide database on a subset of 1M reads. Only bacterial reads (representing an estimated 7% of total reads) are shown here.

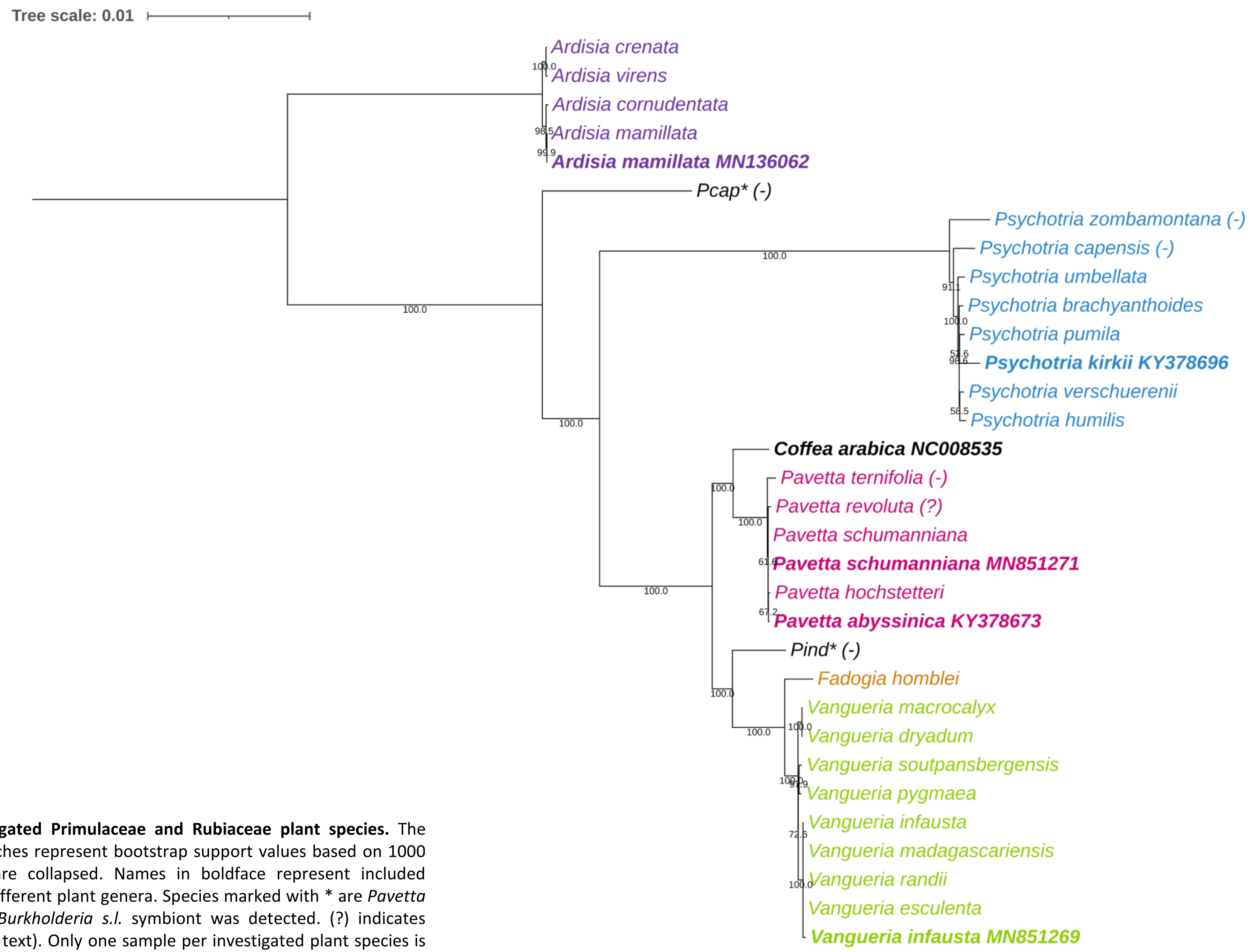


**Figure S2: SNP-based chloroplast phylogeny of investigated Primulaceae and Rubiaceae plant species.** The *Ardisia* clade was set as outgroup. Numbers on the branches represent bootstrap support values based on 1000 replications. Branches with <50% bootstrap support are collapsed. Names in boldface represent included reference chloroplast sequences. Colours represent the different plant genera. Species marked with * are *Pavetta* species likely misidentified (see text). (-) indicates no *Burkholderia s.l.* symbiont was detected. (?) indicates presence of a *Burkholderia s.l.* symbiont is uncertain (see text). Only one sample per investigated plant species is included in the phylogenetic tree. *Pcap: Psychotria capensis*; *Pind*: *Psychotria indica*

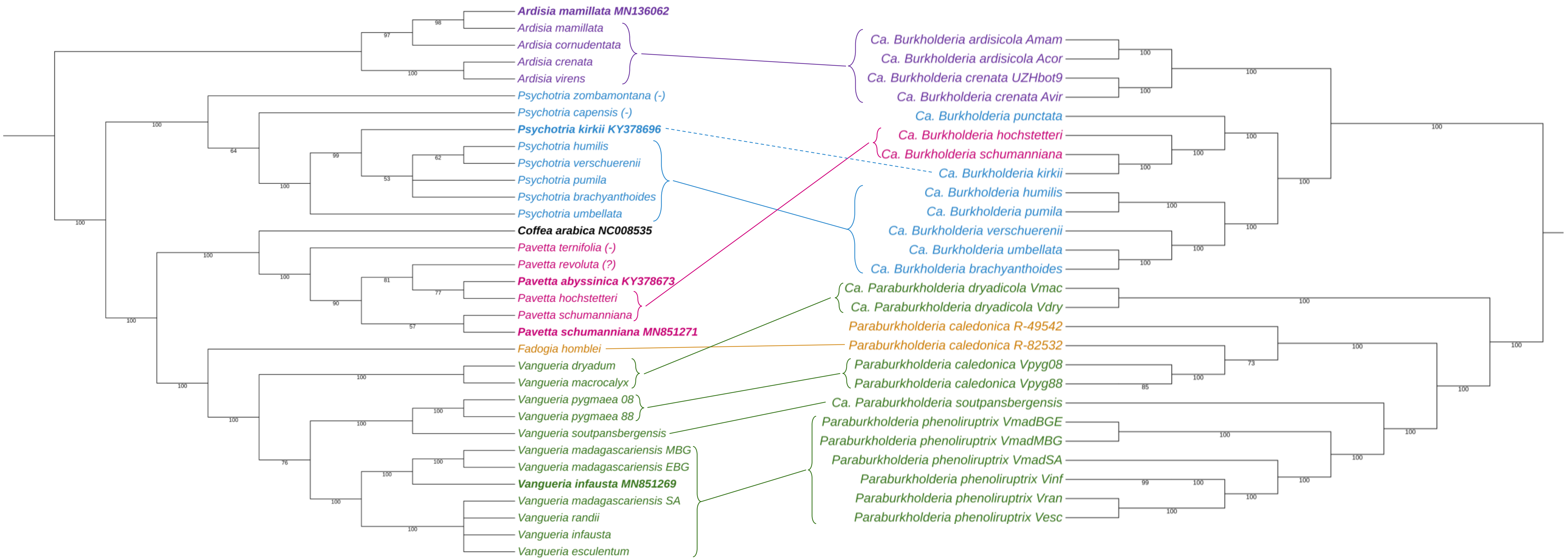


**Figure S3: Cophylogenetic patterns between leaf endophytes and their hosts.** Left: Chloroplast phylogeny of host plants. Right: Core-genome phylogeny of leaf endophytes. Branch lengths are not representative. Numbers on branches represent bootstrap support values based on 100 bootstrap replications. Connections were drawn between representative groups of endophytes and their host plants. The dotted line for *Psychotria kirkii* indicates that the endophyte and host plant are not derived from the same voucher.

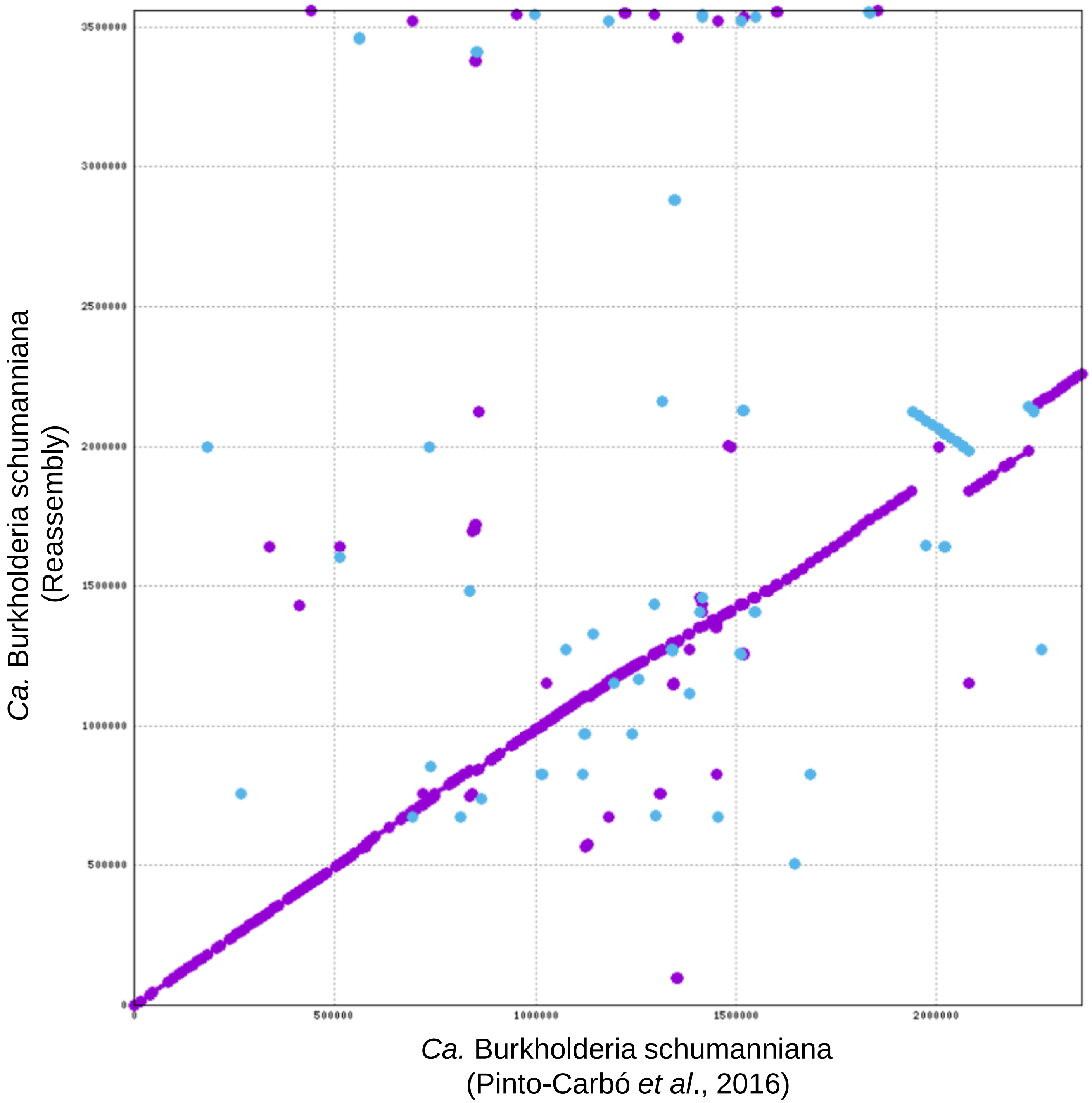


**Figure S4: Dotplot of the reassembly of *Ca.* Burkholderia schumanniana versus the published genome sequence.** Numbers on X- and Y-axis represent the position in the assembly, in bases. Dots represent aligned subsequences between both assemblies. Purple dots represent sequences aligned in the same sense, while blue dots represent sequences aligned in opposite sense.

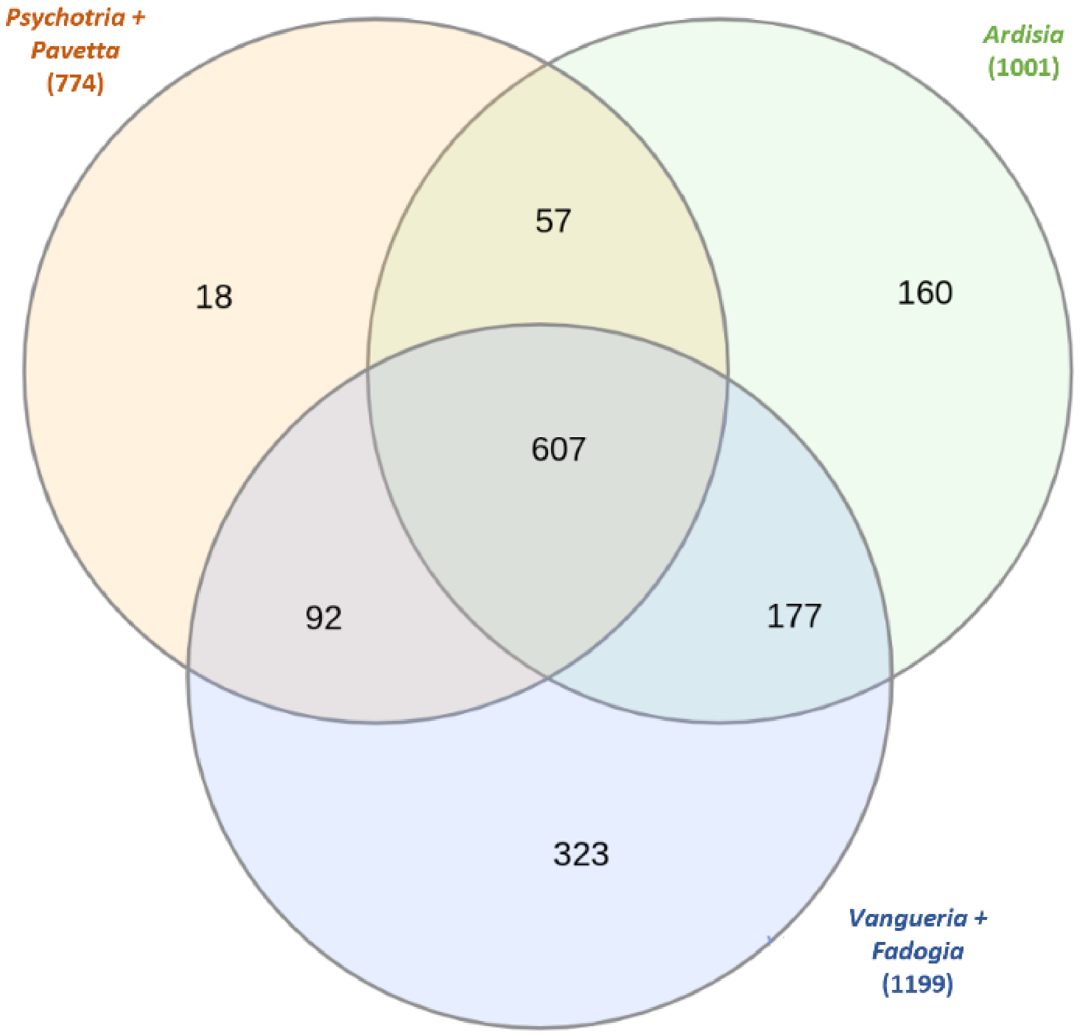


**Figure S5: Core-genome overlap between leaf endophytes of different plant genera.** Venn diagram showing the overlap of the core genome between endophytes of different plant hosts. Numbers between brackets represent the total core genome size of a certain clade.

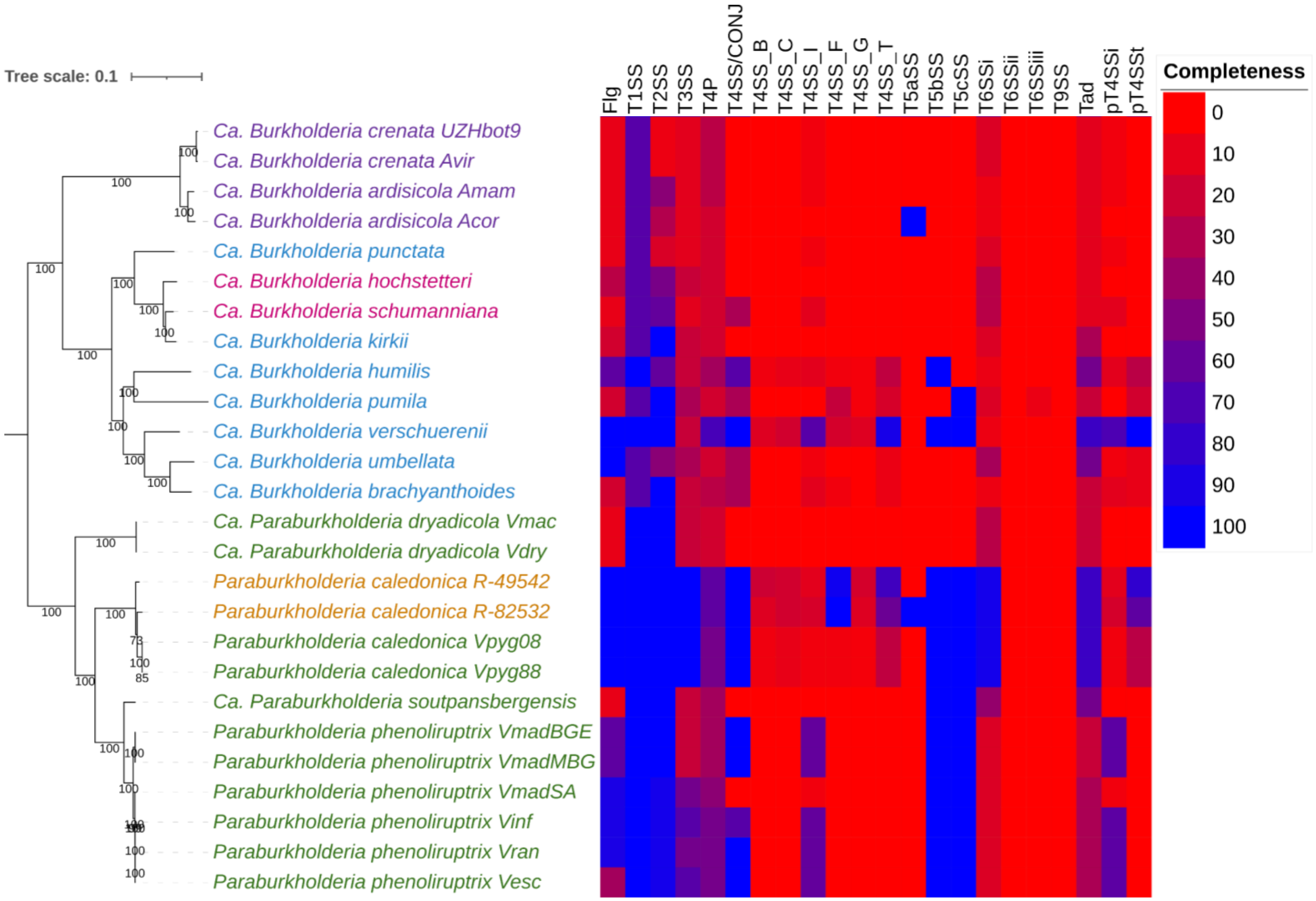


**Figure S6: Presence and completeness of flagellar apparatus and secretion systems in leaf endophytes.** Completeness refers to the proportion of genes in a certain cluster that are found present in a certain genome. Coloured names represent the host species of the endophytes: Purple – *Ardisia* spp.; Blue – *Psychotria* spp.; Pink – *Pavetta* spp.; Green – *Vangueria* spp.; Orange – *Fadogia* spp. Abbreviations: SA – South Africa; 08/88 last two digits of *V. pygmaea* voucher number; BGE – Botanic Garden Edinburgh; MBG – Meise Botanic Garden; Flg – Flagellar apparatus; TXSS – Type X Secretion System; T4P – Type IV Pilus; Tad – Tight Adherence pilus,

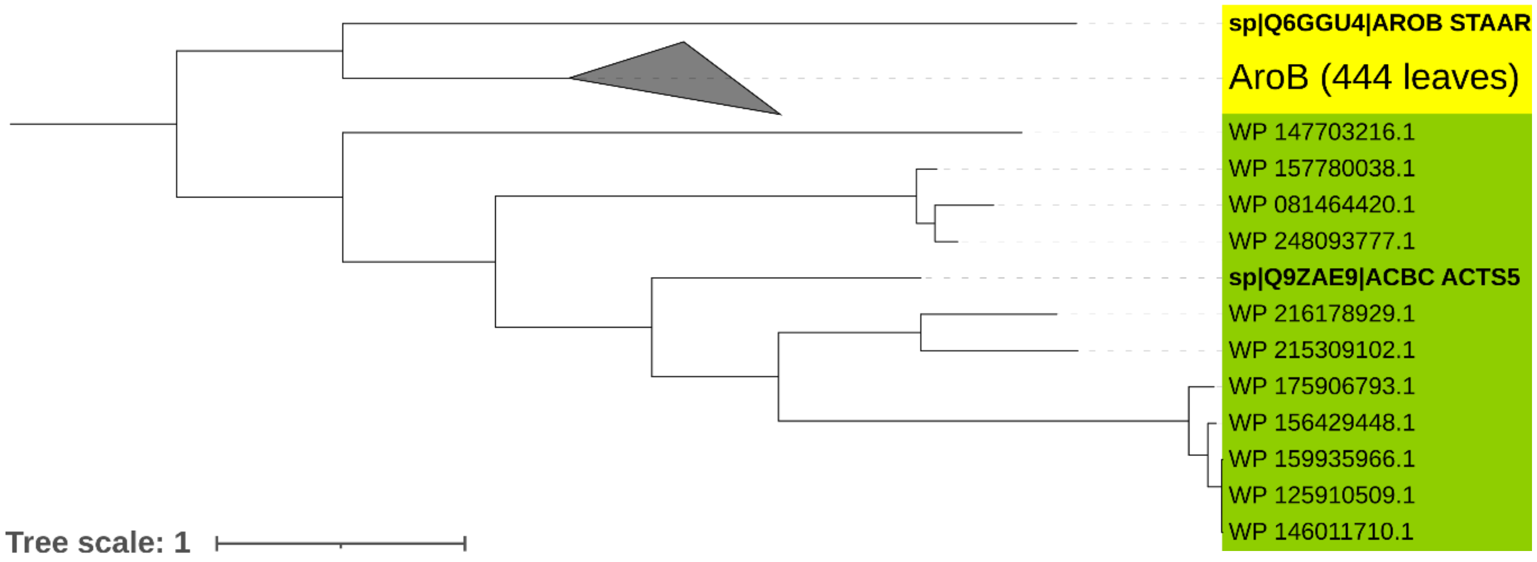


**Figure S7: Phylogenetic analysis of putative EEVS in the *Burkholderiaceae***. The amino acid sequence of the 2-epi-5-epi-valiolone synthase from Actinoplanes sp. ATCC 31044 (accession Q9ZAE9) was retrieved from the UniProt database and used as query in a BlastP search against the NCBI RefSeq database using the NCBI Blast online service with default settings except: the “max target sequences” parameter was increased to 5000; Taxonomic filters were applied to limit hits to the family *Burkholderiaceae* (taxid: 119060); and only hits with e-value < $10^{-3}$ were considered. The searched retrieved 456 matches, which were aligned using MAFFT v7.475[1] in “auto” mode together with the EEVS sequence of Actinoplanes sp. ATCC 31044 (UniProt accession Q9ZAE9) and the DHQS AroB sequence of Staphylococcus aureus (UniProt accession Q6GGU4). Sequences corresponding to leaf nodule *Burkholderia* were manually removed from the alignment. Maximum-likelihood phylogenetic analysis was performed using IQ-TREE v.2.0.3[2] with the “JTT+R6” model and visualized in iTOL[3]. Tree labels correspond to NCBI RefSeq or UniProt accession numbers, with labels in bold indicating reference EEVS and AroB sequences. The green-colored range indicates putative EEVS samples, and the yellow-colored range indicated putative DHQS (AroB) sequences.

References

1. Katoh K, Standley DM. 2013. MAFFT multiple sequence alignment software version 7: improvements in performance and usability. Mol Biol Evol 30:772–80.

2. Letunic I, Bork P. 2016. Interactive tree of life (iTOL) v3: an online tool for the display and annotation of phylogenetic and other trees. Nucleic Acids Res 44:W242–W245.

3. Minh BQ, Schmidt HA, Chernomor O, Schrempf D, Woodhams MD, Von Haeseler A, Lanfear R, Teeling E. 2020. IQ-TREE 2: New Models and Efficient Methods for Phylogenetic Inference in the Genomic Era. Mol Biol Evol 37:1530–1534.
