## Supplementary tables for "Cyclitol secondary metabolism is a central feature of *Burkholderia* leaf symbionts"

**Table S1: Plant samples investigated for leaf endophytes**. Closest relatives were determined using the EzBiocloud identification service (<https://www.ezbiocloud.net/identify>; accessed March 2022). Species marked with * are likely misidentified. ^a^ Estimation for *F. homblei* made by shotgun sequencing of whole leaf tissue ^b^ Presence of *Burkholderia* s.l. symbiont unclear (see results). ^c^ based on only 808bp of 16S rRNA. Ratio bacteria/plant and *Burkholderiaceae*/total bacteria determined by blastn identification of a subset of 1 million reads against the NCBI nucleotide database. Endophyte isolation: +: endophyte isolated successfully; -: endophyte isolation failed; NA: endophyte isolation not attempted

| Species | Voucher | Collection Location | Investigated material | Specialised endophyte structure | Ratio bacteria/plant | Ratio *Burkholderiaceae*/  total bacteria | Endophyte  detected | Endophyte isolation | Closest relative  (16S rRNA; %ID) |
| --- | --- | --- | --- | --- | --- | --- | --- | --- | --- |
| *Ardisia cornudentata* | GU 19862434 | Ghent University Botanical Garden, Belgium | Fresh leaves | Leaf galls | 14.30 | 95.88 | + | NA | *Caballeronia udeis* LMG 27134  (98.48%) |
| *Ardisia mamillata* | GU 19730151 | Ghent University Botanical Garden, Belgium | Fresh leaves | Leaf galls | 1.23 | 86.01 | + | NA | *Caballeronia choica* LMG 22940  (98.83%) |
| *Ardisia virens* | ED 200420025 | Royal Botanic Garden Edinburgh, United Kingdom | Silica-dried leaves | Leaf galls | 0.62 | 94.49 | + | NA | *Caballeronia udeis* LMG 27134  (98.62%) |
| *Fadogia homblei* | PRU 128010 | Roodeplaat, South Africa | Bacterial isolate  (from leaf tissue) | - | 2.84^a^ | 99.15^a^ | + | + | *Paraburkholderia*  *strydomiana* Wk1.1f (100%) |
| *Pavetta capensis** | ED 19697408 | Royal Botanic Garden Edinburgh, United Kingdom | Silica-dried leaves | - | 0.09 | 0.36 | - | NA | NA |
| *Pavetta hochstetteri* | ED 19671929 | Royal Botanic Garden Edinburgh, United Kingdom | Silica-dried leaves | Leaf galls | 18.60 | 97.70 | + | NA | *Caballeronia*  *ptereochthonis* LMG 29326 (97.52%) |
| *Pavetta indica** | BR 2014024336 | Meise Botanic Garden, Belgium | Silica-dried leaves | - | 0.39 | 0.75 | - | NA | NA |
| *Pavetta revoluta* | NA | The Manie van der Schijff Botanical Garden,  Pretoria, South Africa | Fresh leaves | Leaf galls | 0.04 | 69.28 | (+)^b^ | *-* | *Caballeronia calidae* LMG 29321  (98.89%)^c^ |
| *Pavetta ternifolia* | BR 2002160529 | Meise Botanic Garden, Belgium | Silica-dried leaves | Leaf galls | 0.07 | 0.70 | - | NA | NA |
| *Psychotria capensis* | NA | The Manie van der Schijff Botanical Garden,  Pretoria, South Africa | Fresh leaves | Leaf galls | 0.06 | 1.82 | - | - | NA |
| *Psychotria zombamontana* | NA | The Manie van der Schijff Botanical Garden,  Pretoria, South Africa | Fresh leaves | - | 0.00 | 3.92 | - | - | NA |
| *Vangueria dryadum* | PRU 128005 | Lowveld National Botanic Gardens, South Africa | Fresh leaves | - | 1.07 | 97.13 | + | *-* | *Paraburkholderia*  *insulsa* PNG-April (97.65%) |
| *Vangueria esculenta* | BR 2012116870 | Meise Botanic Garden, Belgium | Silica-dried leaves | - | 4.60 | 98.58 | + | *-* | *Paraburkholderia*  *phenoliruptrix* AC1100 (99.59%) |
| *Vangueria infausta* | PRU 126086 | Voortrekker Monument, Pretoria, South Africa | Fresh leaves | - | 0.06 | 85.15 | + | *-* | *Paraburkholderia*  *phenoliruptrix* AC1100 (99.66%) |
| *Vangueria infausta* | PRU 128003 | Lowveld National Botanic Gardens, South Africa | Fresh leaves | - | 9.85 | 98.76 | + | *-* | *Paraburkholderia*  *phenoliruptrix* AC1100 (99.66%) |
| *Vangueria macrocalyx* | PRU 128006 | Lowveld National Botanic Gardens, South Africa | Fresh leaves | - | 1.67 | 97.78 | + | *-* | *Paraburkholderia*  *insulsa* PNG-April (97.65%) |
| *Vangueria madagascariensis* | PRU 128004 | Lowveld National Botanic Gardens, South Africa | Fresh leaves | - | 2.39 | 99.30 | + | *-* | *Paraburkholderia*  *phenoliruptrix* AC1100 (99.64%) |
| *Vangueria madagascariensis* | ED 19715346 | Royal Botanic Garden Edinburgh, United Kingdom | Silica-dried leaves | - | 2.73 | 98.40 | + | *-* | *Paraburkholderia*  *phenoliruptrix* AC1100 (99.66%) |
| *Vangueria madagascariensis* | BR 19584419 | Meise Botanic Garden, Belgium | Silica-dried leaves | - | 3.29 | 98.11 | + | *-* | *Paraburkholderia*  *phenoliruptrix* AC1100 (99.66%) |
| *Vangueria pygmaea* | PRU 126008 | Cullinan, South Africa | Fresh leaves | - | 0.57 | 99.39 | + | *-* | *Paraburkholderia*  *strydomiana* Wk1.1f (100%) |
| *Vangueria pygmaea* | PRU 126088 | Cullinan, South Africa | Fresh leaves | - | 1.63 | 96.97 | + | *-* | *Paraburkholderia*  *strydomiana* Wk1.1f (100%) |
| *Vangueria randii* | PRU 128008 | Lowveld National Botanic Gardens, South Africa | Fresh leaves | - | 1.74 | 98.24 | + | *-* | *Paraburkholderia*  *phenoliruptrix* AC1100 (99.52%) |
| *Vangueria soutpansbergensis* | PRU 128002 | Lowveld National Botanic Gardens, South Africa | Fresh leaves | - | 2.73 | 99.35 | + | *-* | *Paraburkholderia*  *phenoliruptrix* AC1100 (99.45%) |

| Species | Voucher | Collection Location | Investigated material | Specialised endophyte structure | Reference | NCBI Accession |
| --- | --- | --- | --- | --- | --- | --- |
| *Ardisia crenata* | BR 19073685 | Meise Botanic Garden, Belgium | Dissected leaf galls from fresh leaves | Leaf galls | Carlier et al., 2015 | PRJNA253365 |
| *Fadogia homblei* | NA | Meise Botanic Garden, Belgium  (Grown from seeds originally collected in South Africa) | Bacterial isolate R-49542  (isolated from leaves) | - | Verstraete et al., 2014 | Not previously assembled |
| *Pavetta schumanniana* | BR 2000194257 | Meise Botanic Garden, Belgium | Dissected leaf galls from silica-dried leaves | Leaf galls | Pinto-Carbó et al., 2016 | PRJNA253363 |
| *Psychotria brachyanthoides* | BR 2009844596 | Meise Botanic Garden, Belgium | Dissected leaf galls from silica-dried leaves | Leaf galls | Pinto-Carbó et al., 2016 | PRJNA253362 |
| *Psychotria humilis* | BR 2009135940 | Meise Botanic Garden, Belgium | Dissected leaf galls from silica-dried leaves | Leaf galls | Pinto-Carbó et al., 2016 | PRJNA253360 |
| *Psychotria pumila* | BR 2004143571 | Meise Botanic Garden, Belgium | Dissected leaf galls from silica-dried leaves | Leaf galls | Pinto-Carbó et al., 2016 | PRJNA253357 |
| *Psychotria umbellata* | BR 2007130262 | Meise Botanic Garden, Belgium | Dissected leaf galls from silica-dried leaves | Leaf galls | Pinto-Carbó et al., 2016 | PRJNA253361 |
| *Psychotria verschuerenii* | BR 19750204 | Meise Botanic Garden, Belgium | Dissected leaf galls from silica-dried leaves | Leaf galls | Pinto-Carbó et al., 2016 | PRJNA253359 |
| *Psychotria kirkii** | NA | University of Zurich Botanic Garden, Switzerland | Dissected leaf galls from fresh leaves | Leaf galls | Carlier & Eberl, 2012;  Carlier et al., 2015 | GCF_000234195.1 |
| *Psychotria punctata** | NA | University of Zurich Botanic Garden, Switzerland | Dissected leaf galls from fresh leaves | Leaf galls | Pinto-Carbó et al., 2016 | GCF_001189345.1 |

**Table S2: Sample details and accessions numbers for re-assembled and reference genomes.** Samples marked with * are references that were not re-assembled.

**Table S3: *Burkholderia, Caballeronia, and Paraburkholderia* reference genomes used for comparative analysis.**

| **Species** | **NCBI Accession** |
| --- | --- |
| *Burkholderia thailandensis* | GCF_000012365 |
| *Burkholderia lata* | GCF_000012945 |
| *Burkholderia pseudomallei* | GCF_000756125 |
| *Burkholderia plantarii* | GCF_000835205 |
| *Burkholderia oklahomensis* | GCF_000959365 |
| *Burkholderia dolosa* | GCF_000959505 |
| *Burkholderia multivorans* | GCF_000959525 |
| *Burkholderia pyrrocinia* | GCF_001028665 |
| *Burkholderia humptydooensis* | GCF_001462435 |
| *Burkholderia singularis* | GCF_001523725 |
| *Burkholderia vietnamiensis* | GCF_001523785 |
| *Burkholderia territorii* | GCF_001527205 |
| *Burkholderia seminalis* | GCF_001718535 |
| *Burkholderia metallica* | GCF_001718555 |
| *Burkholderia ubonensis* | GCF_001718655 |
| *Burkholderia cenocepacia* | GCF_001718895 |
| *Burkholderia stagnalis* | GCF_001718955 |
| *Burkholderia mallei* | GCF_002346025 |
| *Burkholderia reimsis* | GCF_003294055 |
| *Burkholderia contaminans* | GCF_004723625 |
| *Burkholderia cepacia* | GCF_009586235 |
| *Burkholderia glumae* | GCF_009931375 |
| *Burkholderia guangdongensis* | GCF_013403875 |
| *Burkholderia stabilis* | GCF_900240005 |
| *Burkholderia gladioli* | GCF_900608535 |
| *Burkholderia anthina* | GCF_902498995 |
| *Burkholderia paludis* | GCF_902499105 |
| *Burkholderia arboris* | GCF_902499125 |
| *Burkholderia aenigmatica* | GCF_902499295 |
| *Burkholderia ambifaria* | GCF_902829835 |
| *Burkholderia diffusa* | GCF_902830815 |
| *Burkholderia latens* | GCF_902832795 |
| *Burkholderia pseudomultivorans* | GCF_902832925 |
| *Burkholderia puraquae* | GCF_902859845 |
| *Caballeronia insecticola* | GCF_000402035 |
| *Caballeronia grimmiae* | GCF_000698555 |
| *Caballeronia mineralivorans* | GCF_001028175 |
| *Caballeronia sordidicola* | GCF_001544455 |
| *Caballeronia humi* | GCF_001544475 |
| *Caballeronia telluris* | GCF_001544495 |
| *Caballeronia terrestris* | GCF_001544515 |
| *Caballeronia choica* | GCF_001544535 |
| *Caballeronia udeis* | GCF_001544555 |
| *Caballeronia cordobensis* | GCF_001544575 |
| *Caballeronia concitans* | GCF_001544615 |
| *Caballeronia arvi* | GCF_001544695 |
| *Caballeronia catudaia* | GCF_001544755 |
| *Caballeronia temeraria* | GCF_001544795 |
| *Caballeronia fortuita* | GCF_001544835 |
| *Caballeronia hypogeia* | GCF_001544875 |
| *Caballeronia pedi* | GCF_001544915 |
| *Caballeronia glebae* | GCF_001545035 |
| *Caballeronia ptereochthonis* | GCF_001545075 |
| *Caballeronia calidae* | GCF_900044055 |
| *Caballeronia arationis* | GCF_900230245 |
| *Caballeronia novacaledonica* | GCF_900258035 |
| *Caballeronia glathei* | GCF_902833485 |
| *Caballeronia zhejiangensis* | GCF_902833575 |
| *Paraburkholderia xenovorans* | GCF_000013645 |
| *Paraburkholderia phymatum* | GCF_000020045 |
| *Paraburkholderia phytofirmans* | GCF_000020125 |
| *Paraburkholderia kururiensis* | GCF_000341045 |
| *Paraburkholderia dilworthii* | GCF_000472525 |
| *Paraburkholderia mimosarum* | GCF_000472825 |
| *Paraburkholderia nodosa* | GCF_000519185 |
| *Paraburkholderia acidipaludis* | GCF_000684975 |
| *Paraburkholderia bannensis* | GCF_000685015 |
| *Paraburkholderia ferrariae* | GCF_000685035 |
| *Paraburkholderia oxyphila* | GCF_000685075 |
| *Paraburkholderia heleia* | GCF_000739775 |
| *Paraburkholderia sacchari* | GCF_000785435 |
| *Paraburkholderia monticola* | GCF_001580545 |
| *Paraburkholderia ginsengiterrae* | GCF_001645135 |
| *Paraburkholderia sprentiae* | GCF_001865575 |
| *Paraburkholderia acidophila* | GCF_002097715 |
| *Paraburkholderia aromaticivorans* | GCF_002278075 |
| *Paraburkholderia fungorum* | GCF_002891075 |
| *Paraburkholderia terrae* | GCF_002902925 |
| *Paraburkholderia insulsa* | GCF_003002115 |
| *Paraburkholderia eburnea* | GCF_003003375 |
| *Paraburkholderia unamae* | GCF_003096875 |
| *Paraburkholderia silvatlantica* | GCF_003217075 |
| *Paraburkholderia bryophila* | GCF_003269035 |
| *Paraburkholderia dokdonella* | GCF_003286395 |
| *Paraburkholderia graminis* | GCF_003330785 |
| *Paraburkholderia terricola* | GCF_003330825 |
| *Paraburkholderia lacunae* | GCF_003353175 |
| *Paraburkholderia caffeinilytica* | GCF_003368325 |
| *Paraburkholderia phosphatilytica* | GCF_003443895 |
| *Paraburkholderia dinghuensis* | GCF_003837865 |
| *Paraburkholderia guartelaensis* | GCF_004353905 |
| *Paraburkholderia rhizosphaerae* | GCF_004366595 |
| *Paraburkholderia dipogonis* | GCF_004402975 |
| *Paraburkholderia azotifigens* | GCF_007995085 |
| *Paraburkholderia panacisoli* | GCF_008369935 |
| *Paraburkholderia franconis* | GCF_009362735 |
| *Paraburkholderia bonniea* | GCF_009455625 |
| *Paraburkholderia agricolaris* | GCF_009455635 |
| *Paraburkholderia hayleyella* | GCF_009455685 |
| *Paraburkholderia madseniana* | GCF_009690905 |
| *Paraburkholderia acidiphila* | GCF_009789655 |
| *Paraburkholderia acidisoli* | GCF_009789675 |
| *Paraburkholderia youngii* | GCF_013366925 |
| *Paraburkholderia caribensis* | GCF_013378095 |
| *Paraburkholderia tropica* | GCF_014171495 |
| *Paraburkholderia atlantica* | GCF_014200895 |
| *Paraburkholderia ginsengisoli* | GCF_016128195 |
| *Paraburkholderia caledonica* | GCF_902833635 |
| *Paraburkholderia piptadeniae* | GCF_900007165 |
| *Paraburkholderia ribeironis* | GCF_900019265 |
| *Paraburkholderia lycopersici* | GCF_900096975 |
| *Paraburkholderia phenazinium* | GCF_900100735 |
| *Paraburkholderia tuberum* | GCF_900101795 |
| *Paraburkholderia caballeronis* | GCF_900104845 |
| *Paraburkholderia sartisoli* | GCF_900107685 |
| *Paraburkholderia diazotrophica* | GCF_900108945 |
| *Paraburkholderia megapolitana* | GCF_900113825 |
| *Paraburkholderia aspalathi* | GCF_900116445 |
| *Paraburkholderia hospita* | GCF_900167965 |
| *Paraburkholderia susongensis* | GCF_900177725 |
| *Paraburkholderia rhynchosiae* | GCF_902859775 |
| *Paraburkholderia sediminicola* | GCF_902859805 |
| *Paraburkholderia phenoliruptrix* | GCF_902859825 |
| *Paraburkholderia humisilvae* | GCF_902859855 |
| *Paraburkholderia solisilvae* | GCF_902859875 |
| *Paraburkholderia ultramafica* | GCF_902859915 |
| *Paraburkholderia fynbosensis* | GCF_902859935 |
| *Paraburkholderia caffeinitolerans* | GCF_902859945 |
| *Paraburkholderia kirstenboschensis* | GCF_904848585 |
| *Paraburkholderia metrosideri* | GCF_904848625 |
| *Paraburkholderia sabiae* | GCF_904848645 |
| *Paraburkholderia hiiakae* | GCF_904848665 |
| *Paraburkholderia strydomiana* | GCF_004334935 |

**Table S4: Average nucleotide identities between leaf endophyte genomes.** ANI values are based on blastn alignments. Genome abbreviations refer to the species of host plant (e.g. Acor = endophyte of *Ardisia cornudentata*)

|  | **Acor** | **Acre** | **Amam** | **Avir** | **FhomR** | **FhomSA** | **Pbra** | **Phoc** | **Phum** | **Pkir** | **Ppum** | **Ppun** | **Psch** | **Pumb** | **Pver** | **Vdry** | **Vesc** | **Vinf** | **Vmac** | **Vmad** | **VmadE** | **VmadM** | **Vpyg08** | **Vpyg88** | **Vran** | **Vsou** |
| --- | --- | --- | --- | --- | --- | --- | --- | --- | --- | --- | --- | --- | --- | --- | --- | --- | --- | --- | --- | --- | --- | --- | --- | --- | --- | --- |
| **Acor** | 1.00 | 0.94 | 0.96 | 0.94 | 0.75 | 0.75 | 0.77 | 0.77 | 0.77 | 0.77 | 0.77 | 0.78 | 0.77 | 0.76 | 0.77 | 0.76 | 0.76 | 0.76 | 0.76 | 0.76 | 0.76 | 0.76 | 0.75 | 0.75 | 0.76 | 0.76 |
| **Acre** | 0.94 | 1.00 | 0.94 | 0.99 | 0.76 | 0.76 | 0.78 | 0.78 | 0.78 | 0.78 | 0.78 | 0.78 | 0.78 | 0.77 | 0.78 | 0.77 | 0.77 | 0.77 | 0.77 | 0.77 | 0.77 | 0.77 | 0.76 | 0.76 | 0.77 | 0.77 |
| **Amam** | 0.96 | 0.94 | 1.00 | 0.94 | 0.75 | 0.75 | 0.77 | 0.77 | 0.76 | 0.77 | 0.77 | 0.77 | 0.77 | 0.76 | 0.77 | 0.76 | 0.76 | 0.76 | 0.76 | 0.76 | 0.76 | 0.76 | 0.75 | 0.75 | 0.76 | 0.76 |
| **AvirE** | 0.94 | 0.99 | 0.94 | 1.00 | 0.76 | 0.76 | 0.78 | 0.78 | 0.78 | 0.78 | 0.78 | 0.78 | 0.78 | 0.77 | 0.78 | 0.77 | 0.77 | 0.77 | 0.77 | 0.77 | 0.77 | 0.77 | 0.76 | 0.76 | 0.77 | 0.77 |
| **FhomR** | 0.76 | 0.77 | 0.75 | 0.77 | 1.00 | 0.98 | 0.77 | 0.78 | 0.76 | 0.77 | 0.77 | 0.77 | 0.78 | 0.77 | 0.76 | 0.82 | 0.86 | 0.86 | 0.82 | 0.86 | 0.86 | 0.86 | 0.98 | 0.98 | 0.86 | 0.86 |
| **FhomSA** | 0.76 | 0.77 | 0.75 | 0.77 | 0.98 | 1.00 | 0.77 | 0.78 | 0.76 | 0.77 | 0.77 | 0.77 | 0.78 | 0.77 | 0.76 | 0.82 | 0.86 | 0.86 | 0.82 | 0.86 | 0.86 | 0.86 | 0.98 | 0.98 | 0.86 | 0.85 |
| **Pbra** | 0.77 | 0.78 | 0.77 | 0.78 | 0.76 | 0.77 | 1.00 | 0.83 | 0.82 | 0.83 | 0.82 | 0.83 | 0.83 | 0.91 | 0.86 | 0.78 | 0.78 | 0.78 | 0.78 | 0.78 | 0.78 | 0.78 | 0.77 | 0.77 | 0.78 | 0.78 |
| **Phoc** | 0.77 | 0.78 | 0.77 | 0.78 | 0.77 | 0.77 | 0.82 | 1.00 | 0.82 | 0.94 | 0.82 | 0.87 | 0.95 | 0.82 | 0.83 | 0.78 | 0.78 | 0.78 | 0.78 | 0.78 | 0.78 | 0.78 | 0.77 | 0.77 | 0.78 | 0.78 |
| **Phum** | 0.77 | 0.78 | 0.77 | 0.78 | 0.76 | 0.76 | 0.83 | 0.82 | 1.00 | 0.82 | 0.82 | 0.83 | 0.83 | 0.82 | 0.83 | 0.77 | 0.77 | 0.77 | 0.77 | 0.77 | 0.77 | 0.77 | 0.76 | 0.76 | 0.77 | 0.77 |
| **Pkir** | 0.77 | 0.78 | 0.77 | 0.78 | 0.76 | 0.76 | 0.82 | 0.94 | 0.82 | 1.00 | 0.82 | 0.87 | 0.96 | 0.82 | 0.82 | 0.78 | 0.78 | 0.78 | 0.78 | 0.78 | 0.78 | 0.78 | 0.77 | 0.77 | 0.78 | 0.78 |
| **Ppum** | 0.77 | 0.78 | 0.77 | 0.78 | 0.76 | 0.76 | 0.82 | 0.82 | 0.82 | 0.82 | 1.00 | 0.82 | 0.82 | 0.82 | 0.82 | 0.77 | 0.77 | 0.77 | 0.77 | 0.77 | 0.77 | 0.77 | 0.76 | 0.76 | 0.77 | 0.77 |
| **Ppun** | 0.77 | 0.78 | 0.77 | 0.78 | 0.77 | 0.77 | 0.83 | 0.87 | 0.82 | 0.87 | 0.82 | 1.00 | 0.88 | 0.82 | 0.83 | 0.78 | 0.79 | 0.79 | 0.78 | 0.79 | 0.79 | 0.79 | 0.77 | 0.77 | 0.79 | 0.79 |
| **Psch** | 0.77 | 0.78 | 0.77 | 0.78 | 0.77 | 0.77 | 0.83 | 0.95 | 0.82 | 0.96 | 0.82 | 0.88 | 1.00 | 0.82 | 0.83 | 0.78 | 0.78 | 0.78 | 0.78 | 0.78 | 0.78 | 0.78 | 0.77 | 0.77 | 0.78 | 0.78 |
| **Pumb** | 0.76 | 0.78 | 0.76 | 0.78 | 0.76 | 0.76 | 0.91 | 0.82 | 0.82 | 0.82 | 0.82 | 0.83 | 0.82 | 1.00 | 0.85 | 0.77 | 0.77 | 0.77 | 0.77 | 0.77 | 0.77 | 0.77 | 0.76 | 0.76 | 0.77 | 0.77 |
| **Pver** | 0.77 | 0.78 | 0.77 | 0.78 | 0.76 | 0.76 | 0.86 | 0.83 | 0.83 | 0.83 | 0.82 | 0.83 | 0.83 | 0.86 | 1.00 | 0.77 | 0.78 | 0.78 | 0.77 | 0.78 | 0.78 | 0.78 | 0.76 | 0.76 | 0.78 | 0.78 |
| **Vdry** | 0.76 | 0.77 | 0.76 | 0.77 | 0.82 | 0.82 | 0.78 | 0.78 | 0.77 | 0.78 | 0.77 | 0.78 | 0.78 | 0.77 | 0.77 | 1.00 | 0.83 | 0.83 | 1.00 | 0.83 | 0.83 | 0.83 | 0.82 | 0.82 | 0.83 | 0.83 |
| **Vesc** | 0.76 | 0.77 | 0.76 | 0.77 | 0.85 | 0.85 | 0.78 | 0.79 | 0.77 | 0.78 | 0.77 | 0.78 | 0.79 | 0.77 | 0.78 | 0.83 | 1.00 | 1.00 | 0.83 | 1.00 | 0.99 | 0.99 | 0.85 | 0.85 | 1.00 | 0.95 |
| **Vinf** | 0.76 | 0.77 | 0.76 | 0.77 | 0.85 | 0.85 | 0.78 | 0.79 | 0.77 | 0.78 | 0.77 | 0.78 | 0.79 | 0.77 | 0.77 | 0.83 | 1.00 | 1.00 | 0.83 | 1.00 | 0.99 | 0.99 | 0.85 | 0.85 | 1.00 | 0.95 |
| **Vmac** | 0.76 | 0.77 | 0.76 | 0.77 | 0.82 | 0.82 | 0.78 | 0.78 | 0.77 | 0.78 | 0.77 | 0.78 | 0.78 | 0.77 | 0.77 | 1.00 | 0.83 | 0.83 | 1.00 | 0.83 | 0.83 | 0.83 | 0.82 | 0.82 | 0.83 | 0.83 |
| **Vmad** | 0.76 | 0.77 | 0.76 | 0.77 | 0.85 | 0.85 | 0.78 | 0.78 | 0.77 | 0.78 | 0.77 | 0.78 | 0.79 | 0.77 | 0.77 | 0.83 | 1.00 | 1.00 | 0.83 | 1.00 | 0.99 | 0.99 | 0.85 | 0.85 | 1.00 | 0.95 |
| **VmadE** | 0.76 | 0.77 | 0.76 | 0.77 | 0.85 | 0.85 | 0.78 | 0.79 | 0.77 | 0.78 | 0.77 | 0.79 | 0.79 | 0.78 | 0.78 | 0.83 | 0.99 | 0.99 | 0.83 | 0.99 | 1.00 | 1.00 | 0.85 | 0.85 | 0.99 | 0.95 |
| **VmadM** | 0.76 | 0.77 | 0.76 | 0.77 | 0.85 | 0.85 | 0.78 | 0.79 | 0.77 | 0.78 | 0.77 | 0.79 | 0.79 | 0.78 | 0.78 | 0.83 | 0.99 | 0.99 | 0.83 | 0.99 | 1.00 | 1.00 | 0.85 | 0.85 | 0.99 | 0.95 |
| **Vpyg08** | 0.76 | 0.77 | 0.75 | 0.77 | 0.97 | 0.97 | 0.77 | 0.78 | 0.76 | 0.78 | 0.77 | 0.78 | 0.78 | 0.77 | 0.77 | 0.82 | 0.86 | 0.86 | 0.82 | 0.86 | 0.86 | 0.86 | 1.00 | 1.00 | 0.86 | 0.86 |
| **Vpyg88** | 0.76 | 0.77 | 0.75 | 0.77 | 0.97 | 0.97 | 0.77 | 0.78 | 0.76 | 0.78 | 0.77 | 0.78 | 0.78 | 0.77 | 0.77 | 0.82 | 0.86 | 0.86 | 0.82 | 0.86 | 0.86 | 0.86 | 1.00 | 1.00 | 0.86 | 0.86 |
| **Vran** | 0.76 | 0.77 | 0.76 | 0.78 | 0.85 | 0.85 | 0.78 | 0.79 | 0.77 | 0.78 | 0.78 | 0.78 | 0.79 | 0.77 | 0.77 | 0.83 | 1.00 | 1.00 | 0.83 | 1.00 | 0.99 | 0.99 | 0.85 | 0.85 | 1.00 | 0.95 |
| **Vsou** | 0.76 | 0.77 | 0.76 | 0.77 | 0.85 | 0.85 | 0.78 | 0.78 | 0.77 | 0.78 | 0.78 | 0.79 | 0.78 | 0.78 | 0.77 | 0.83 | 0.95 | 0.95 | 0.83 | 0.95 | 0.95 | 0.95 | 0.85 | 0.85 | 0.95 | 1.00 |

**Table S5: Non-essential core genes of leaf endophytes.** Gene identifiers are from the genome of *Ca.* Burkholderia kirkii (NCBI accession GCF_000234195.1). Abbreviations: COG – Cluster of Orthologues Genes. COG Category meanings: D – Cell cycle control, cell division, chromosome portioning; E – Amino acid transport and metabolism; F – Nucleotide metabolism and transport; G – Carbohydrate transport and metabolism; H – Coenzyme transport and metabolism; L – Replication, recombination and repair; M – Cell wall/membrane/envelope biogenesis; O – Post-translational modification, protein turnover, and chaperones; P – Inorganic ion transport and metabolism; Q – Secondary metabolite biosynthesis, transport and catabolism; S – Function unknown; T – Signal transduction mechanisms.

| Gene | COG category | Functional prediction (EGGNOG) | Best blastp hit in NCBI nr protein database | Best blastp hit in UniPept/SwissProt |
| --- | --- | --- | --- | --- |
| CCD_35550.1 | H | Removes the pyruvyl group from chorismate, with concomitant aromatization of the ring, to provide 4- hydroxybenzoate (4HB) for the ubiquinone pathway | Chorismate lyase (*Caballeronia* *ptereochtonis*) | Probable chorismate pyruvate-lyase  (*Paraburkholderia* *xenovorans*) |
| CCD_35680.1 | M | Membrane protein | Colicin transporter (*Caballeronia* *ptereochtonis*) | No hit |
| CCD_36330.1 | M | Membrane protein | Phage holin family protein (*Caballeronia* *novacaledonica*) | Uncharacterized membrane protein YvID (*Bacillus* *subtilis*) |
| CCD_36810.1 | Q | Carboxymethylenebutenolidase | Dienelactone hydrolase family protein  (*Caballeronia* *calidae*) | Putative carboxymethylenebutenolidase/Dienelactone hydrolase (*Azospirillum* *brasilense*) |
| CCD_37159.1 | M | (Lipo)protein | Outer membrane protein assembly factor BamC (*Caballeronia* *catudaia*) | Outer membrane protein assembly factor BamC (*Thiobacillus* *denitrificans*) |
| CCD_37257.1 | D | Cell division protein ZapD | Cell division protein ZapD (*Caballeronia*) | Cell division protein ZapD (*Burkholderia* *lata*) |
| CCD_37658.1 | S | Trm112 family protein | Acyl-ACP desaturase (*Caballeronia* *pedi*) | No hit |
| CCD_37721.1 | S | Protein of unknown function (DUF2909) | Twin transmembrane helix small protein (*Burkholderiaceae*) | No hit |
| CCD_37723.1 | S | Signal sequence binding sco1 | Cytochrome C oxidase subunit I (*Caballeronia* *glebae*) | No hit |
| CCD_37888.1 | S | Bacterial protein of unknown function (DUF883) | DUF883 family protein (*Caballeronia* *ptereochtonis*) | Uncharacterized protein YgjD (*Escherichia* *coli*) |
| CCD_37889.1 | S | Membrane protein | Phage holin family protein (*Caballeronia* *ptereochtonis*) | No hit |
| CCD_37890.1 | S | Protein of unknown function (DUF3318) | DUF3318 domain-containing protein (Caballeronia *calidae*) | No hit |
| CCD_37930.1 | F | Phosphoribosylaminoimidazolesuccinocarboxamide synthase purC | Phosphoribosylaminoimidazolesuccinocarboxamide synthase (*Caballeronia* *megalochromosomata*) | Phosphoribosylaminoimidazole-succinocarboxamide synthase (*Cupriavidus* *metalluridans*) |
| CCD_37995.2 | O | Peptide-methioine (S)-S-oxide reductase MsrA | Peptide-methioine (S)-S-oxide reductase MsrA  (*Caballeronia* *calidae*) | Peptide methionine sulfoxide reductase MsrA  (*Ralstonia* *solanacearum*) |
| CCD_38181.1 | P | Catalase activity | Ferritin-like domain-containing protein  (*Caballeronia* *temeraria*) | No hit |
| CCD_38266.1 | GM | Nad-dependent epimerase dehydratase | SDR family oxidoreductase (*Caballeronia* *ptereochtonis*) | UDP-glucose 4-epimerase (*Vibrio* *cholerae*) |
| CCD_38823.1 | S | STAS domain-containing protein | anti-anti-sigma regulatory factor (*Caballeronia* *jiangsuensis*) | No hit |
| CCD_39033.1 | S | Protein of unknown function (DUF2863) | DUF2863 family protein (*Caballeronia* *ptereochtonis*) | No hit |
| CCD_39432.1 | L | Involved in DNA repair and RecF pathway recombination | DNA repair protein RecO (*Caballeronia* *jiangsuensis*) | DNA repair protein RecO (*Paraburkholderia* *xenovorans*) |
| CCD_39439.1 | T | Regulatory protein | MucB/RseB C-terminal domain-containing protein (*Caballeronia* *fortuita*) | Sigma-E factor regulatory protein RseB  (*Haemophilus* *influenzae*) |
| CCD_39635.1 | S | Sterol-binding domain protein | Sterol-binding protein (*Caballeronia* *peredens*) | No hit |
| CCD_39933.1 | S | Hypothetical protein | Regulator (*Caballeronia* *ptereochtonis*) | No hit |
| CCD_39967.1 | M | (Lipo)protein | Outer membrane protein assembly factor BamC (*Caballeronia* *catudaia*) | Outer membrane protein assembly factor BamC (*Thiobacillus* *denitrificans*) |
| CCD_40265.1 | H | ATP-dependent carboxylate-amine ligase which exhibits weak glutamate--cysteine ligase activity | Carboxylate-amine ligase (*Caballeronia* *turbans*) | Putative glutamate-cysteine ligase 2  (*Paraburkholderia* *phytofirmans*) |
| CCD_40266.1 | P | Sodium:hydrogen antiporter | Sodium/hydrogen exchanger (*Caballeronia* *peredens*) | No hit |
| CCD_40282.1 | E | Arginine/lysine/ornithine decarboxylase | Arginine/lysine/ornithine decarboxylase  (*Caballeronia* *ptereochtonis*) | Lysine decarboxylase LdcA (*Pseudomonas* *aeruginosa*) |
| CCD_40366.1 | O | Arginyl-tRNA-protein transferase | Arginyl-tRNA-protein transferase (*Caballeronia* *hypogeia*) | Aspartate/glutamate leucyltransferase  (*Paraburkholderia* *phymatum*) |
| CCD_40735.1 | S | Fe-S cluster assembly protein IscX | Fe-S cluster assembly protein IscX (*Burkholderiaceae*) | Protein IscX (*Haemophilus* *influenzae*) |

**Table S6: Leaf endophyte core genes not conserved in *Burkholderia*, *Caballeronia*, and *Paraburkholderia* genomes.** Gene identifiers are from the genome of *Ca.* Burkholderia kirkii (NCBI accession GCF_000234195.1). Abbreviations: COG – Cluster of Orthologues Genes. COG Category meanings: C – Energy production and conversion; E – Amino acid transport and metabolism; F – Nucleotide metabolism and transport; G – Carbohydrate metabolism and transport; H – Coenzyme metabolism and transport; J – Translation, ribosomal structure and biogenesis; K – Transcription; L – Replication, recombination and repair; M – Cell wall/membrane/envelope biogenesis; O – Post-translational modification, protein turnover, and chaperones; P – Inorganic ion transport and metabolism; Q – Secondary metabolite biosynthesis, transport and catabolism; S – Function unknown; T – Signal transduction mechanisms; V – Defence mechanisms.

| Gene | COG category | Functional prediction (EGGNOG) | Best blastp hit in NCBI nr protein database | Best blastp hit in UniPept/SwissProt |
| --- | --- | --- | --- | --- |
| CCD_35375.1 | T | histidine kinase A domain protein | Molecular chaperone DnaK (*Caballeronia* *ptereochtonis*) | Chaperone protein DnaK (*Paraburkholderia* *phymatum*) |
| CCD_36748.1 | O | Binds to Cpn60 in the presence of Mg-ATP and suppresses the ATPase activity of the latter | Co-chaperone GroES (*Burkholderiaceae*) | 10kDa chaperonin/GroES/Cpn10  (*Paraburkholderia* *phytofirmans*) |
| CCD_36810.1 | Q | Carboxymethylenebutenolidase | Dienelactone hydrolase family protein (*Caballeronia* *calidae*) | Putative carboxymethylenebutenolidase/Dienelactone hydrolase (*Azospirillum* *brasilense*) |
| CCD_36831.1 | H | dihydroneopterin aldolase | Dihydroneopterin aldolase (*Caballeronia* *insecticola*) | No hit |
| CCD_37258.2 | S | Protein conserved in bacteria | Dephospho-CoA kinase (*Caballeronia* *ptereochtonis*) | Dephospho-CoA kinase (*Burkholderia* *lata*) |
| CCD_37382.1 | V | PFAM ABC transporter related | ABC transporter (*Caballeronia* *cordobensis*) | Uncharacterized ABC transporter ATP-binding protein YadG (*Escherichia* *coli*) |
| CCD_37823.1 | C | Belongs to the citrate synthase family | Citrate (Si)-synthase (*Caballeronia*) | Citrate synthase (*Bradyrhizobium* *diazoefficiens*) |
| CCD_38121.1 | L | TIGRFAM hydrolase, TatD family | TatD family hydrolase (*Caballeronia* *ptereochtonis*) | Uncharacterized metal-dependent hydrolase  (*Haemophilus* *influenzae*) |
| CCD_38703.1 | K | Belongs to the ParB family | ParB/RepB/Spo0J family partition protein (*Caballeronia* *pedi*) | Probable chromosome-partitioning protein ParB  (*Pseudomonas* *putida*) |
| CCD_38731.1 | M | Belongs to the D-alanine--D-alanine ligase family | D-alanine-D-alanine ligase (*Caballeronia* *novacaledonica*) | D-alanine-D-alanine ligase (*Paraburkholderia* *xenovorans*) |
| CCD_39839.1 | L | Histone-like DNA-binding protein which is capable of wrapping DNA to stabilize it, and thus to prevent its denaturation under extreme environmental conditions | HU family DNA-binding protein (*Caballeronia* *zhejiangensis*) | DNA-binding protein HU-beta (*Pseudomonas* *aeruginosa*) |
| CCD_40423.1 | H | Methyltransferase type 11 | Bifunctional 2-polyprenyl-6-hydroxyphenol methylase/3-demethylubiquinol 3-O-methyltransferase UbiG  (*Caballeronia* *zhejiangensis*) | Ubiquinone biosynthesis O-methyltransferase  (*Paraburkholderia* *phymatum*) |
| CCD_40926.1 | EGP | PFAM major facilitator superfamily MFS_1 | Lysophospholipid transporter LplT (*Caballeronia* *zhejiangensis*) | Lysophospholipid transporter LplT (*Yersinia* *enterocolitica*) |
| CCD_41480.1 | FJ | Catalyses the deamination of adenosine to inosine at the wobble position 34 of tRNA(Arg2) | tRNA adenosine(34) deaminase TadA (*Caballeronia* *insecticola*) | tRNA-specific adenosine deaminase (*Haemophilus* *influenzae*) |

**Table S7: Top 25 orthologues groups of genes enriched in endophytes compared to other members of *Burkholderia, Caballeronia,* and *Paraburkholderia*.** The representative gene of *Ca.* Burkholderia kirkii was chosen as representative if possible, otherwise it was decided randomly. EEVS-cluster denotes if the gene is present in one of the two EEVS (2-*epi*-5-*epi*-valiolone synthase) gene clusters identified in *Ca.* Burkholderia kirkii. Closest relative represent the genus of the closest relative of that orthogroup by blastp searches against the RefSeq protein database (accessed June 2021). Endophyte/BCP genomes and proportion denotes the number and proportion of endophyte/BCP genomes present in the orthogroup. The difference is calculated as Endophyte proportion – BCP proportion. Abbreviations: EEVS - 2-*epi*-5-*epi*-valiolone synthase; BCP – *Burkholderia*/*Caballeronia*/*Paraburkholderia*; COG – Cluster of Orthologues Genes; COG category meaning: C – Energy production and conversion; E – Amino acid metabolism and transport; G – Carbohydrate metabolism and transport; H – Coenzyme metabolism and transport; J – Translation, ribosomal structure and biogenesis; K – Transcription; M – Cell wall/membrane/envelope biogenesis; O – Post-translational modifications, protein turnover, and chaperones; Q – Secondary metabolite metabolism, transport, and catabolism; S – Function unknown.

| Representative gene | EEVS-cluster | | Closest relative (RefSeq) | Endophyte genomes | BCP genomes | Endophyte proportion | BCP proportion | Difference | COG | Functional annotation |
| --- | --- | --- | --- | --- | --- | --- | --- | --- | --- | --- |
| CCD_36718.2 | | X | *Pseudomonas* | 24 | 1 | 0.92 | 0.01 | 0.92 | E | 2-*epi*-5-*epi*-valiolone synthase |
| CCD_39382.1 | |  | *Xenorhabdus* | 21 | 1 | 0.81 | 0.01 | 0.80 | S | Unknown |
| CCD_39384.2 | |  | *Xenorhabdus* | 20 | 0 | 0.77 | 0.00 | 0.77 | J | Radical SAM superfamily |
| CCD_36712.1 | | X | *Pseudomonas* | 17 | 0 | 0.65 | 0.00 | 0.65 | G | Mannose-6-phosphate isomerase, cupin superfamily |
| CCD_36715.1 | | X | *Pseudomonas* | 17 | 0 | 0.65 | 0.00 | 0.65 | G | Glycoside Hydrolases Family 4; Likely 6-phosho-beta-glucosidase |
| CCD_39395.1 | | X | *Noviherbaspirillum* | 15 | 0 | 0.58 | 0.00 | 0.58 | G | Trehalose-6-phosphate synthase |
| CCD_39415.1 | |  | *Burkholderia* | 19 | 27 | 0.73 | 0.20 | 0.53 | S | Unknown |
| CCD_39396.1 | | X | *Noviherbaspirillum* | 13 | 0 | 0.50 | 0.00 | 0.50 | E | Medium chain reductase/dehydrogenase (MDR)/zinc-dependent alcohol dehydrogenase-like family |
| CCD_39398.1 | | X | *Noviherbaspirillum* | 13 | 0 | 0.50 | 0.00 | 0.50 | G | Major Facilitator Superfamily Transporter |
| PCALR49542_2702 | |  | *Paraburkholderia* | 13 | 4 | 0.50 | 0.03 | 0.47 | K | NA-binding transcriptional regulator, IclR family |
| CCD_39397.1 | | X | *Noviherbaspirillum* | 12 | 0 | 0.46 | 0.00 | 0.46 | S | HAD-family hydrolase |
| PPHERAN_6119 | |  | *Pseudomonas* | 12 | 2 | 0.46 | 0.02 | 0.45 | C | Pyruvate-formate lyase-activating enzyme |
| PPHERAN_2366 | |  | *Paraburkholderia* | 13 | 8 | 0.50 | 0.06 | 0.44 | Q | Homospermidine synthase |
| CCD_39196.1 | |  | *Caballeronia* | 22 | 56 | 0.85 | 0.42 | 0.42 | H | NAD-synthase |
| CCD_39408.1 | |  | *Nitrospira* | 10 | 0 | 0.38 | 0.00 | 0.38 | H | N-Acyltransferase superfamily; possibly -acyl-L-homoserine lactone synthetase |
| PCALR49542_2579 | |  | No Hit | 10 | 0 | 0.38 | 0.00 | 0.38 | S | Unknown |
| PPHERAN_5893 | |  | *Breoghania* | 10 | 0 | 0.38 | 0.00 | 0.38 | E | Amidinotransferase; possibly N-Dimethylarginine dimethylaminohydrolase or N-Dimethylarginine dimethylaminohydrolase |
| PPHERAN_5895 | |  | *Breoghania* | 10 | 0 | 0.38 | 0.00 | 0.38 | J | Aspartyl-tRNA synthetase |
| PPHERAN_6120 | |  | *Pseudomonas* | 10 | 0 | 0.38 | 0.00 | 0.38 | S | Unknown |
| PCALR49542_6990 | |  | *Paraburkholderia* | 11 | 8 | 0.42 | 0.06 | 0.36 | O | Glycosyltransferase Family 4 protein |
| PCALR49542_6995 | |  | *Paraburkholderia* | 11 | 8 | 0.42 | 0.06 | 0.36 | M | SAM-dependent methyltransferase |
| CCD_35310.1 | |  | *Caballeronia* | 23 | 70 | 0.88 | 0.53 | 0.35 | S | Bacterial protein of unknown function (DUF883) |
| PPHERAN_5481 | |  | *Paraburkholderia* | 10 | 4 | 0.38 | 0.03 | 0.35 | M | RfaE bifunctional ADP-heptose synthase |
| PCALR49542_6993 | |  | *Paraburkholderia* | 11 | 11 | 0.42 | 0.08 | 0.34 | S | GNAT family N-acetyltransferase |
| PPHERAN_1463 | |  | *Paraburkholderia* | 10 | 6 | 0.38 | 0.05 | 0.34 | S | Unknown |

**Table S8: EEVS cluster organisation of other EEVS-clusters in endophyte genomes.** Genomes of the same host with the same cluster layout are merged. Genera in brackets represents the genus of the closest protein relative. *A large region has three predicted genes in different frames that show homology with EEVS genes, possibly due to one or multiple frameshift mutations. Abbreviations: EEVS – 2-*epi*-5-*epi*-valiolone synthase; IS – Insertion element; Ψ – gene predicted to be a pseudogene.

| *Ca*. Burkholderia verschuerenii | IS5 | EEVS (Pver_5505) (*Pseudomonas)* | Group II intron reverse transcriptase/  maturase (*Burkholderia*) | Gfo/Idh/MocA family oxidoreductase (*Pedobacter*) | ATP-grasp domain protein (*Pseudomonas*) | Inosamine-phosphate amidinotransferase 1 (*Streptomyces*) | Branched-chain amino acid aminotransferase (Mixed origin) | 2OG-Fe(II) oxygenase (*Pseudomonas*) | Argininosuccinate synthase (*Burkholderia*/  *Salmonella*) | Contig end |
| --- | --- | --- | --- | --- | --- | --- | --- | --- | --- | --- |
| *Ca.* Burkholderia ardisicola Acor | IS630 | **EEVS** (Ψ)* (CBARDCOR_4200) *Pseudomonas*) | ROK family (*Pseudomonas*) | DegT/DnrJ/EryC1/StrS family aminotrasferase (*Pseudomonas*) | 6-phospho-beta-glucosidase (*Pseudomonas*) | Gfo/Idh/MocA family oxidoreductase (*Pseudomonas*) | 3-phosphoshikimate 1-carboxylvinyltransferase (*Pseudomonas*) | HAD family hydrolase (*Pseudomonas*) | Class-I Dependent methyltransferase (Ψ; *Pseudomonas*) | Contig end |
| *Ca.* Paraburkholderia dryadicola | IS3 | Hypothetical protein (*Burkholderia*) | **EEVS** (CPDRYDRY_6570) (*Burkholderia singularis/*  *Streptomyces*) | SDR family oxidoreductase (*Burkholderia singularis/*  *Streptomyces*) | GMC family oxidoreductase (*Burkholderia singularis/*  *Streptomyces*) | DegT/DnrJ/EryC1/StrS family aminotransferase (*Burkholderia* *singularis*/*Streptomyces*) | GNAT-family N-acetyltransferase (*Burkholderia*/  *Pseudomonas*) | Carbamoyl transferase (*Pseudomonas*) | Contig end |  |
