## Supplementary information for "Cyclitol secondary metabolism is a central feature of *Burkholderia* leaf symbionts"

**Plant collection and extraction**

Fresh leaves of *Fadogia homblei*, *Vangueria macrocalyx, V. pygmaea, V. dryadum, V. infausta, V. lasiantha, V. madagascariensis, V. randii and V. soutpansbergensis* were collected (Table SI-1) and stored at 5 °C. Leaves of *Psychotria kirkii* were collected from the Botanical Garden of Zurich, Switzerland and used as positive control for kirkamide and streptol-glucoside (Hsiao et al., 2019; Pinto-Carbó et al., 2016).

The extraction method of Sieber et al. (2015) was adjusted as follows: Dried leaf material (3 g of 5x5 mm pieces) was extracted with 80% distilled methanol (Merck Ltd) in a speed-extractor (Büchi E-91) at 50 °C and 100 bar. The speed-extractor was set to 5 cycles with a heating phase of 1 min each, a solvent holding phase of 9 min and a discharge phase of 5 min. The extracts were dried using a Büchi Genevac plus centrifugal evaporator (EZ-2 Plus) at 45 °C and set to low boiling point. After extraction and drying, all the extracts were stored at approximately 5 °C.

**Table SI-1**: Plant species analysed for the production of kirkamide, streptol and streptol glucoside. *Psychotria kirkii* was collected from the Botanical Garden of Zurich, Switzerland and used as positive control. PRU: H.G.W.J. Schweickerdt Herbarium, University of Pretoria, South Africa.

| **Plant species** | **Coordinates** | **Location of collection** | **Pant tissue collected** | **Voucher number** |
| --- | --- | --- | --- | --- |
| ***Fadogia homblei*** | S 25° 34' 20,5''  E 28° 25' 58,4'' | Roodeplaat | Leaf tissue | PRU 128010 |
| ***Vangueria dryadum*** | S 25° 26' 35,5''  E 30° 58' 9,3'' | Lowveld National Botanical Gardens | Leaf tissue | PRU 128005 |
| ***Vangueria infausta*** | S 25° 26' 35,2''  E 30° 58' 16,4'' | Lowveld National Botanical Gardens | Leaf tissue | PRU 128003 |
| ***Vangueria lasiantha*** | S 25° 26' 36,5''  E 30° 58' 9,4'' | Lowveld National Botanical Gardens | Leaf tissue | PRU 128007 |
| ***Vangueria macrocalyx*** | S 25° 26' 34,6''  E 30° 58' 15,6'' | Lowveld National Botanical Gardens | Leaf tissue | PRU 128006 |
| ***Vangueria madagascariensis*** | S 25° 26' 34,7''  E 30° 58' 16'' | Lowveld National Botanical Gardens | Leaf tissue | PRU 128004 |
| ***Vangueria pygmaea*** | S 25° 44' 10,2''  E 28° 31' 59,3'' | Cullinan | Leaf tissue | PRU 126008 |
| ***Vangueria randii*** | S 25° 26' 41,4''  E 30° 58' 7,7'' | Lowveld National Botanical Gardens | Leaf tissue | PRU 128008 |
| ***Vangueria soutpansbergensis*** | S 25° 26' 34,5''  E 30° 58' 15,6'' | Lowveld National Botanical Gardens | Leaf tissue | PRU 128002 |

**Description of novel bacterial taxa**

**Description of *Candidatus* Burkholderia ardisicola sp. nov.**

Burkholderia ardisicola [ar.di.si.i’co.la N.L. fem. n. Ardisia a plant genus; L. suff. -cola (from L. n. incola) a dweller, inhabitant; N.L. fem. n. ardisicola a dweller of Ardisia]. Not cultivated. Rod-shaped. Obtains energy by respiration. Detected as a symbiont in leaf galls of *Ardisia cornudentata* and *Ardisia mamillata* from the Ghent University Botanical Garden (Belgium) in 2018. Represented by the draft genome (GenBank GCA_940588625).

**Description of *Candidatus* Burkholderia dryadicola sp. nov.**

Burkholderia dryadicola [dry.a.di’co.la dryadum specific epithet of a host plant species; L. suff. -cola (from L. n. incola) a dweller, inhabitant; N.L. fem. n. dryadicola a dweller of (*Vangueria*) *dryadum*]. Not cultivated. Rod-shaped. Obtains energy by respiration. Detected as a symbiont in leaves of *Vangueria dryadum* and *Vangueria macrocalyx* from the Lowveld National Botanic Gardens (South Africa) in 2019. Represented by the draft genome (GenBank GCA_940590165).

**Description of *Candidatus* Paraburkholderia soutpansbergensis sp. nov.**

Burkholderia soutpansbergensis [sout.pans.ber.gen.si’s N.L. fem. adj. soutpansbergensis, name based on the specific epithet of the host plant]. Not cultivated. Rod-shaped. Obtains energy by respiration. Detected as a symbiont in leaves of *Vangueria soutpansbergensis* from the Lowveld National Botanic Gardens (South Africa) in 2019. Represented by the draft genome (GenBank GCA_940746715).

**Analytical Chemistry - Methods**

**GC-MS analysis of kirkamide:**

Extracts were derivatised with N-methyl-N-(trimethyl-silyl)-trifluoroacetamide (MSTFA, Merck Ltd) according to the method of Pinto-Carbó *et al.* (2016). Three replicates of the plant extracts were derivatised from a concentration of 1 mg/ml in 2 ml double distilled water as follows: The extracts were filtered through 0.22 µm syringe fitted filters and 100 µl transferred to 2.0 ml screw top glass vials with 200 µl inserts and dried overnight under a nitrogen stream. The residues were dissolved in 50 µl MSTFA, vortexed for two minutes, left at 70 °C for one hour and then 50 µl pyridine was added as the solute.

The derivatised samples were analysed on a Shimadzu GC-MS-QP2010 (Shimadzu Corporation, Japan) with ionization energy set at 70 eV. The compounds were separated using a Rtx – 5MS column (29.3 m x 250 µm x 0.25 µm i.d.; 0.25 µm df) with helium as the carrier gas. Splitless injections of 1 µl were performed, with the column flow set to linear velocity. Sampling time was set to 2 min, with the solvent cut-off time set to 3.5 min. The injector and interface temperatures were set at 250°C. The GC oven temperature program was set to an initial 40°C and held for 1 min, thereafter it was increased to 330°C at a rate of 7°C min^-1^ which was held for 10 min, bringing the total run time to 52 min. The MS ion source and interface temperatures were set to 250°C. The detector voltage was set to 0.1 kV, relative to the instrument tuning results. The mass-to-charge ratio (m/z) detection was set to start at 7 min (ensuring complete solvent elimination) and ranged from 45 to 650 m/z with a scan speed of 2 500 aum s^-1^. Pyridine was used as a blank at the start of the analysis to observe any instrumental errors.

**UPLC-QToF-MS analysis of streptol and streptol glucoside**

The presence of underivatised streptol and streptol glucoside in the plant extracts was analysed using a Waters Synapt G2 high-definition mass spectrometry (HDMS) system (Waters Inc., Milford, Massachusetts, USA). The apparatus consists of a Waters Acquity UPLC connected to a quadropole-time-of-flight (QToF) instrument. The method of Georgiou et al. (2021) was followed for the detection of streptol and streptol glcoside in negative mode [M-H]^-^. The samples were analysed using a Luna Omega 1.6 µm C_18_ 100 A, 100 x 2.1 mm (Phenomenex, Separations) column and a solvent system that consisted of MeCN:H_2_O (A, 8:2, 0.1 % NH_4_OAc) and MeCN:H_2_O (B, 2:8, 0.1 % NH_4_OAc). The gradient was set to start at 95 % of B and to decrease to 50 % of B in 7 min, for the next 2 min the gradient was kept at 50 % of B, the gradient was then gradually decreased from 50 % to 5 % of B for the next 3 min and was followed by a column wash for the next 2 min giving a total run time of 12 min. The column temperature was 40 °C, injection volume 7 µl and the flow rate 0.3 ml min^-1^.

Mass to charge ratios (*m/z*) were recorded between 50 and 1 200 Da. High energy collision induced dissociation (CID) was used for tandem MS fragmentation. The collision energy for the ramping was set to increase from 10 V to 20 V in order to get a range of data.

**Analytical Chemistry - Results**

Derivatised samples of the Rubiaceae species, *Psychotria kirkii* (positive control), *Fadogia homblei*, *Vangueria dryadum*, *V. infausta*, *V. lasiantha*, *V. macrocalyx*, *V. madagascariensis*, *V. pygmaea* *V. randii*, and *V. soutpansbergensis* were analysed for the presence of kirkamide by GC-MS. The ion fragments 73, 147, 282, 332, 415, 431, 490 and 505 *m/z* were used to confirm the presence of kirkamide in the crude derivatised extracts). All the fragments were present in the extract of *P. kirkii* at rt 27.830, confirming presence of kirkamide in this species, but these were not detected in any of the other plant species (Figures SI-1 to SI-10).

An initial UPLC-QToF-MS analysis showed the presence of streptol and streptol gluscoside in *P. kirkii* and in some of the gousiekte causing species (Figure SI-11), but this needs further confirmation with plant material that will be collected during the spring season (Sept-Oct). The streptol (m/z = 175.0607, [M-H]^-^, calculated MF = C_7_H_12_O_5_ at 2.8 ppm) ion fragments 85, 111, 121 and 175 *m/z* and the streptol glucoside (m/z = 337.1126, [M-H]-, calculated MF = C_13_H_22_O_10_ at 4.2 ppm) ones of 112, 139, 175 and 337 *m/z* were used in the analyses. Streptol eluted at 0.716 min and streptol glucoside at 0.774 min.

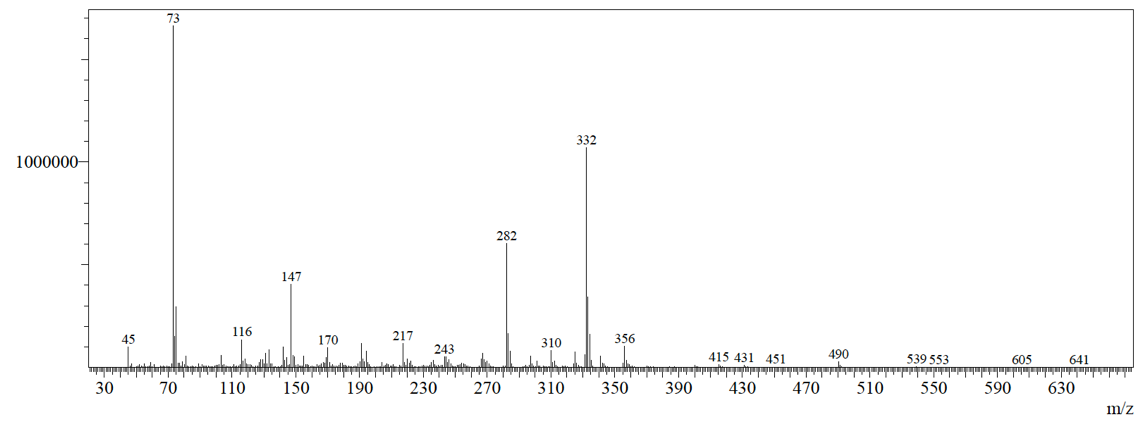

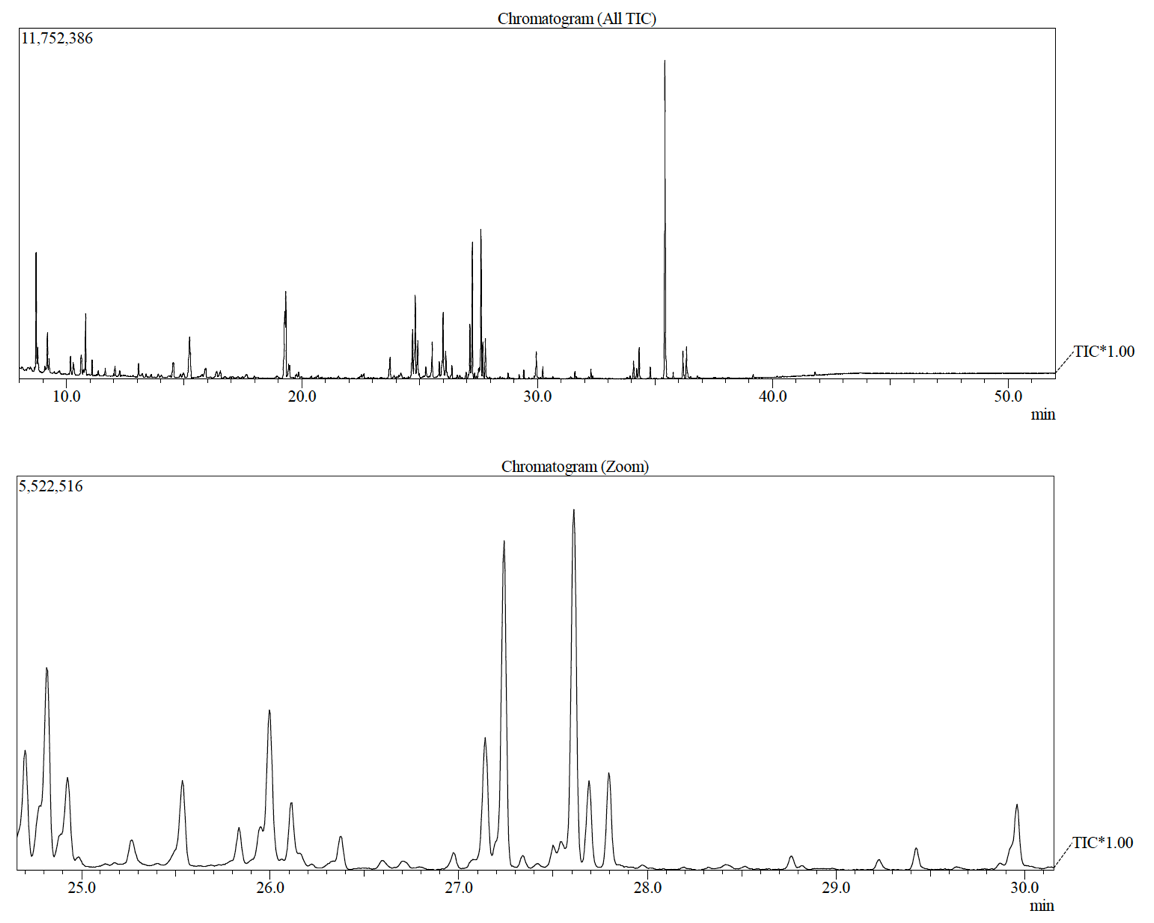

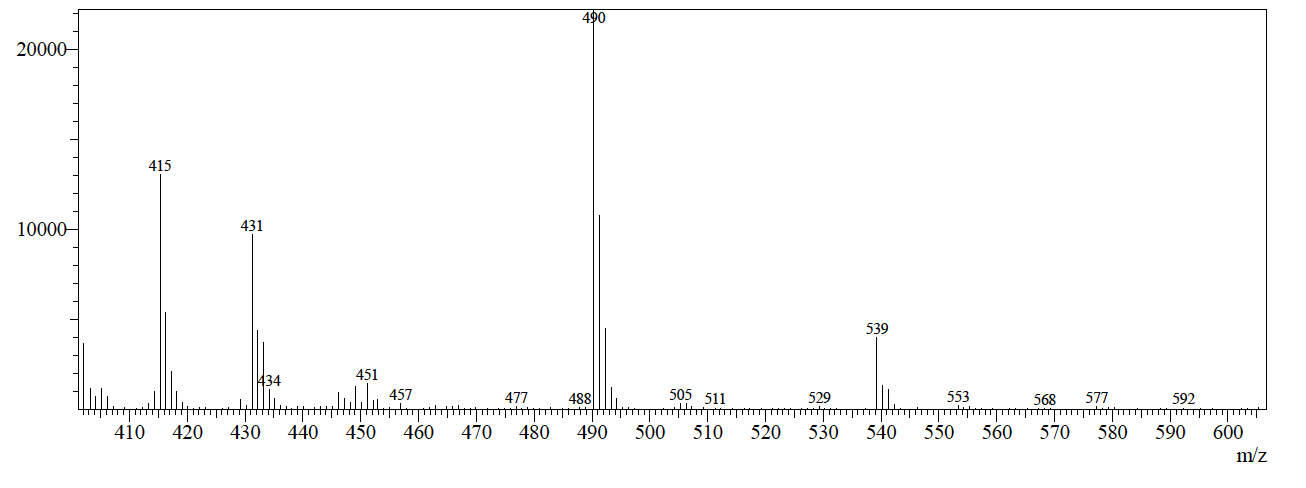

**Figure SI-1**: GC-MS results of *Psychotria kirkii* showing the presence of kirkamide in the chromatograms and the mass spectra with ion fragments 73, 147, 282, 332, 415, 431, 490 and 505 *m/z* and the enlarged *m/z* region of kirkamide from 410 to 600 *m/z*.

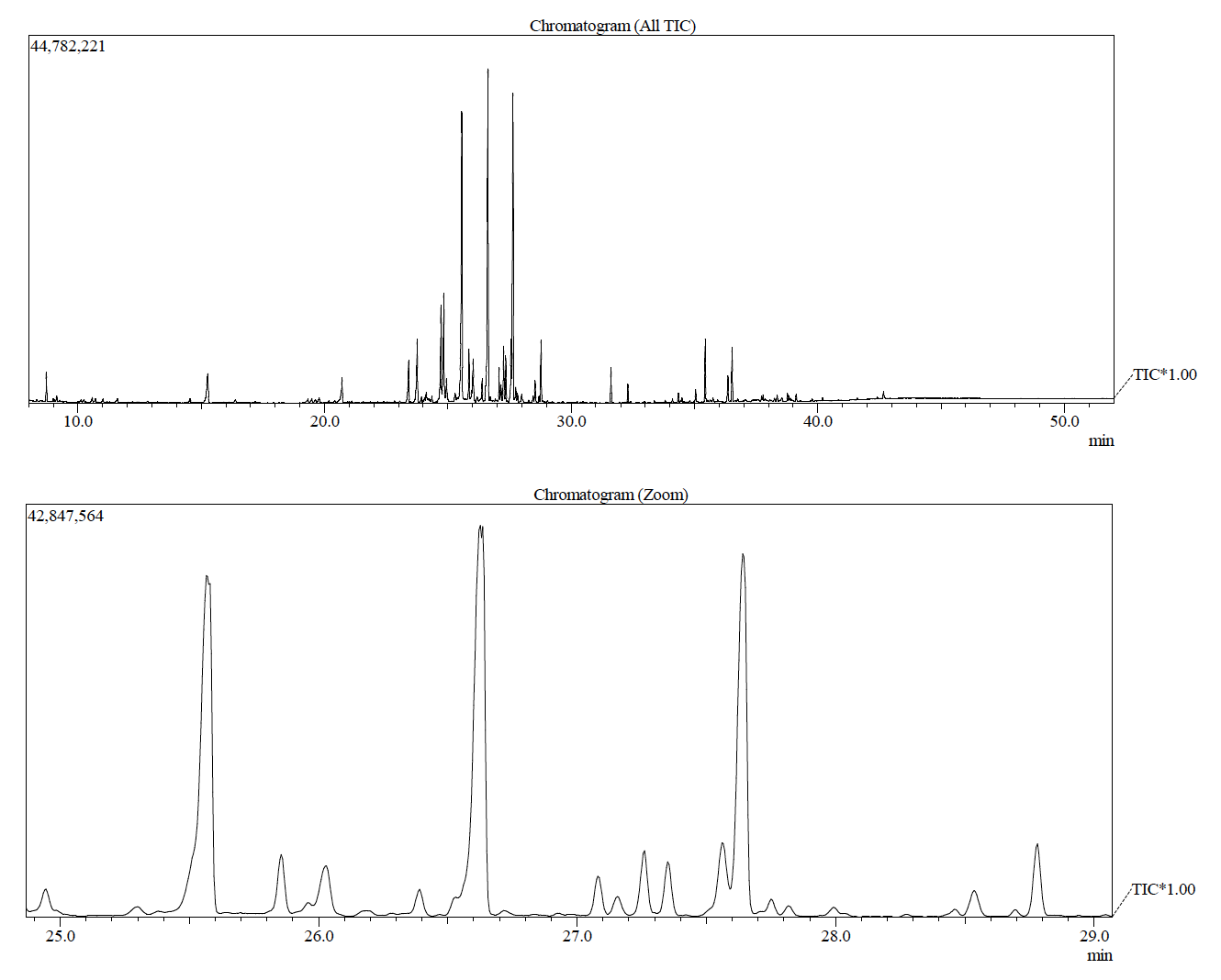

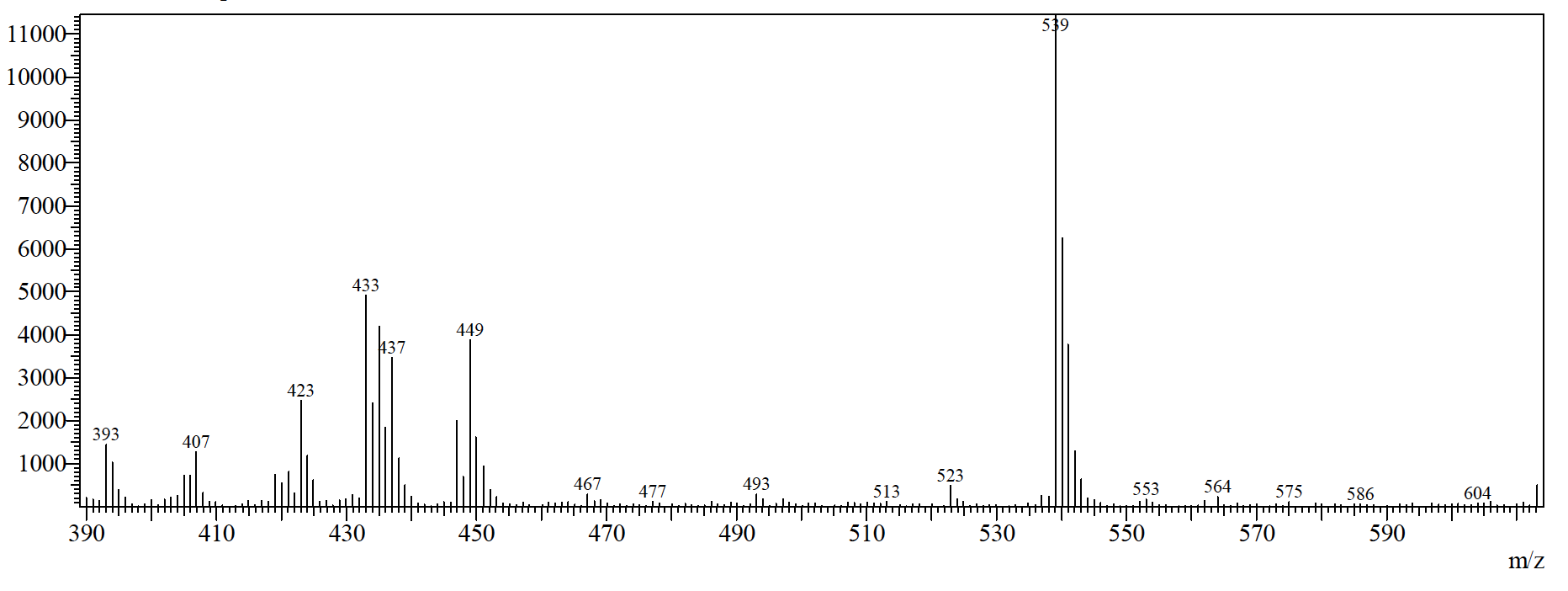

**Figure SI-2**: GC-MS result of *Fadogia homblei* showing the absence of kirkamide in the chromatograms and the enlarged *m/z* region from 400 to 600 *m/z*.

**Figure SI-3**: GC-MS result of *Vangueria dryadum* showing the absence of kirkamide in the chromatograms and the enlarged *m/z* region from 400 to 600 *m/z*.

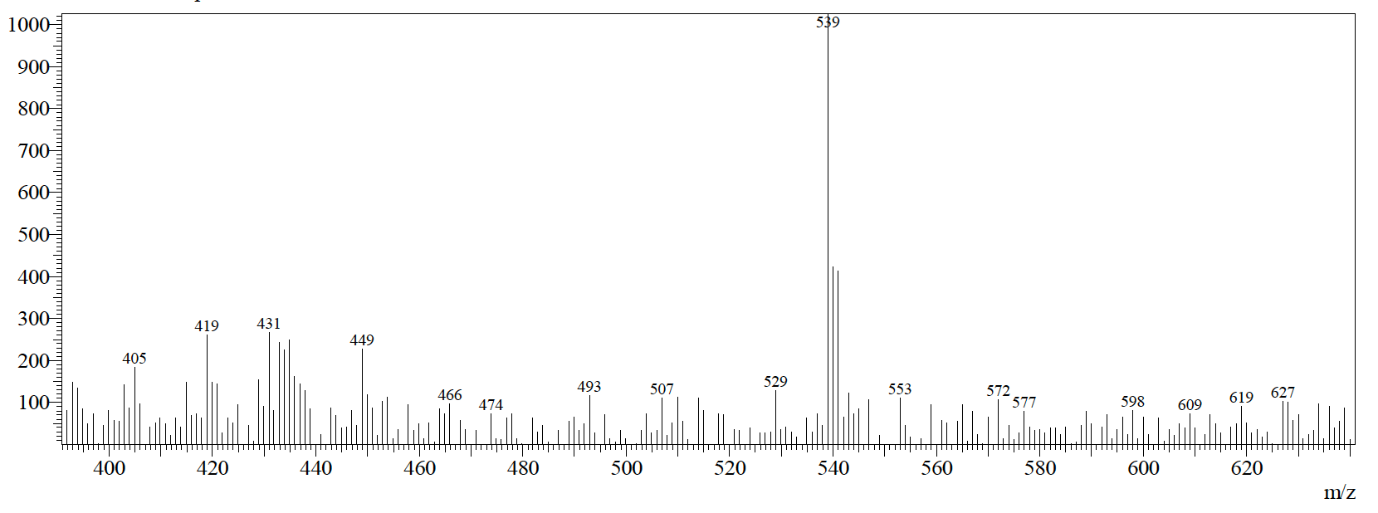

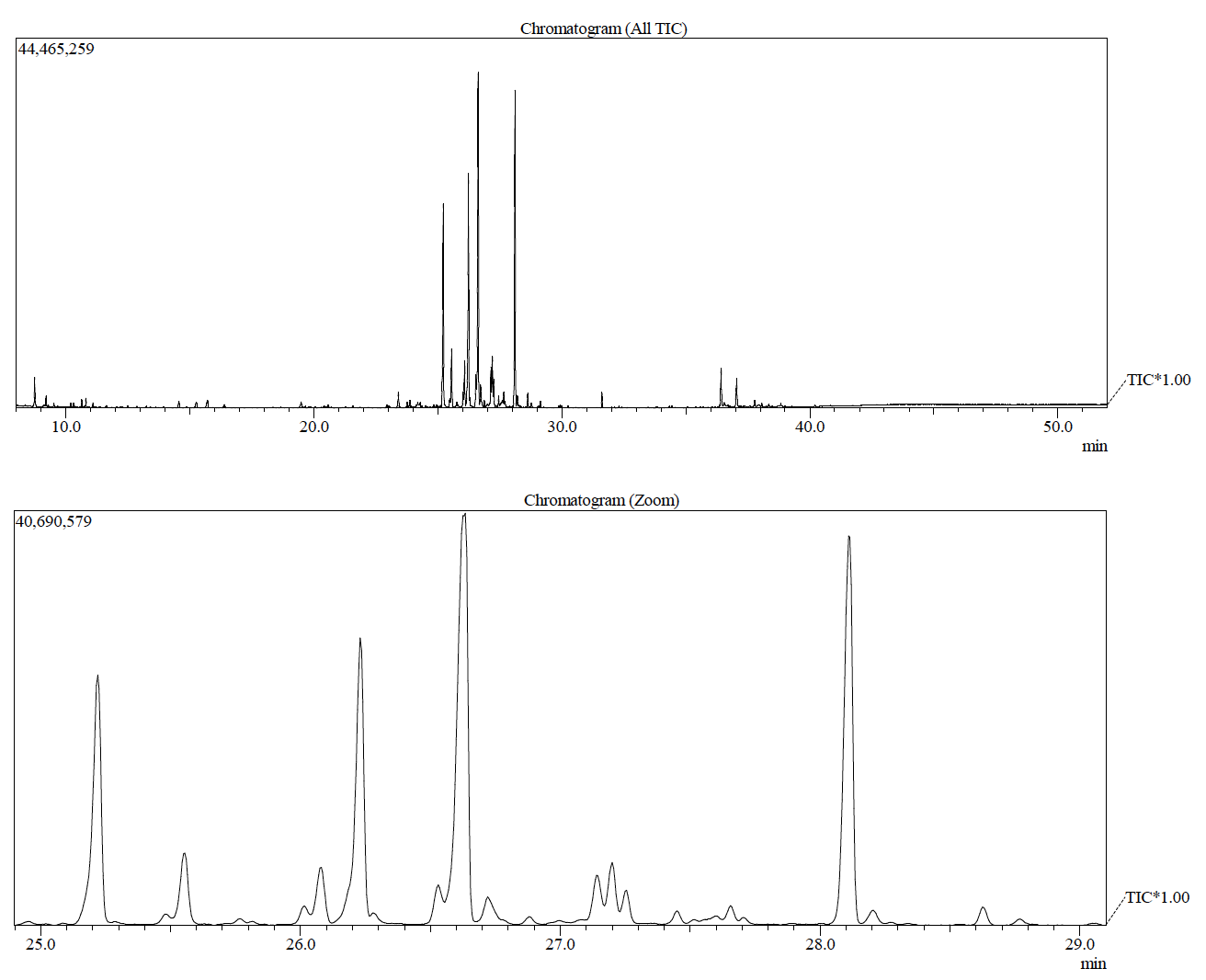

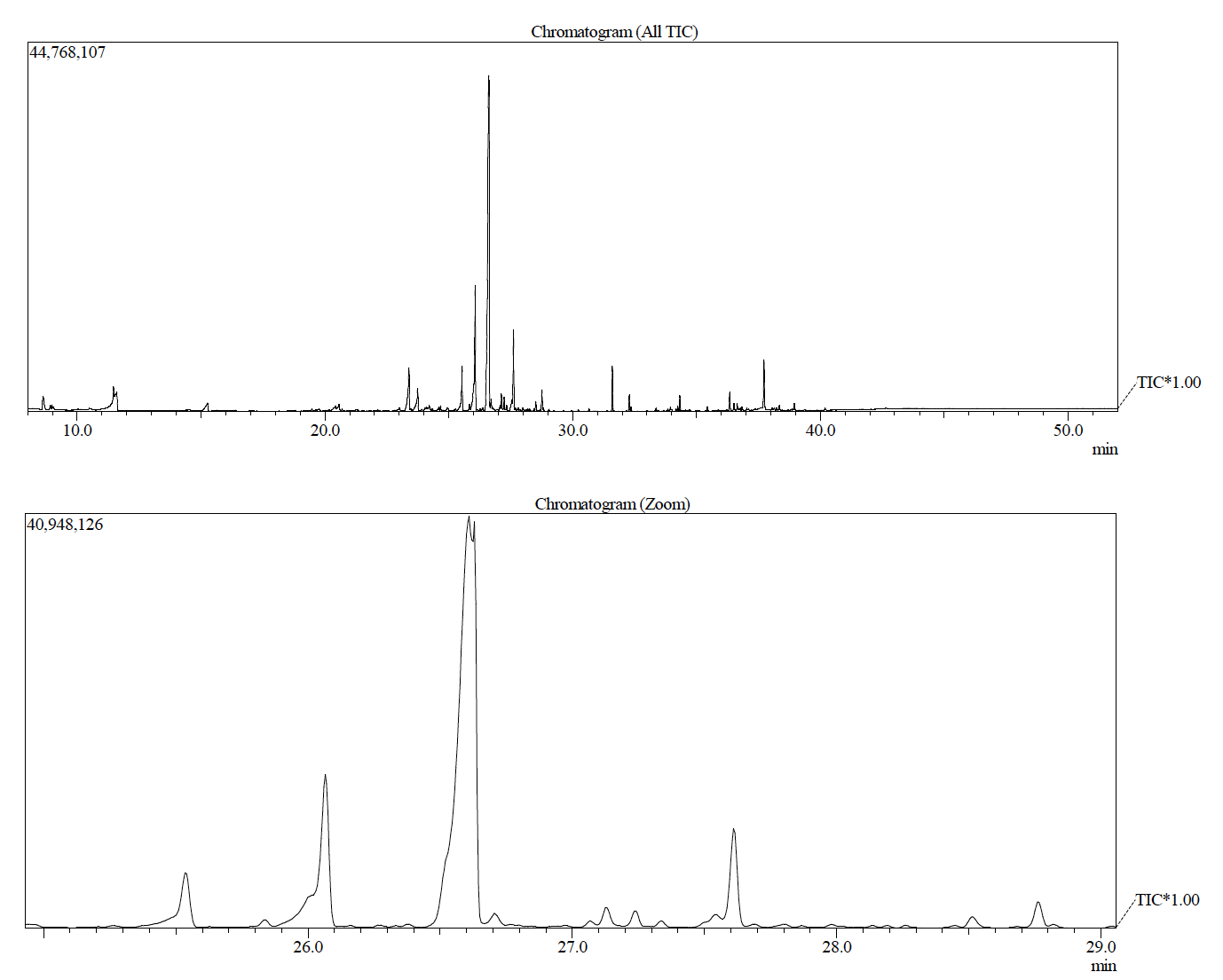

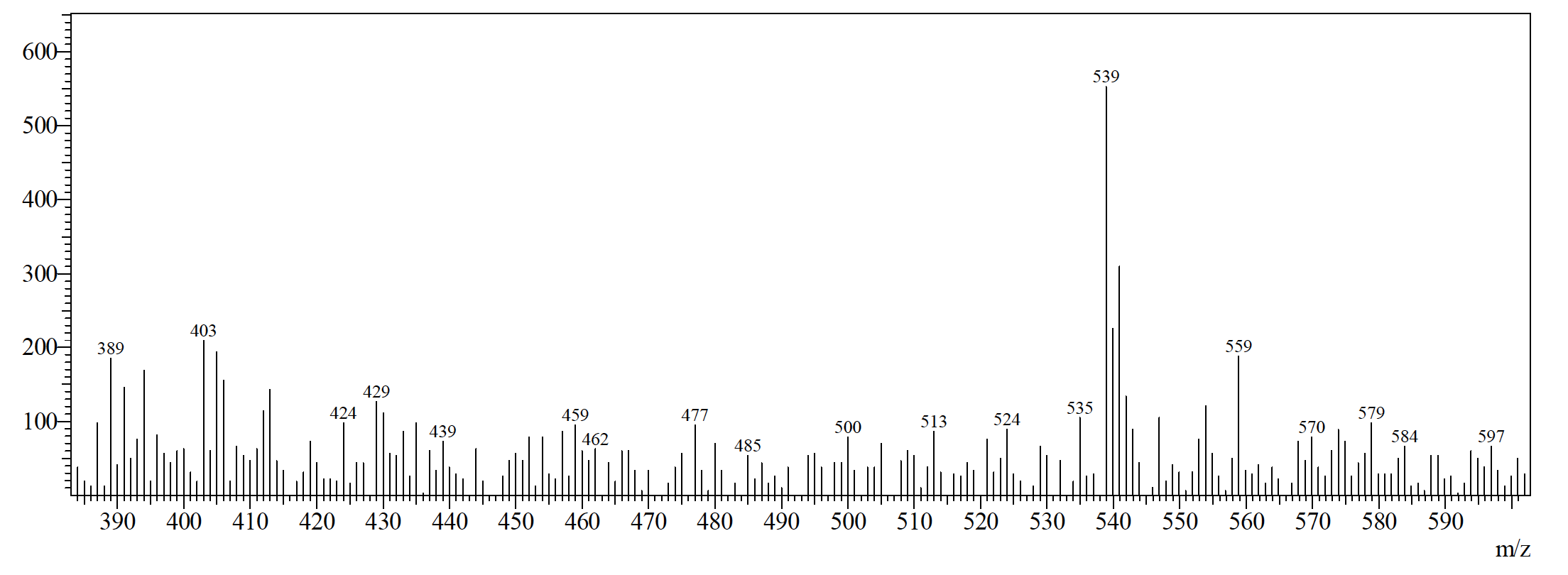

**Figure SI-4**: GC-MS result of *Vangueria infausta* showing the absence of kirkamide in the chromatograms and the enlarged *m/z* region from 400 to 600 *m/z*.

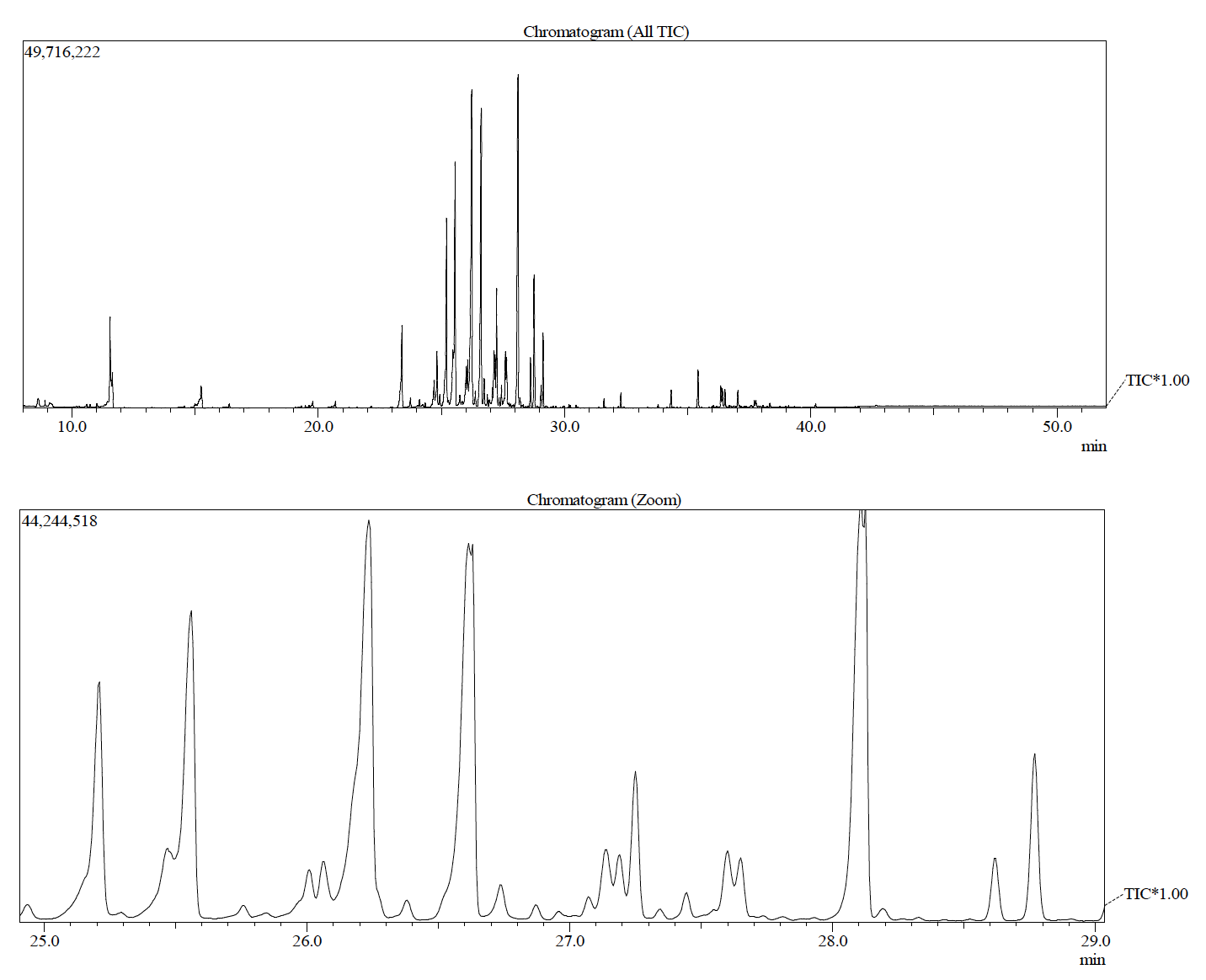

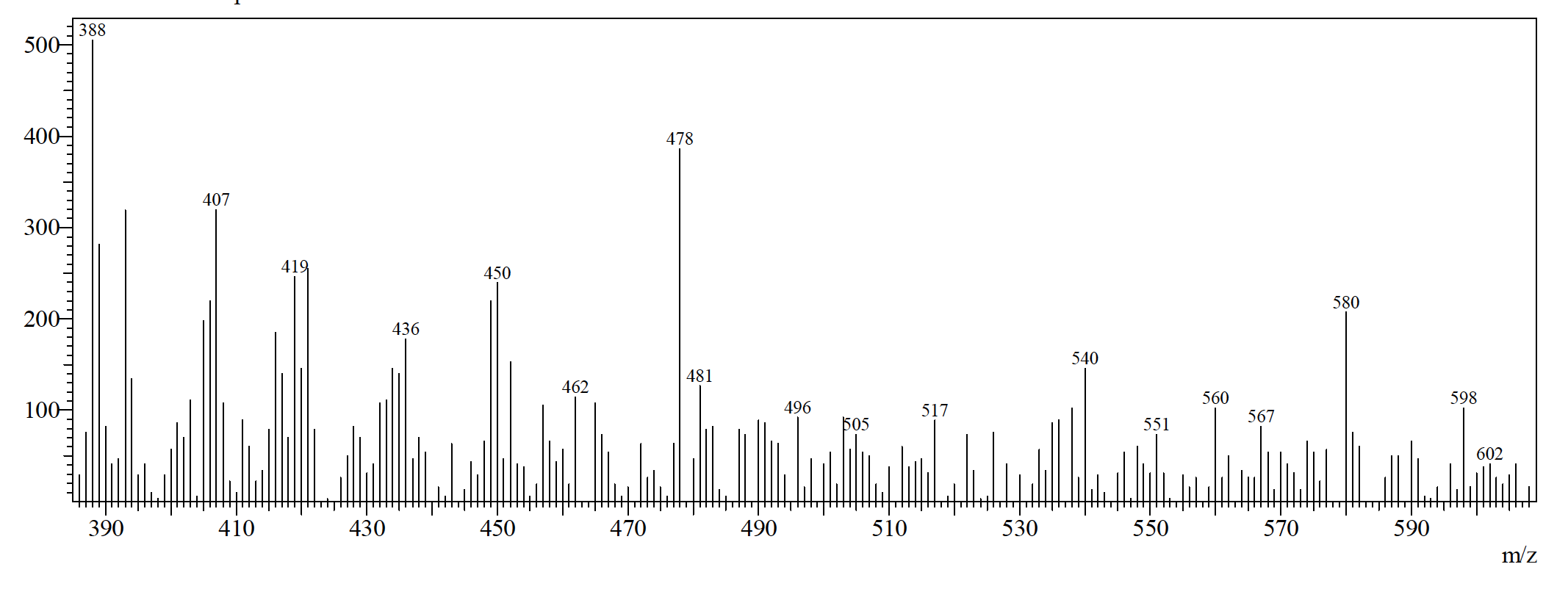

**Figure SI-5**: GC-MS result of *Vangueria lasianta* showing the absence of kirkamide in the chromatograms and the enlarged *m/z* region from 400 to 600 *m/z*.

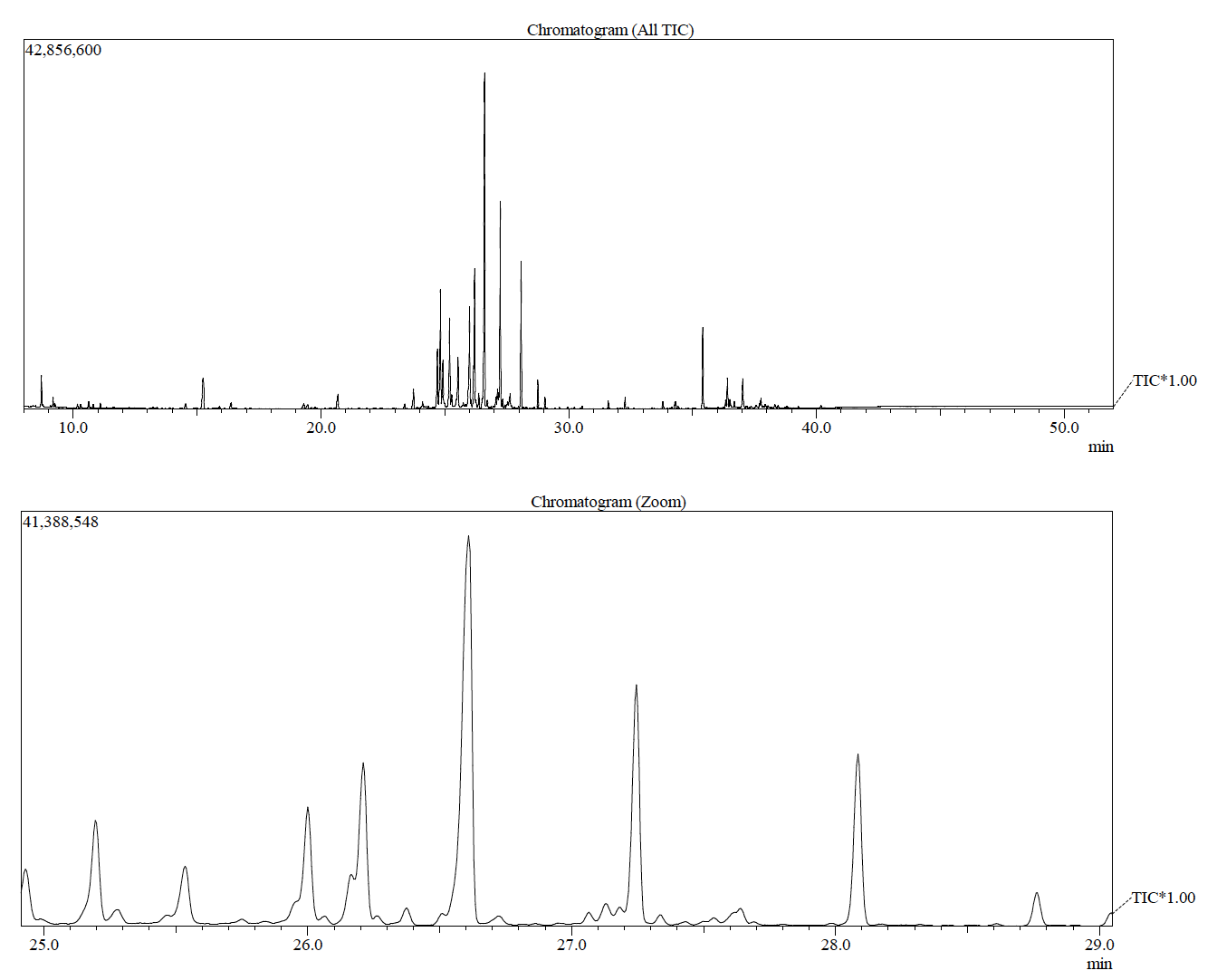

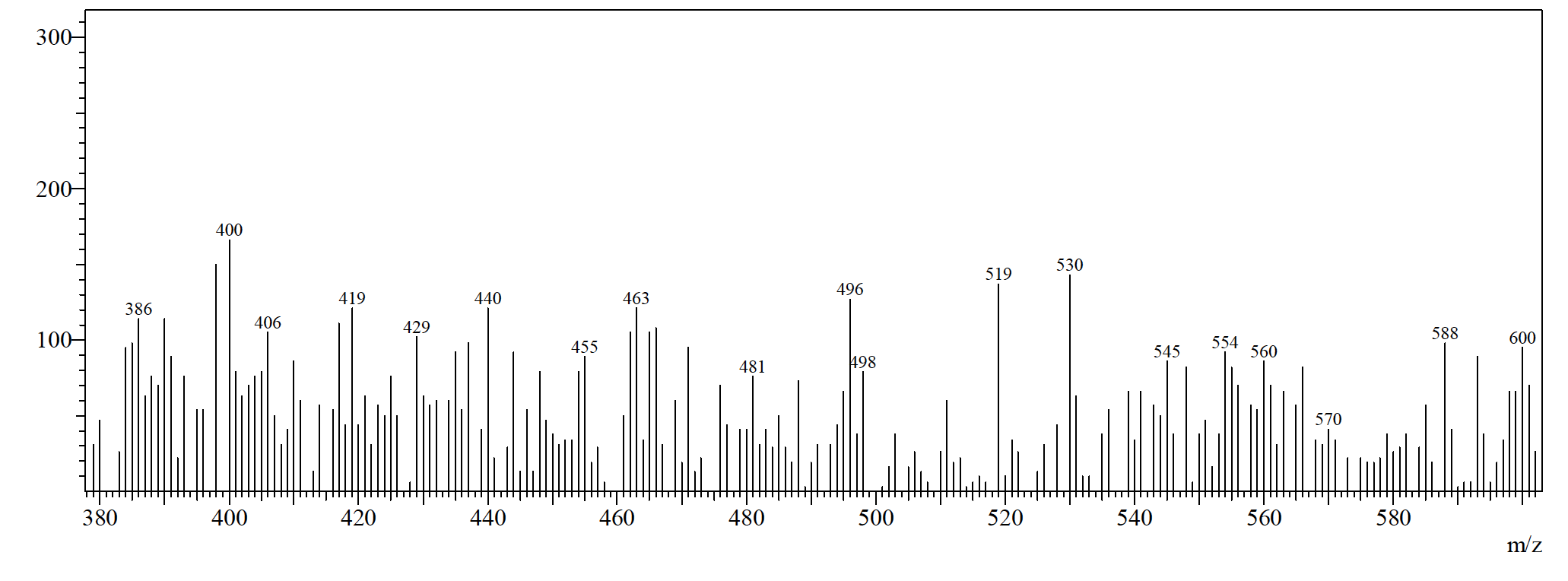

**Figure SI-6**: GC-MS result of *Vangueria macrocalyx* showing the absence of kirkamide in the chromatograms and the enlarged *m/z* region from 400 to 600 *m/z*.

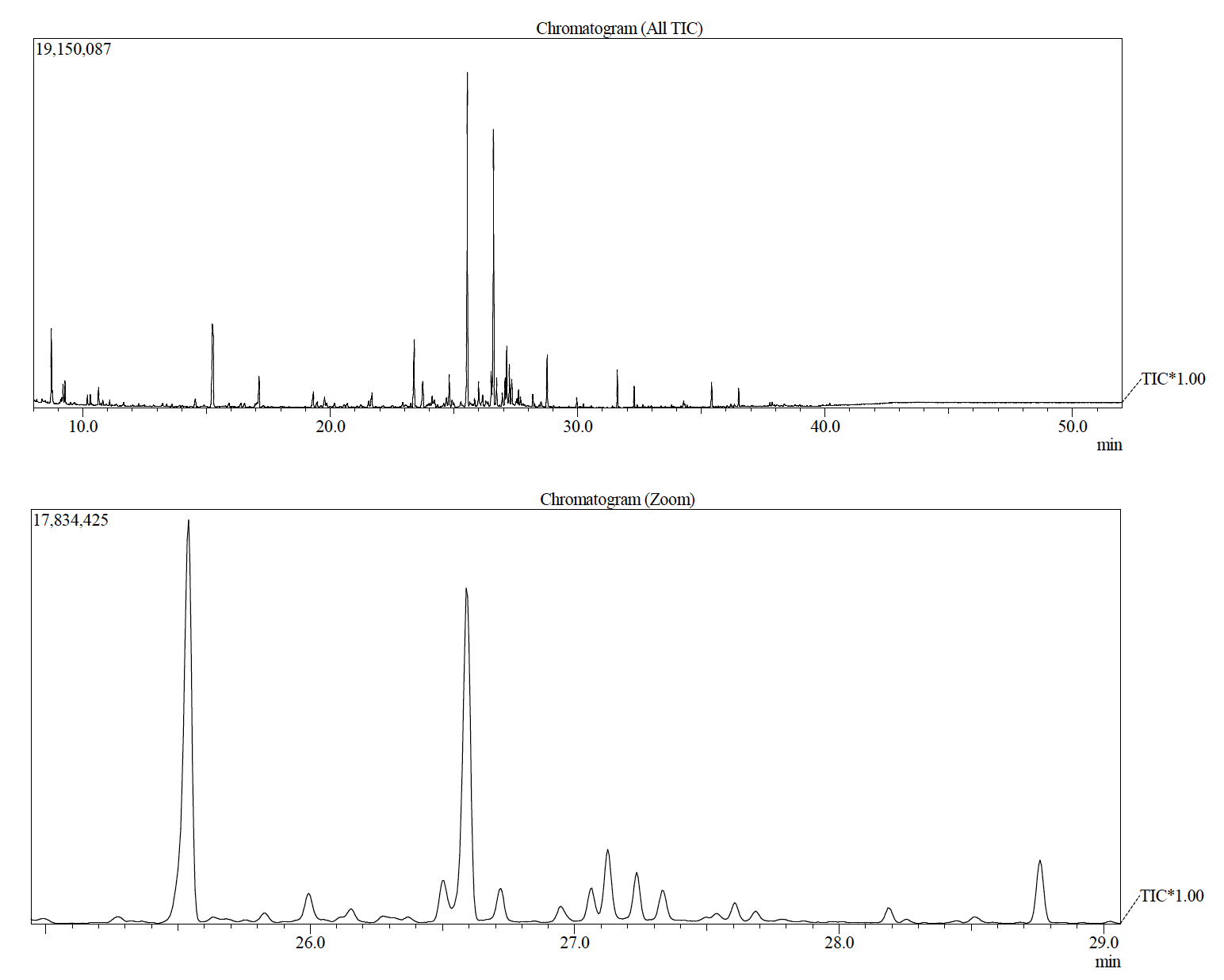

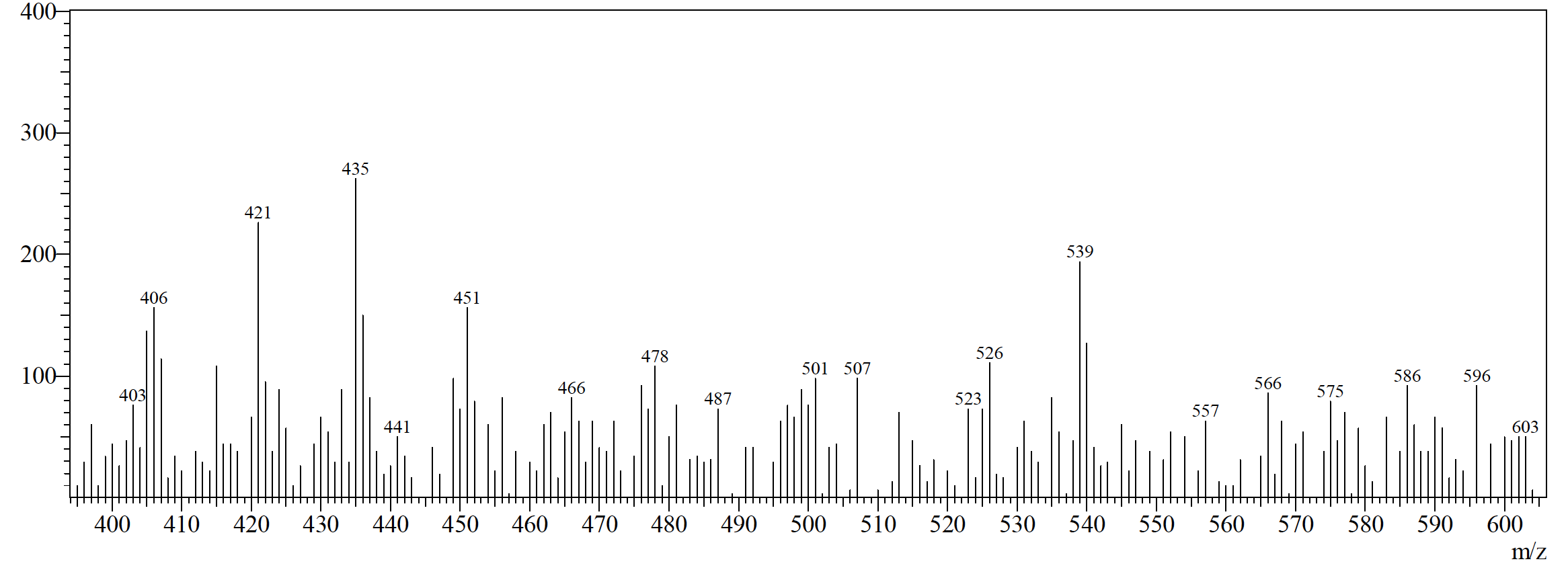

**Figure SI-7**: GC-MS result of *Vangueria madagascariensis* showing the absence of kirkamide in the chromatograms and the enlarged *m/z* region from 400 to 600 *m/z*.

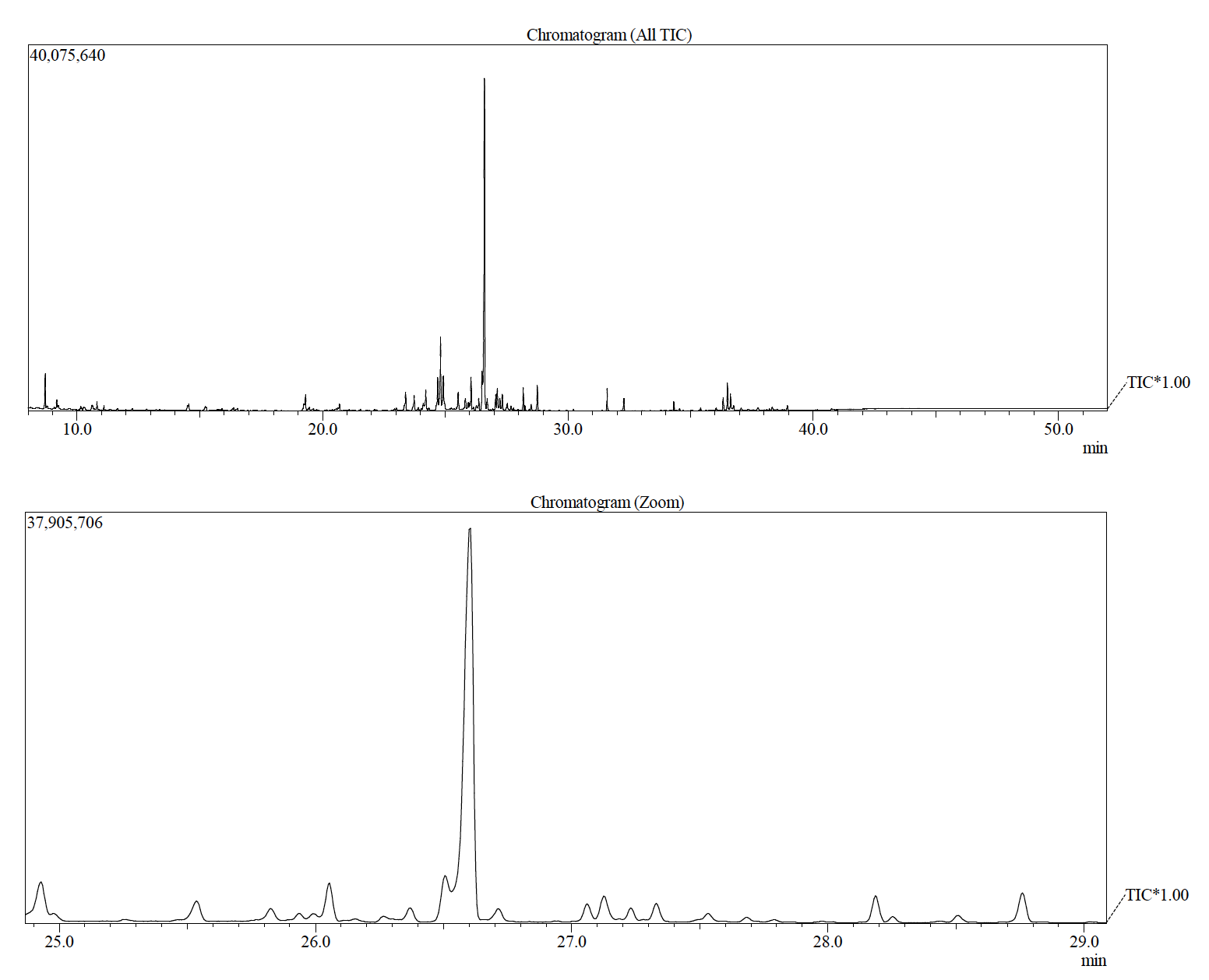

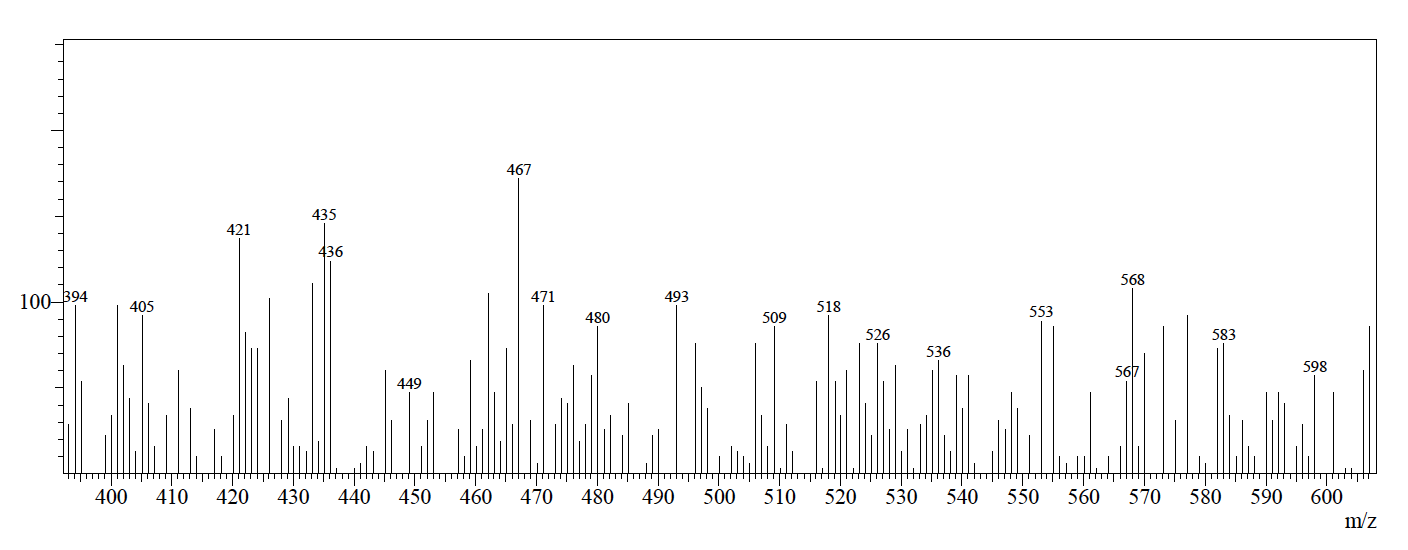

**Figure SI-8**: GC-MS result of *Vangueria pygmaea* showing the absence of kirkamide in the chromatograms and the enlarged *m/z* region from 400 to 600 *m/z*.

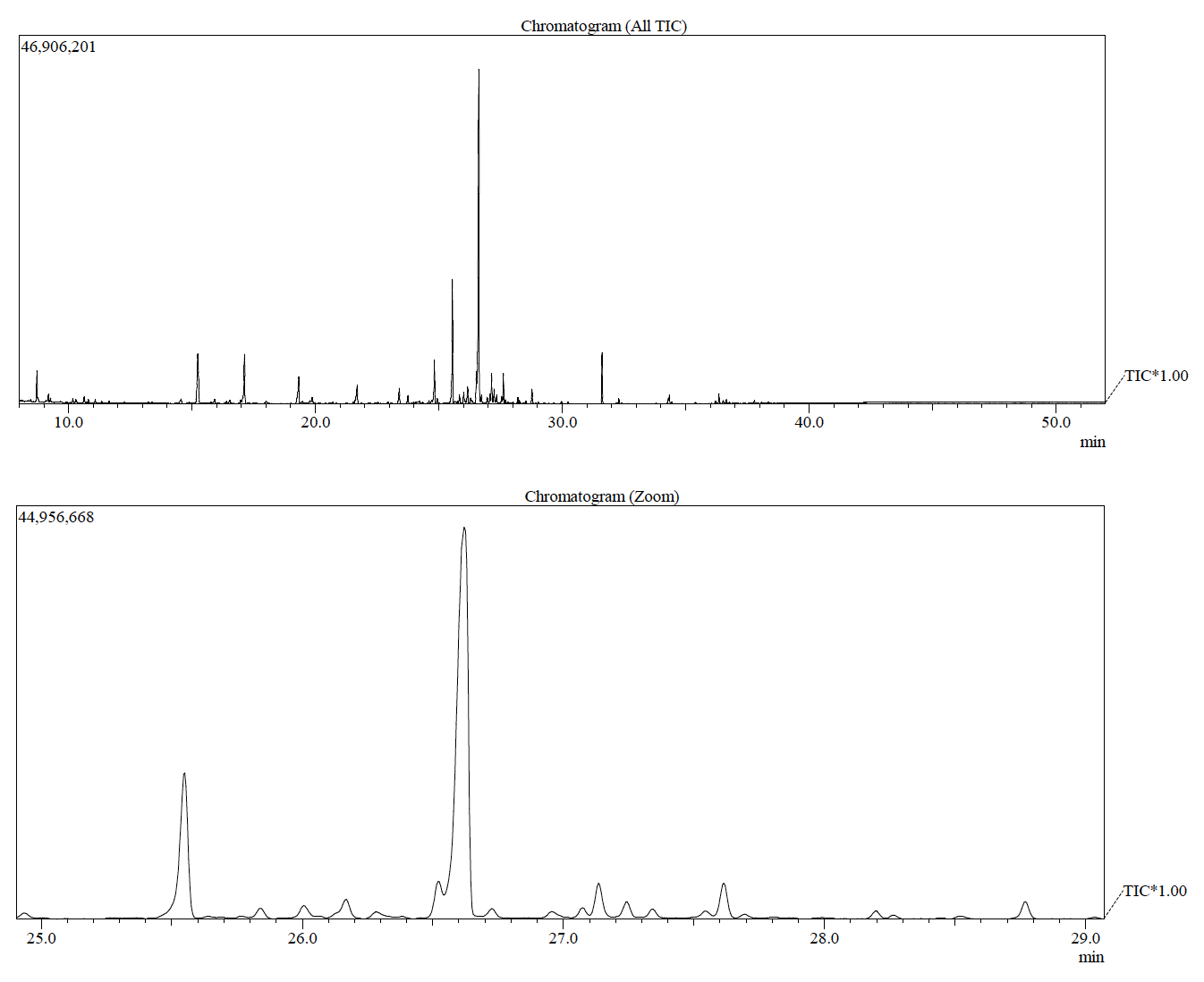

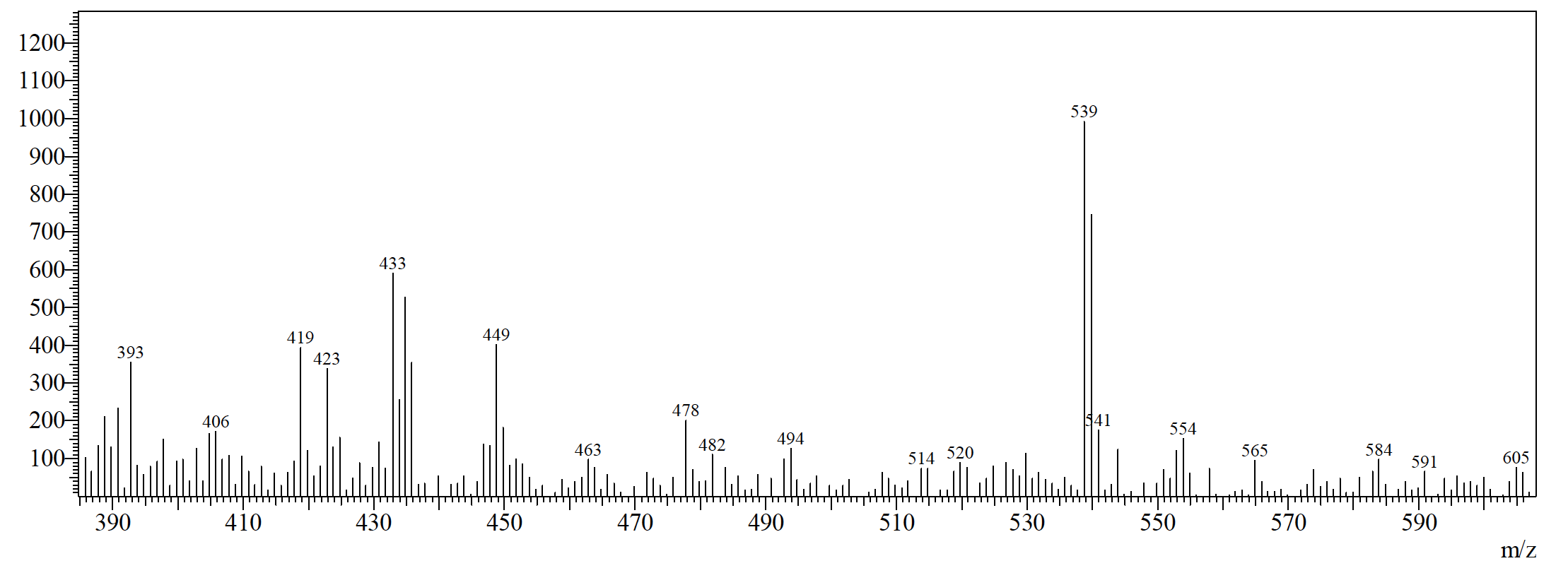

**Figure SI-9**: GC-MS result of *Vangueria randii* showing the absence of kirkamide in the chromatograms and the enlarged *m/z* region from 400 to 600 *m/z*.

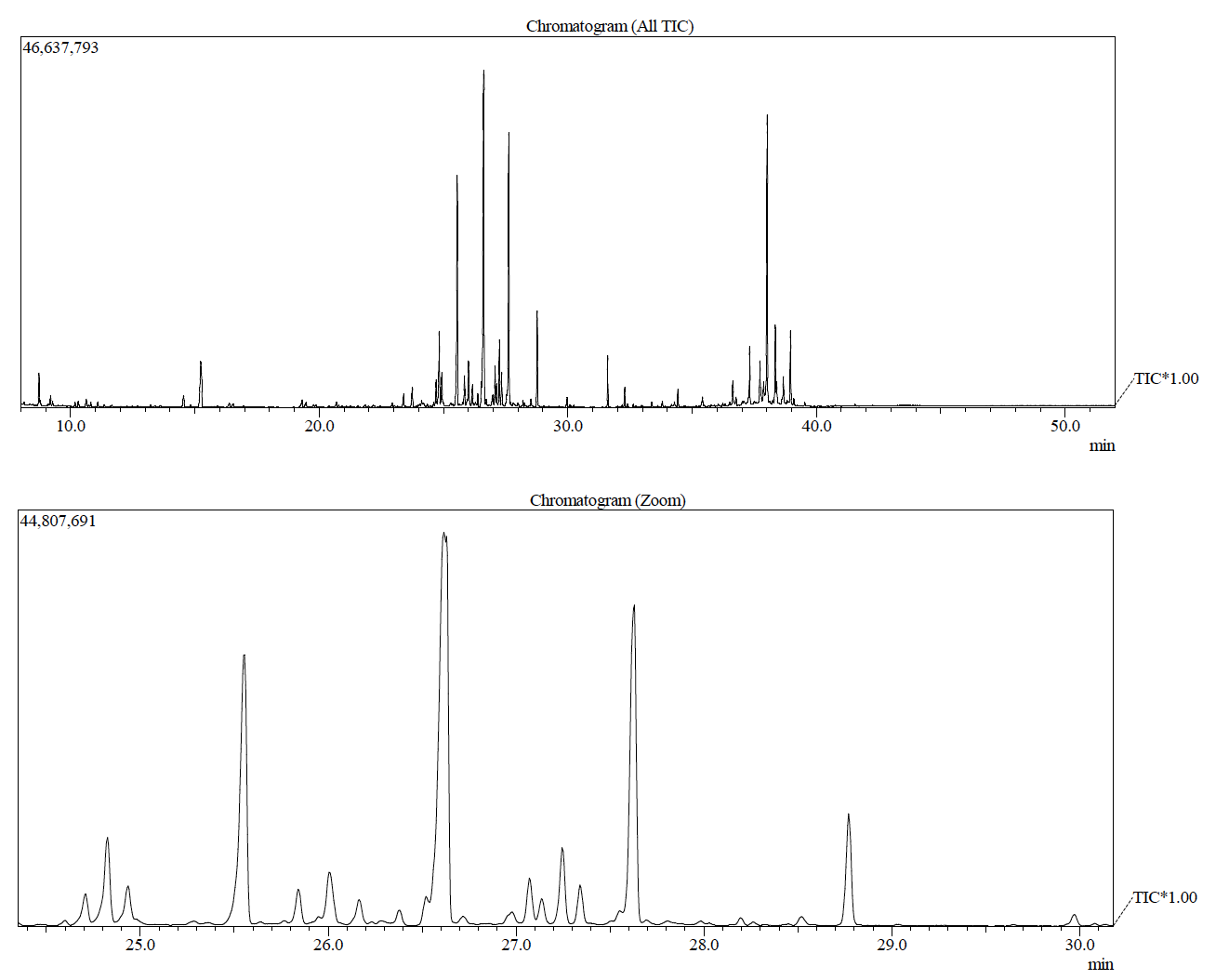

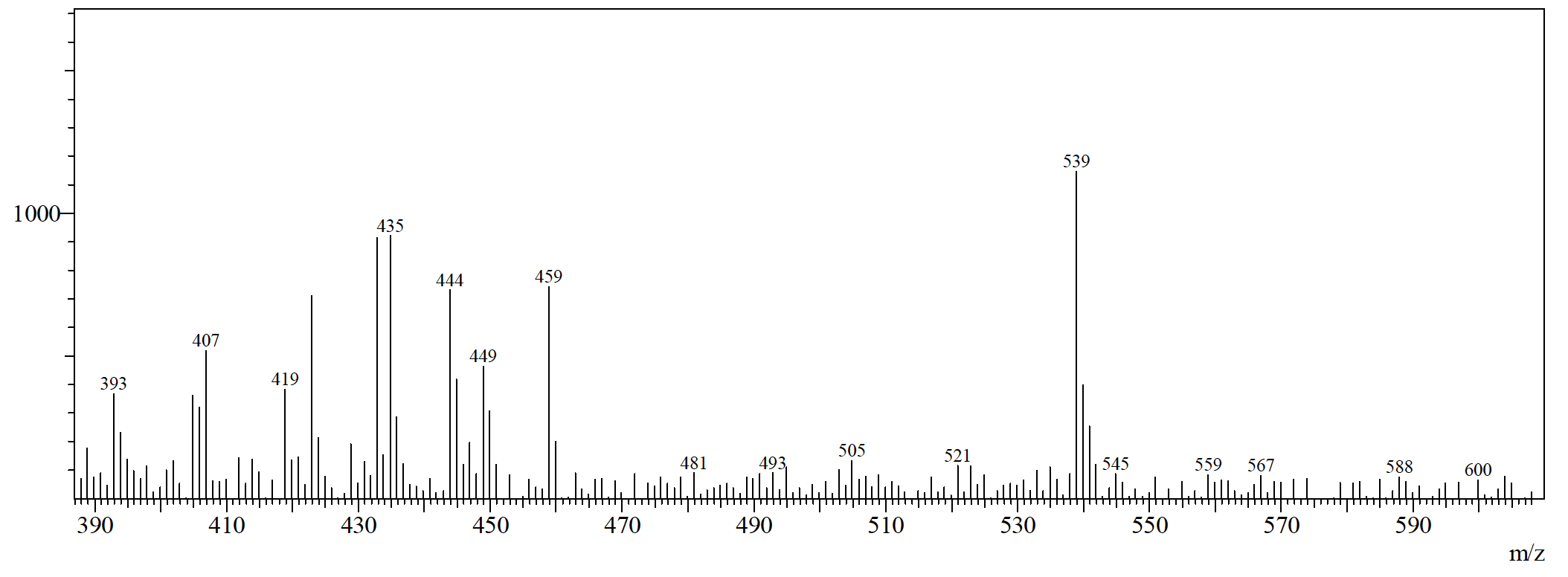

**Figure SI-10**: GC-MS result of *Vangueria soutpansbergensis* showing the absence of kirkamide in the chromatograms and the enlarged *m/z* region from 400 to 600 *m/z*.

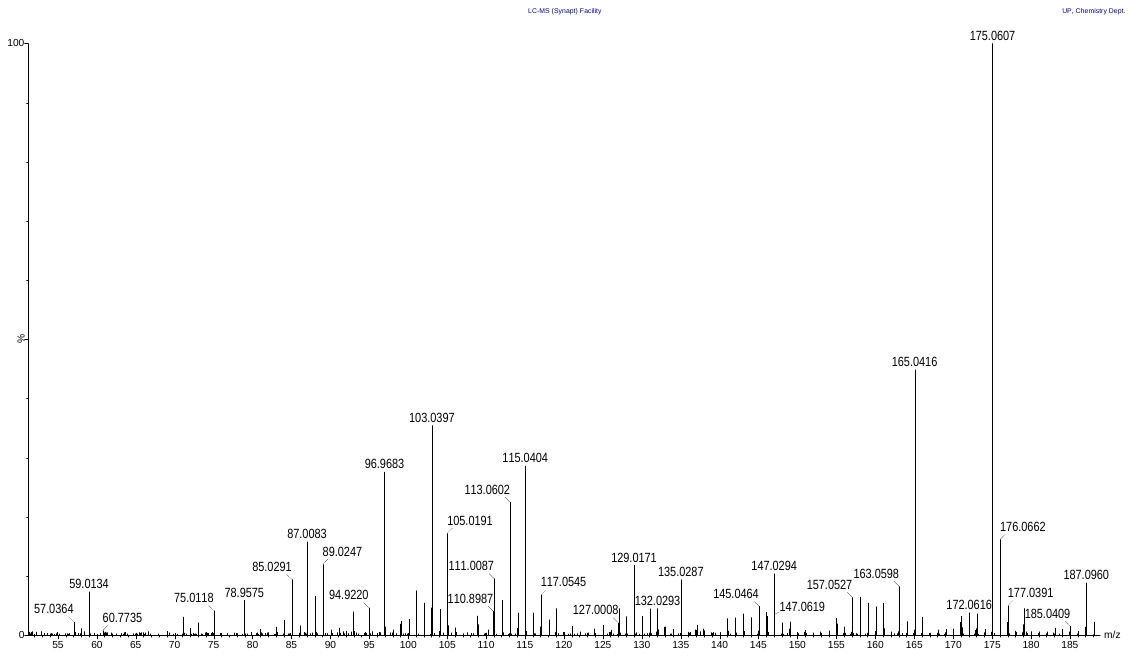

**Figure SI-11**: UPLC-QToF-MS results of streptol in *Psychotria kirkii* showing the presence of the ion fragments 85, 108, 111, 121 and 175 *m/z.*

**References**

Pinto-Carbó, M., Sieber, S., Dessein, S., Wicker, T., Verstraete, B., Gademann, K., Eberl, L., Carlier, A., 2016. Evidence of horizontal gene transfer between obligate leaf nodule symbionts. ISME Journal 10, 2092–2105. <https://doi.org/10.1038/ismej.2016.27>

Hsiao, C.C., Sieber, S., Georgiou, A., Bailly, A., Emmanouilidou, D., Carlier, A., Eberl, L., Gademann, K., 2019. Synthesis and biological evaluation of the novel growth inhibitor streptol glucoside, isolated from an obligate plant symbiont. Chemistry - A European Journal 25, 1722–1726. <https://doi.org/10.1002/chem.201805693>

Georgiou, A., Sieber, S., Hsiao, C.C., Grayfer, T., Gorenflos López, J.L., Gademann, K., Eberl, L., Bailly, A., 2021. Leaf nodule endosymbiotic *Burkholderia* confer targeted allelopathy to their *Psychotria* hosts. Scientific Reports 11, 1–15. <https://doi.org/10.1038/s41598-021-01867-2>
